# Nitrogen deletion in HDAC-targeting anticancer drug biosynthesis

**DOI:** 10.1101/2025.03.13.642940

**Authors:** Xinyun Jian, Jinlian Zhao, Edward Marschall, Kai-Chen Wu, Chuan Huang, Munro Passmore, Douglas M. Roberts, David P. Fairlie, Max J. Cryle, Lona M. Alkhalaf, Gregory L. Challis

## Abstract

Small molecules play indispensable roles in living systems as hormones, membrane bilayer constituents, enzyme cofactors, metal chelators, and defensive chemicals. The exceptional structural diversity of such molecules underpins their wide-ranging biological functions. Precise functional group insertion into various molecular scaffold classes is a hallmark of small molecule biosynthesis and is frequently important for biological activity. Functional group deletion is also important, but mechanisms are less well understood. Here, we report deletion of a cysteine-derived nitrogen atom during assembly of the conserved pharmacophore in the anticancer drug romidepsin, and related depsipeptide HDAC inhibitors. A shunt metabolite hydroxylated at the cysteine-α-carbon-derived position is a thousand-fold less active, indicating nitrogen deletion is important for potent HDAC inhibition. In vitro reconstitution and dissection of the complete nonribosomal peptide synthetase-polyketide synthase-mediated pathway for pharmacophore assembly reveal that cryptic *S*-octanoylation is catalysed by an atypical heterocyclisation domain, while multifunctional dehydratase and ketoreductase domains and *trans*-acting phosphotransferase and flavin-dependent oxidoreductase enzymes catalyse successive transformations in nitrogen deletion. Our findings significantly advance the understanding of heteroatom deletion mechanisms in small molecule biosynthesis and highlight the key role this can play in enhancing bioactivity. One-pot biocatalytic synthesis of the pharmacophore provides foundations for chemoenzymatic approaches to next-generation HDAC inhibitors.

## Main

Hydrocarbon functionalization via site-specific insertion of carbon, oxygen, nitrogen, sulfur, and halogen atoms is chemically challenging^1^, yet plays an important role in the biosynthesis of most classes of naturally occurring small molecule, including antibiotics, metallophores, cofactors, hormones, and bilayer constituents (**Fig. 1a-c**). Important examples include addition of methyl groups by methyl-cobalamin dependent radical-*S*-adenosylmethionine (SAM) enzymes and oxidative carbocyclizations by Rieske oxygenase-like enzymes^2–4^; insertion of hydroxyl groups and oxidative oxacyclizations by cytochromes P450 (CYPs) and non-heme iron-dependent oxygenases (NIDOs)^5–7;^ insertion of nitro groups and oxidative azacyclizations by CYPs^8,9^; insertion of thiol groups and oxidative thiacyclisations by radical-SAM enzymes and NIDOs^10–12;^ and aromatic and aliphatic halogenation by flavin and non-heme iron-dependent halogenases^13,14^. In addition, site-specific hydrocarbon hydroxylation and methylation plays important roles in modulating the function of proteins, such as hypoxia inducible factor-α and methyl-coenzyme M reductase^15,16^.

Functional group deletion also plays an important role in the assembly of several small molecule classes, for example carbon deletion in steroid hormone and glycopeptide antibiotic biosynthesis^17,18^, and oxygen deletion in fatty acid and polyketide antibiotic biosynthesis (**Fig. 1d, e**)^19,20^. However, such processes have been less widely investigated and, to the best of our knowledge, deletion of heteroatoms other than oxygen in small molecule biosynthesis remains hitherto unreported. Although nitrogen deletion is proposed to be involved in the biosynthesis of romidepsin and related depsiptide HDAC inhibitors (**Fig. 1f**) ^21^, this has yet to be experimentally validated.

HDACs are epigenetic regulators that play a vital role in chromatin remodelling, gene expression, and DNA repair by modulating the acetylation of histones (**Extended Data Fig. 1a**). Aberrantly high expression of HDACs has been reported for many different types of cancer cell^22^. Consequently, HDACs have emerged as promising targets for therapeutic intervention^23^. There are 18 mammalian HDAC isoforms, which are divided into four classes based on structural homology. Classes I, II and IV are zinc-dependent, whereas class III is NADH-dependent^23^. Three families of anti-cancer drugs that target zinc-dependent HDACs have been approved for clinical use: hydroxamic acids, benzamides, and depsipeptides (exemplified by romidpesin **1**) (**Extended Data Fig. 1 b & c**)^23^.

**Figure 1.**
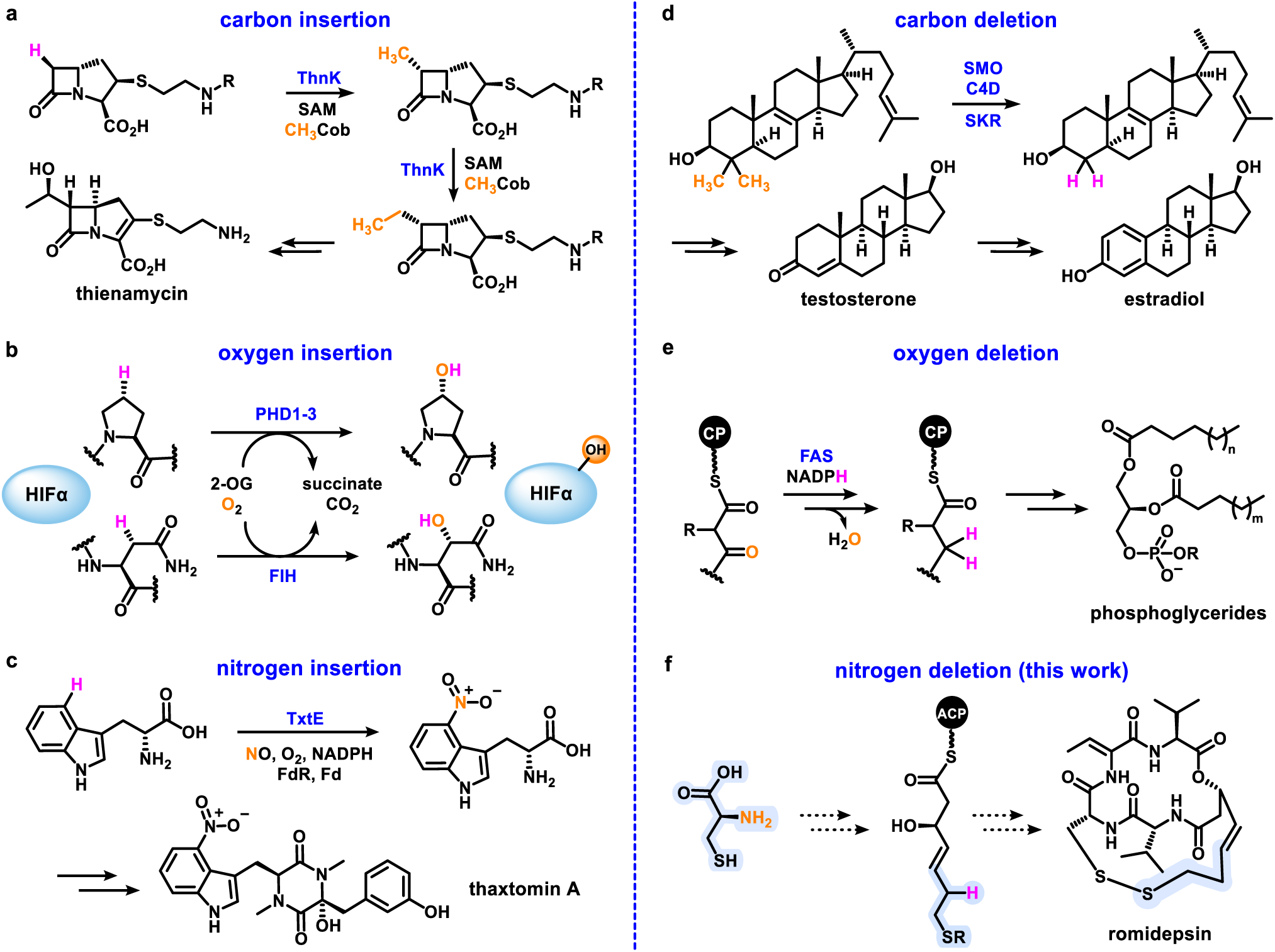
Examples of site-specific atom insertion and deletion in small molecule biosynthesis and protein post-translational modification. a. Methyl-cobalamin dependent radical-SAM enzyme ThnK catalyses insertion of two successive carbon atoms into a bicyclic β-lactam intermediate in the biosynthesis of the antibiotic thienamycin. b. prolyl hydroxylases (PHDs) and an asparaginyl hydroxylase (factor inhibiting HIF, (FIH)) catalyse oxygen insertion into specific proline and asparagine residues, respectively, of hypoxia inducible factor-α (HIFα). c. Cytochrome P450 TxtE catalyses nitrogen insertion (in the form of a nitro group) into L-tryptophan during the biosynthesis of the herbicide thaxtomin A. d. Sterol-4α-methyl-oxidase (SMO), 3β-hydroxysteroid dehydrogenases/C-4 decarboxylase (C4D) and sterone ketoreductase (SKR) catalyse two successive carbon deletion reactions to form zymosterol en route to the mammalian steroid hormones testosterone and estradiol. e. Ketoreductase (KR), dehydratase (DH), and enoylreductase (ER) enzymes catalyse oxygen deletion in the biosynthesis of fatty acids incorporated into membrane bilayer-forming phosphoglycerides. f. Proposed deletion of nitrogen from cysteine during the biosynthesis of the conserved pharmacophore incorporated into the anticancer drug romidepsin and related bicyclic HDAC inhibitors (investigated in this work). Atoms inserted or deleted are highlighted in orange and the corresponding H atoms removed or added, respectively, are highlighted in pink.

While benzamides and hydroxamic acids are synthetic drugs, depsipeptides are specialised metabolites produced by diverse bacterial genera containing a conserved (*3S*, *4E*)-3-hydroxy-7-mercapto-4-heptenoic acid pharmacophore fused to a variable peptidyl cap (**Extended Data Fig. 1c**)^21–24^. Variations in the composition of the peptidyl cap have the potential to alter HDAC isoform selectivity. The pharmacophore thiol group is masked via either disulfide formation with a Cys residue in the peptidyl cap or octanoylation (**Extended Data Fig. 1c**). Intracellular disulfide reduction (in the case of romidepsin, burkholdacs, spiruchostain, and FR901375) and thioester hydrolysis (in the case of largazole) unmasks the thiol group enabling coordination to the catalytically essential Zn^2+^ ion in the active site of HDACs. (**Fig. 2 a & b**)^25–27^. The narrow hydrophobic channel leading to the active site Zn^2+^ ion in class I HDACs imposes significant structural constraints on the linker between the Zn^2+^-binding thiol group and the peptidyl cap^28^. Functionalisation of this linear hydrophobic moiety, e.g. by introduction of a heteroatom, can impact HDAC inhibition, explaining why it may be necessary to delete the Cys-derived nitrogen during pharmacophore assembly.

**Figure 2.**
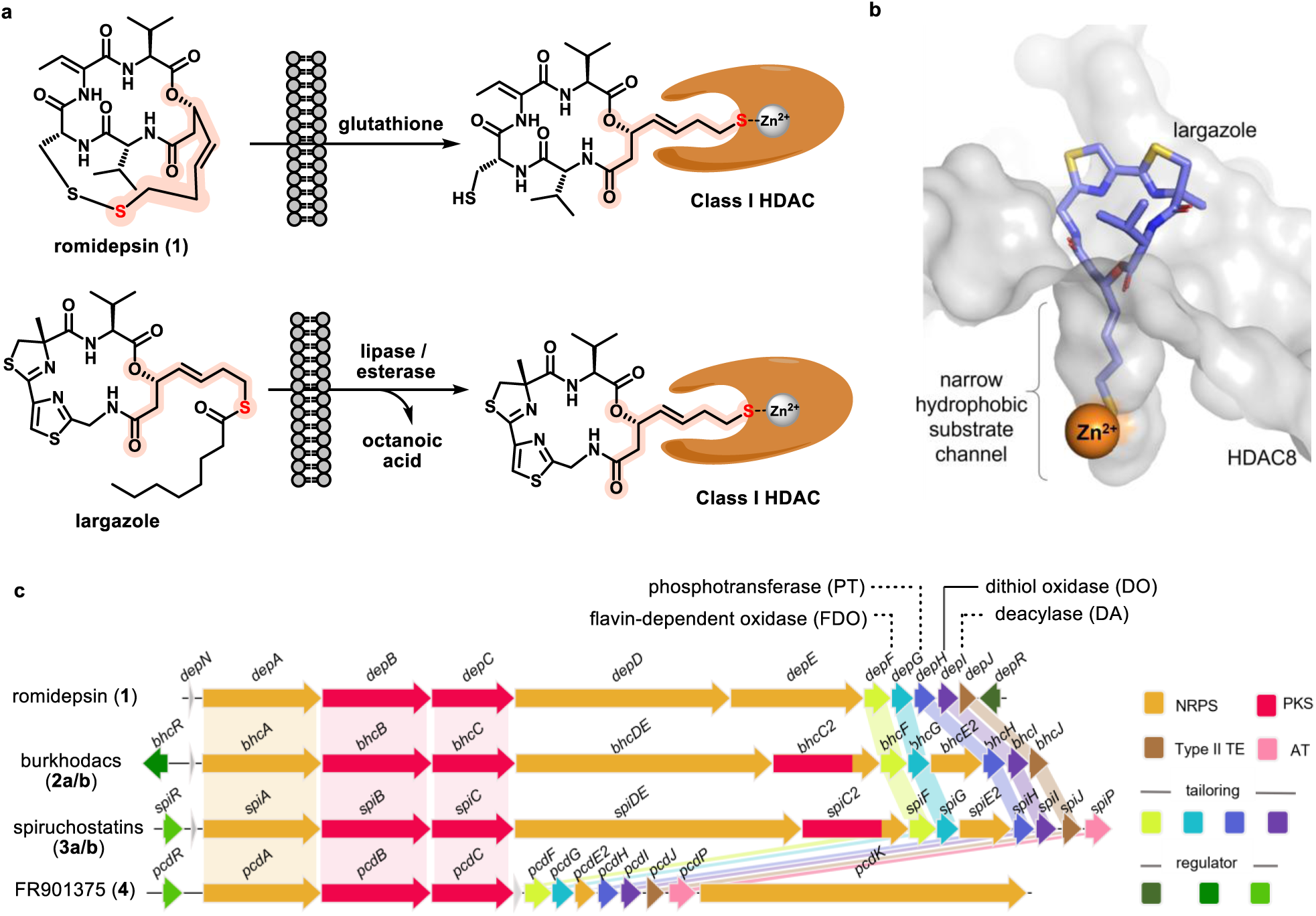
Mechanism of action and biosynthetic gene clusters for romidepsin and related HDAC-targeting depsipeptides. **a**, Mechanism of action of depsipeptide HDAC inhibitors. Either intracellular reduction of the disulfide (top, in most cases) or thioester hydrolysis (bottom, in the case of largazole) unmasks the terminal thiol (in red) of the conserved pharmacophore (highlighted in orange). This thiol binds the Zn^2+^ ion in the HDAC catalytic pocket. **b**, X-ray crystal structure of largazole-HDAC8 complex (PDB ID: 3RQD) showing the peptidyl cap interacts with the rim of the pocket, while the pharmacophore extends into the narrow, hydrophobic substrate channel^25^. **c**, Comparison of the romidepsin (*dep*), burkholdac (*bhc*), spiruchostatin (*spi*) and FR901375 (*pcd*) biosynthetic gene clusters. Highly conserved genes encode an NRPS-PKS (*A*, *B*, *C*) and four tailoring enzymes (*F*, *G*, *H*, *I*). DepH has been shown to catalyse the last step in romidepsin biosynthesis – disulfide formation. Variable genes encoding NRPSs (*D*, *E*, *DE*, *E2*, *K*) and an a PKS-NRPS (*C2*) are proposed to catalyse peptidyl cap assembly.

The biosynthetic gene clusters (BGCs) for romidepsin **1**, burkholdacs A and B **2a**/**b**, and spiruchostatins A and B **3a**/**b**, were identified and sequenced more than a decade ago (**Fig. 2c**)^21,29–31^. These three HDAC inhibitors have similar structures with a cysteine residue at the second position in the peptidyl cap that forms a disulfide linkage with the terminal thiol group of the conserved pharmacophore (**Extended Data Fig. 1c**). The corresponding BGCs have similar architectures, with *depA*/*bhcA*/*spiA*, *depB*/*bhcB*/*spiB*, and *depC*/*bhcC*/*spiC* encoding a highly conserved hybrid nonribosomal peptide synthetase (NRPS)-modular polyketide synthase (PKS) that is hypothesised to play a key role in building the pharmacophore^32^. An NRPS encoded by *depD*/*depE* and very similar NRPS-PKSs encoded by *spiDE*/*bhcDE*, and *spiE2/bhcE2* are proposed to assemble the peptidyl caps of romidepsin and the burkholdacs/spiruchostatins, respectively (**Extended Data Fig. 2**). The final step in romidpesin biosynthesis is catalysed by the flavin-dependent dithiol oxidase encode by *depH*, resulting in disulfide formation between the cysteine residue in the peptidyl cap and the terminal thiol of the pharmacophore (**Extended Data Fig. 2**)^33,34^.

In 2019, we identified previously overlooked β-hairpin docking (β-HD) domains appended to the N-termini of the first subunits (DepD/BhcDE/SpiDE) of the machinery for variable peptidyl cap assembly in romidespin, burkholdac, and spircuhostatin biosynthesis (**Extended Data Fig. 2**) ^35,36^. These were hypothesised to interact with similarly overlooked short linear motifs (SLiMs) attached to the C-termini of the final subunits (DepC/BhcC/SpiC) of the pharmacophore biosynthetic apparatus, enabling it to engage with diverse peptidyl cap biosynthetic machineries (**Extended Data Fig. 2**). We recently exploited these findings to identify the FR901375 BGC in the genome sequence of *Pseudomonas chlororaphis* subsp. *piscium* DSM 21509^37^. This revealed that the final subunit of the pharmacophore biosynthetic apparatus (PcdC) also has a SLiM fused to its C-terminus, which was proposed to interact with a β-HD domain appended to the N-terminus of PcdK, the NRPS responsible for assembly of the peptidyl cap in FR901375 (**Extended Data Fig. 2**).

FR901375 has a peptidyl cap structurally distinct from romidepsin and the burkholdacs / spiruchostatins. The cysteine residue linked via a disulfide to the pharmacophore is in the third position rather than the second (**Extended Data Fig. 1c**). Intriguingly, the organisation of the FR901375 BGC deviates markedly from the romisepsin, burkholdac, and spircuhostatin BGCs (**Fig 2c**). The FR901375 BGC appears to have evolved from the spiruchostatin BGC by acquisition of *pcdK* downstream of the *pcdF-pcdP* gene cassette, followed by deletions in *pcdDE*/*pcdC2*, and *pcdE2*, encoding the assembly line for the original tripeptidyl caps of the spiruchostatins (**Fig. 2c**)^37^. Notably, four genes within the *pcdF-pcdP* cassette (*pcdF*, *pcdG*, *pcdI* and *pcdJ*), in addition to *pcdH*, encoding a homologue of the disulfide-forming enzyme DepH, are conserved across all four known depsipetide HDAC inhibitor BGCs.

Here we use a combination of bioinformatics, molecular genetics, structure elucidation of metabolites accumulated in gene deletion mutants, *in vitro* reconstitution, intact protein mass spectrometry, synthetic analogues of key precursors, and cysteamine-mediated / spontaneous offloading of PKS-bound biosynthetic intermediates to elucidate the pathway for assembly of the conserved pharmacophore in romidepsin and related depsipeptide HDAC inhibitors. Our work reveals that the pharmacophore terminal thiol is protected as a thioester throughout metabolite assembly, suggesting a unified mechanism for the biosynthesis of largazole and disulfide-containing members of the depsipeptide HDAC inhibitor family. Moreover, it shows that the Cys-derived nitrogen is deleted from a defined PKS-bound intermediate in pharmacophore assembly. This multistep process involves a combination of unusual multifunctional PKS domains and unique *trans*-acting ATP and flavin-dependent enzymes. A three-orders-of-magnitude reduction in activity for a FR901375 analogue bearing a hydroxyl group in place of the deleted nitrogen indicates the importance of removing heteroatoms from this position for potent HDAC inhibition. Finally, we exploit our findings to produce the *S*-protected pharmacophore in a single reaction vessel from three precursors via seventeen successive chemical transformations catalysed by eight purified recombinant enzymes employing five cofactors.

To the best of our knowledge, this is the first example of nitrogen deletion in small molecule biosynthesis. Delineating each step in this novel process not only expands our understanding of how Nature achieves chemically challenging structural modifications in small molecule assembly, highlighting how molecular scaffolds are selectively sculpted to optimise bioactivity, but also lays the foundations for greener and more sustainable chemoenzymatic approaches to the synthesis of novel depsipeptide HDAC inhibitors.

### Bioinformatics analysis of putative pharmacophore biosynthetic machinery

We were puzzled by previous suggestions that the chain-initiating acyl-CoA ligase (AL) domain, and the condensation (C) in module 1, and the dehydratase (DH) domains in modules 2 and 3 of the NRPS-PKS hypothesised to play a key role in pharmacophore assembly are catalytically inactive, despite being conserved in all homologues encoded by depsipeptide HDAC inhibitor BGCs identified to date ^21,29–31^. Thus, we conducted a careful manual reannotation of these NRPS-PKSs (**Extended Data Fig. 3**). This revealed that the AL and C domains contain all the conserved amino acids necessary for catalysis. Furthermore, a previously overlooked carrier protein (CP) domain between the AL and C domains was identified. These features were conserved in the corresponding NRPS-PKSs involved in romidepsin, burkholdac, and spiruchostatin biosynthesis (**Extended Data Fig. 3a**). Phylogenetic analysis of the module 1 C domains showed they group in a distinct clade closely related to heterocyclisation (Cy) domains, which typically catalyse *N*-acylation of Cys, Thr, or Ser residues loaded onto the downstream CP domain by an adenylation (A) domain, followed by cyclodehydration to form the corresponding thiazoline / oxazoline (**Extended Data Fig. 3b**). Sequence alignment of these C domains and canonical Cy domains showed variations in the conserved DX_5_D motif and other residues known to be catalytically important (**Extended Data Fig. 3c**) ^38^. Together, these findings led us to hypothesise that the likely functional AL domain and the newly discovered CP domain together catalyse chain initiation via incorporation of an unknown starter unit, which the module 1 C domain then utilizes to *S*-acylate the L-Cys residue loaded onto the downstream CP domain (**Figure 3a**).

Similarly, sequence analyses identified an intact His-Asp catalytic dyad in the module 2 DH domains of the NRPS-PKSs suggesting they are functional (**Extended Data Fig. 3a**). Phylogenetic analysis of these DH domains showed they group in a single clade, closely related to the bifunctional dehydratase-enoyl-isomerase (DH^ei^) domain GphDH_1_ and putative diene synthase (DH^d^) domains in several *cis*-AT PKSs (**Extended Data Fig. 3d**) ^39,40^. Previous studies have identified a characteristic substitution of proline in the HX_8_P motif in enoyl-isomerizing DH domains (DH^i^ and DH^ei^) ^40^. This substitution was also found in the HX_8_P motif of all the module 2 DH domains, suggesting they may catalyse Δ2 to Δ3 double bond isomerisation, in addition to canonical β-hydroxy thioester dehydration, during pharmacophore biosynthesis (**Extended Data Figure 3e**).

### Genetic investigation of roles played by putative tailoring enzymes

Four genes in the FR901375 BGC encoding putative tailoring enzymes (*pcdF*, *pcdG*, *pcdH*, and *pcdI*) are universally conserved across all four known depsipeptide HDAC inhibitor BGCs. The function of *pcdH/bhcH/SpiH* can be confidently assigned as formation of the disulfide common to FR901375, the spiruchostatins, the burkholdacs, and romidepsin, based on previous biochemical studies of DepH (**Extended Data** Fig. 2)^33,34^. We hypothesised that *pcdF*, *pcdG*, and *pcdI*, which encode a putative flavin-dependent oxidoreductase (FDO), phosphotransferease (PT), and deacylase (DA), respectively, and their homologues in the other BGCs likely play a key role in formation of the only other structurally conserved feature of all the metabolites – the pharmacophore (**Figure 3a**).

To investigate the roles played by *pcdF*, *pcdG,* and *pcdI*, in-frame deletions were created in each of these genes in *P. chlororaphis* subsp. *piscium* DSM 21509. UHPLC-ESI-Q-TOF-MS analyses of organic extracts from the *pcdF* and *pcdG* mutants showed the production of FR901375 (**4**) was abolished. Interestingly, the *pcdG* mutant accumulated shunt metabolite **6**, which was purified and shown by high resolution MS and NMR spectroscopy to be an analogue of **4** missing a double bond but containing an additional hydroxyl group in the pharmacophore (**Fig. 3b**). The same metabolite profile was observed in a *pcd*F/*pcd*G double mutant (**Fig. 3a**), indicating that PcdF acts after PcdG in FR901375 biosynthesis.

UHPLC-ESI-Q-TOF-MS analysis of organic extracts from the *pcdI* mutant showed that, in addition to FR901375 (**4**), a new metabolite **7** had accumulated. This was purified and shown by high resolution MS and NMR spectroscopy to be an analogue of **4** in which the disulfide is reduced, and the pharmacophore thiol is octanoylated (**Fig. 3a**). This suggests that the AL domain loads an octanoyl residue onto the downstream CP domain, which gets transferred onto the thiol of the cysteine residue attached to the CP domain in module 1 by the C domain. Thus, PcdI appears to hydrolytically cleave the octanoyl thioester as the penultimate step in FR901375 biosynthesis, which unmasks the pharmacophore thiol for subsequent PcdH-catalysed disulfide formation.

Mutants of the burkholdac and romidpesin producers with in-frame deletions in the *pcdG* and *pcdI* homologues accumulated analogous shunt metabolites / biosynthetic intermediates (**Extended Data Fig. 4**), confirming that the tailoring enzymes these genes encode have a conserved role in pharmacophore biosynthesis.

**Figure 3.**
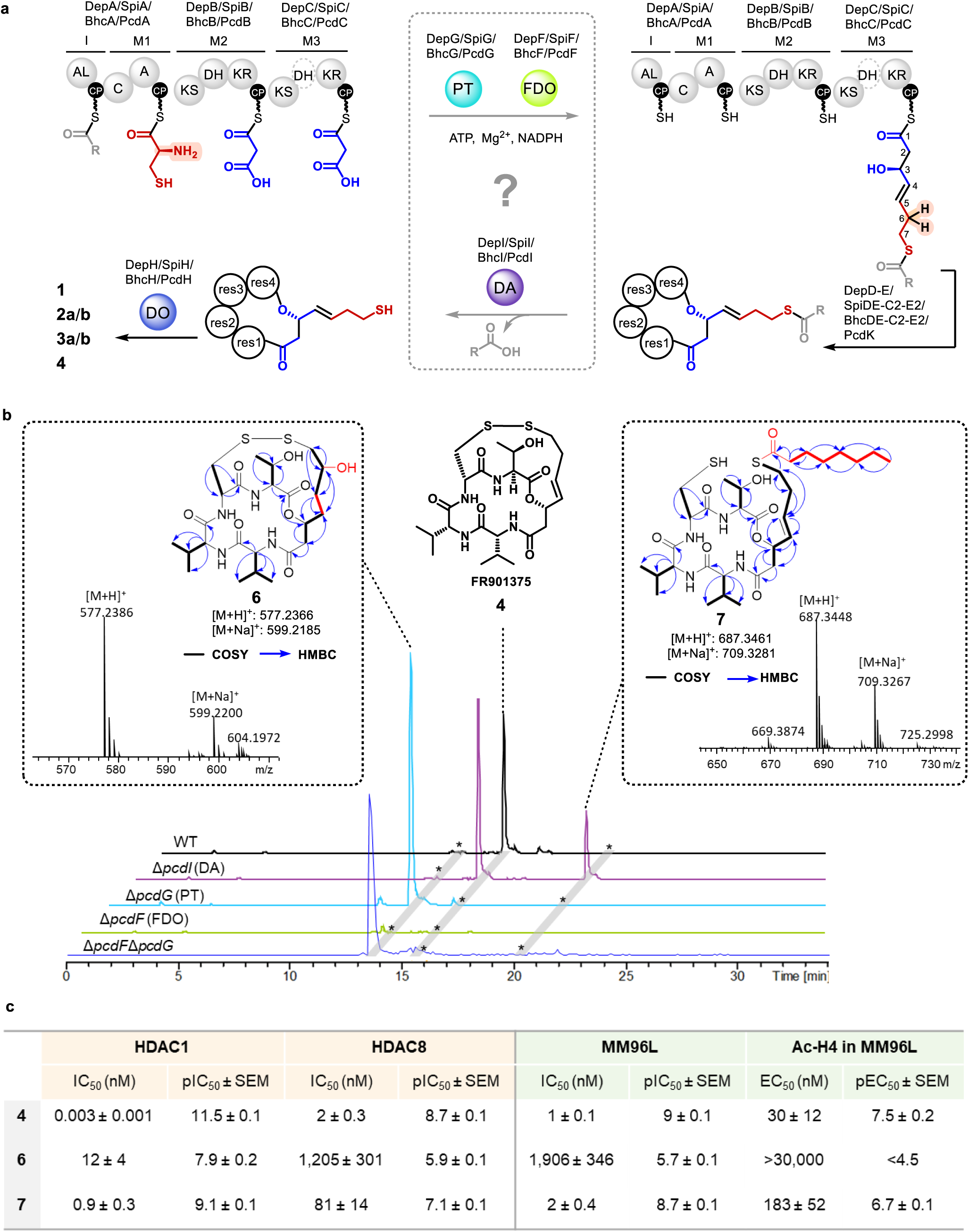
Genetic characterization of the role played by conserved tailoring enzymes in FR901375 biosynthesis and biological activity of FR901375 analogues with modifications to the pharmacophore. **a**, Proposed pathway for assembly of **1**, **2a/b**, **3a/b** and **4** based on reannotation of the NRPS-PKS encoded by the *A*-*C* genes and the universal conservation of the *F*-*I* genes. Predicted building blocks for the pharmacophore are shown bound to the carrier protein (CP) domains in the chain initiation module (labelled I) and each chain elongation module (labelled M1-3). Three conserved tailoring enzymes, encoded by the *F*, *G* and *I* genes, are hypothesised to be involved in pharmacophore biosynthesis, two of which (the phosphotransferase (PT) and flavin-dependent oxidoreductase (FDO)) are proposed to be directly involved in deletion of the Cys-derived amino group during pharmacophore assembly. Variable peptidyl caps are incorporated by diverse NRPS and NRPS-PKS assembly lines encoded by the *D*, *E*, *DE*, *C2*, *E2* and *K* genes (see **Extended Data** Fig.2 for further details). The octanoyl thioester is hydrolysed by the deacetylase (DA) encoded by the *I* gene and the disulfide is formed by the dithiol oxidase (DO) encoded by the *H* gene **b,** UHPLC-ESI-Q-TOF-MS and NMR spectroscopic analysis of shunt metabolite **6** and biosynthetic intermediate **7** produced by the *pcdG* and *pcdI* mutants of *P. chlororaphis* subsp. *piscium* DSM21509. Extracted ion chromatograms at *m/z* = 559.2260 ± 0.002, 577.2366 ± 0.002 and 687.3448 ± 0.002 (corresponding to [M+H]^+^ for **4**, **6**, and **7**, respectively) show that FR901375 (**4**) production is abolished in the *pcdF*, *pcdG*, and *pcdG*/*pcdF* mutants, shunt metabolite **6** accumulates in the *pcdG*, and *pcdG*/*pcdF* mutants, and biosynthetic intermediate **7** is produced alongside **4** by the *pcdI* mutant. Correlations observed in COSY and HMBC NMR spectra are indicated by bold lines and arrows, respectively. Structural changes to the pharmacophore in **6** and **7** are highlighted in red. **c**. Comparison of HDAC1 and HDAC8 inhibition (orange columns), and cytoselectivity / accumulation of acetylated histone H4 in MM96L melanoma cells (green columns) in the presence of **4**, **6,** or **7**.

### Biological activity of FR901375 analogues with modifications to the pharmacophore

The production of shunt metabolite **6** and biosynthetic intermediate **7** in the *pcdG* and *pcdI* mutants, respectively, provided us with an opportunity to assess the impact of structural changes to the conserved pharmacophore on biological activity. Thus, we compared **4**, **6** and **7 for** *in vitro* inhibition of HDAC1 and HDAC8 catalytic activity , in addition to the cytotoxicity of these compounds against MM96L melanoma cells and the accumulation of acetylated histone H4 in these cells using well-established procedures^41^. While inhibition of HDAC1 and HDAC8 activity for the *S*-octanoylated derivative **7** and the hydroxylated derivative **6** was approximately two and three orders of magnitude less than FR901375 (**4**), respectively, **4** and **7** had similar activity against MM96L cells, whereas **6** was three orders of magnitude less active (**Fig. 3c, Extended data Fig.5**). These data indicate that, as for largazole, the pharmacophore thiol of **7** can be unmasked via hydrolysis of the octanoyl thioester *in vivo*. On the other hand, they show that addition of a heteroatom at C6 of the pharmacophore has a very detrimental effect on biological activity, explaining why Nature has evolved a process for the deletion of the Cys-derived amino group during pharmacophore assembly.

### *In vitro* reconstitution of L-cysteine loading and S-acylation

To verify the proposed function of the AL domain and the module 1 C domain, we reconstituted the assembly of CP-bound *S*-octanoyl-L-Cys *in vitro* using purified recombinant proteins overproduced in *E. coli* (**Extended Data Fig. 6**). Owing to the insolubility of proteins encoded by FR901375 BGC, we used homologues from the romidepsin system.

The AL-CP didomain and C-A-CP tridomain were converted to their respective *holo* forms (Reaction 1, **Fig. 4a**; Reaction 3**, Fig.4b**) using the promiscuous phosphopantetheinyl transferase Sfp.^42^ Following incubation of the *holo*-AL-CP didomain with octanoic acid and ATP (Reaction 2, **Fig. 4a**), UHPLC-ESI-QTOF-ESI-MS analysis of the intact protein showed a mass shift consistent with attachment of an octanoyl residue to the CP domain. Similarly, an L-Cys residue was loaded onto the tridomain (Reaction 4, **Fig. 4b**). Incubation of the octanoylated didomain with the cysteinylated tridomain resulted in a mass shift consistent with formation of an *S*-octanoyl-cysteinyl moiety attached to the CP domain in the latter (Reaction 5, **Fig. 4b**).

**Figure 4.**
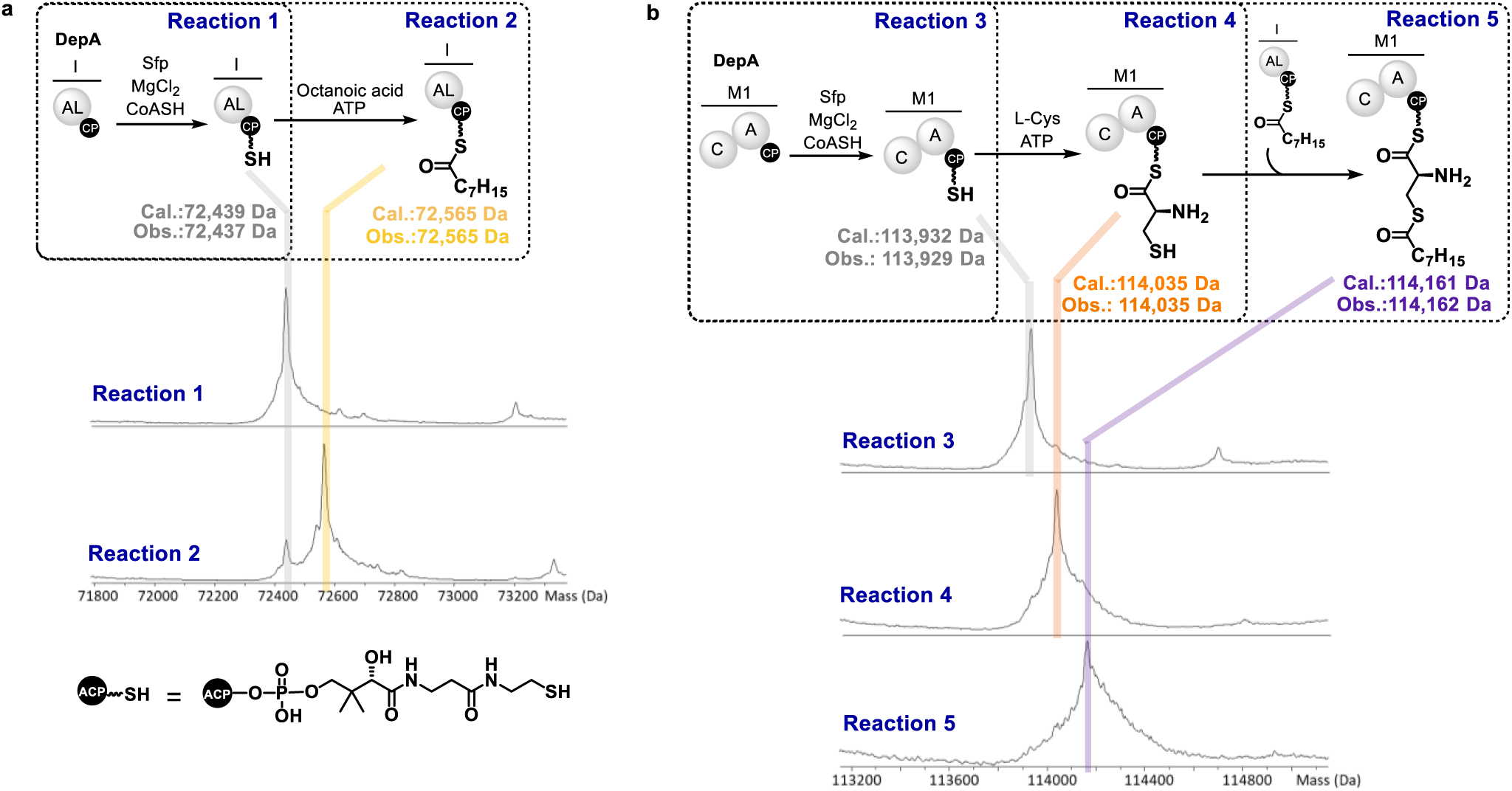
Intact protein mass spectrometric analysis of early stages in pharmacophore biosynthesis. Schematic representations of each reaction are shown at the top of the panels and deconvoluted mass spectra from UHPLC-ESI-Q-ToF-MS analyses of each reaction are shown below. **a**, *holo* modification (Reaction 1) and subsequent octanoylation (Reaction 2) of the chain-initiating AL-CP didomain of DepA. **b**, *apo* to *holo* modification (Reaction 3), cysteinylation (Reaction 4), and S-acylation (Reaction 5) of the DepA C-A-CP tridomain (M1).

### *In vitro* reconstitution of nitrogen deletion from S-acylated L-cysteine

We next investigated the reactions catalysed by purified recombinant DepB, DepG, and DepF (**Extended Data Fig. 6**). No change to the *S*-octanoyl-cysteinyl moiety assembled in reaction 5 was observed when DepG and/or DepF plus relevant cofactors were included. We thus surmised that nitrogen deletion occurs at a later point in pharmacophore assembly and turned our attention to reconstitution of the reactions catalysed by the second chain elongation module DepB.

The CP domain of DepB was malonylated using Sfp (Reaction 6, **Fig. 5**) and incubated with *S*-octanoyl-cysteinylated module 1 (*vide supra*) and NADPH (Reaction 7, **Fig. 5**). UHPLC-ESI-QTOF-ESI-MS analysis of intact DepB revealed complete consumption of the malonyl group and formation of three new species with molecular weights of 165,320, 165,109, and 165,062 Da. The first of these is consistent with the presence of the intermediate resulting from one round of canonical chain elongation, β-ketoreduction, and α, β-dehydration on the CP domain, in addition to attachment of a *S*-octanoyl-cysteinyl moiety (donated by the module 1 CP domain) to the active site Cys residue of the KS domain. The second species is consistent with attachment of the intermediate resulting from one round of chain elongation and β-ketoreduction but no α, β-dehydration to the CP domain, and the third species is consistent with loss of this intermediate from the CP domain via spontaneous γ-lactamisation, followed by repriming of the KS domain with the *S*-octanoyl-cysteinyl moiety from the module 1 CP domain (or *vice versa*).

Addition of DepG and ATP to the mixture (Reaction 8, **Fig. 5**) enabled detection of a fourth species in the UHPLC-ESI-QTOF-ESI-MS analyses of intact DepB with a molecular weight of 165,402 Da. This corresponds to a CP-bound intermediate in which the allylic amine has been replaced with a saturated alcohol that has undergone phosphorylation (**Fig. 5**). This (i) is consistent with the proposal that DepG functions as phosphotransferase; (ii) suggests that the substrate of DepG is the γ-hydroxy thioester, consistent with the structure of FR901375 analogue **6** accumulated in the *depG* mutant of *P. chlororaphis* subsp. *piscium*; (iii) indicates that the γ-hydroxy thioester arises from non-canonical DH domain-catalysed isomerisation of the allylic amine to an imine, followed by hydrolysis, and reduction of the resulting γ-keto group by the KR domain; and (iv) suggests the γ-hydroxy thioester is offloaded from the DepB CP domain via spontaneous lactonization, explaining why it is not observed in Reaction 7 (**Fig. 5**). The species with molecular weight corresponding to the CP-bound phosphorylated alcohol was consumed upon addition of DepF and two new species with molecular weights of 165,305 Da and 165,319 Da emerged (Reaction 9, **Fig. 5**). These are consistent with a CP-bound α, β-enoyl thioester, arising from elimination of phosphate and isomerisation of the resulting β, γ-thioester, and a CP-bound β-hydroxy-thioester arising from DH domain-catalysed hydration of the α, β-enoyl thioester (as observed in other systems, due to the reversibility of the dehydration reaction^39^), respectively.

**Figure 5.**
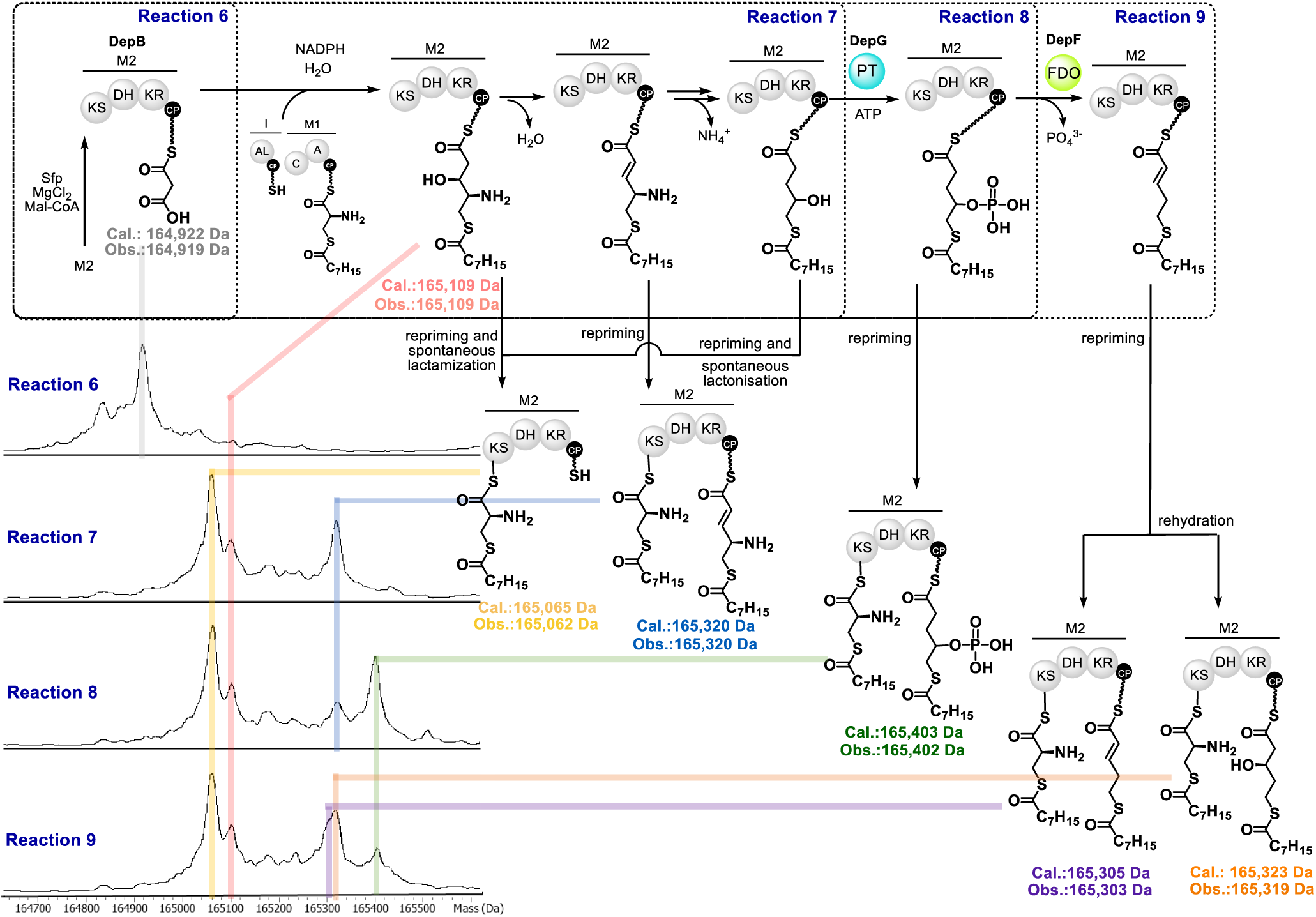
Intact protein mass spectrometric analysis of intermediate steps in pharmacophore biosynthesis. Schematic representations of each reaction are shown at the top of the panels and deconvoluted mass spectra from UHPLC-ESI-Q-ToF-MS analyses of each reaction are shown below. Malonylation (Reaction 6), canonical chain elongation, β-ketoreduction, and α, β-dehydration, followed by non-canonical double bond migration, imine hydrolysis, and γ-ketoreduction (Reaction 7), DepG-catalysed phosphorylation (Reaction 8), and DepF-catalysed conversion of the γ-phosphoryloxy thioester to the α, β-enoyl thioester (Reaction 9) in DepB (M2).

To verify the hypothesis that both the γ-amino-β-hydroxy, and γ-hydroxy thioester intermediates can be offloaded from DepB via spontaneous cyclisation, we analysed organic extracts of Reaction 7 using UHPLC-ESI-Q-ToF-MS. This identified species with molecular formulae corresponding to the expected lactam **13** and lactone **14** products (**Extended Data Fig. 7**). In addition, we observed a species with molecular formula corresponding to lactam **12 when NADPH was omitted**, which appears to arise from spontaneous cyclisation of the γ-amino-β-keto thioester intermediate (**Extended Data Fig. 7**). No species with a molecular formula corresponding to the lactam resulting from offloading of the γ-amino-α, β-enoyl thioester intermediate was observed, consistent with the *E*-configured enone in this intermediate preventing spontaneous cyclisation. When Reaction 7 was conducted in a 1:1 mixture of ^18^O-labelled and unlabelled buffered water, approximately 50% ^18^O-labelling of lactone **14** was observed (**Extended Data Fig. 7**). This is consistent with hydrolysis of a γ-imine to form the corresponding keto group, which gets reduced to a hydroxyl group.

To verify that the DepB KR domain, DepG, and DepF, catalyse non-canonical γ-ketoreduction, phosphorylation, and formal phosphate elimination-double bond isomerisation, respectively, we synthesised the CoA-thioester of a stabilised analogue **15** of the putative γ-keto thioester intermediate and loaded it onto the DepB CP domain (Reaction 10; **Extended Data Fig. 8**). CP-bound products formed after addition NADPH alone, NADPH and DepG/ATP, or NADPH, DepG/ATP and DepF were offloaded with cysteamine^43^ and UHPLC-ESI-QToF-MS analyses were used to identify the cysteamine adducts in organic extracts of each reaction. Compound **16** with molecular formula corresponding to the cysteamine offloaded adduct of the γ-hydroxy thioester was formed when just NAPDH was added (Reaction 11; **Extended Data Fig. 8**). This intermediate is not observed in the reaction with DepB replaced by a Y328F mutant containing a catalytically inactive KR domain. Addition of NADPH, DepG and ATP resulted in formation of a compound with a molecular formula corresponding to the cysteamine offloading adduct of the phosphorylated γ-hydroxy thioester **17** (Reaction 12; **Extended Data Fig. 8**). When DepF was also included in the mixture two distinct cysteamine adducts were formed (Reaction 13; **Extended Data Fig. 8**). One of these has a molecular formula corresponding to adduct **18** resulting from offloading plus conjugate addition of cysteamine to the α, β-enoyl thioester, whereas the other has a molecular formula corresponding to adduct **19** resulting from offloading of the β-hydroxy thioester (formed by DH domain-catalysed rehydration of the α, β-enoyl thioester – as seen in other systems^39^). Neither compound was formed when DepF was substituted with a catalytically inactive mutant in which the conserved active site aspartate residue is replaced by alanine (D367A).

In addition, we observed the formation of two compounds with molecular formulae corresponding to **20** and **21**, the cysteamine cleavage adduct of the γ-keto thioester and the product of spontaneous cyclisation of the γ-keto thioester, respectively, in all reactions, and a compound with a molecular formula corresponding to **22**, resulting from spontaneous cyclisation of the CP-bound γ-hydroxy thioester, in all reactions expect those from which NAPDH had been omitted or a DepB Y328F mutant, was employed (**Extended Data Fig. 8**). The structure of lactone **22** was confirmed by UHPLC-MS comparison with a synthetic standard (**Extended Data Fig. 8**).

### Assembly of the *S*-acylated pharmacophore in a single reaction vessel

Having established the timing of nitrogen deletion and key intermediates in the process using single turnover assays, we turned our attention to complete *in vitro* reconstitution of pharmacophore assembly via inclusion of purified recombinant DepA and DepC (Extended Data Fig. 4). To enable multiple turnovers we employed FabD1 to supply malonyl extender units to DepB and DepC^32^, and the type II thioesterase encoded *pcdJ* in the FR901375 BGC to release the product from DepC (**Extended Data Fig. 9a**). The eight purified recombinant enzymes needed for pharmacophore biosynthesis were incubated with octanoic acid, L-cysteine and malonyl-CoA in the presence of MgCl_2_, ATP, CoA, and NADPH at room temperature for 3 h. UHPLC-ESI-Q-ToF-MS analysis of an organic extract revealed a prominent peak at 21.7 min in the base peak chromatogram, with *m/z* corresponding to [M+H]^-^ for the *S*-protected pharmacophore **25 (Extended Data Fig. 9b)**. This result confirms that DepC catalyses the final round of chain elongation and β-ketoreduction in S-octanoylated pharmacophore assembly. Although some shunt products and intermediates were detected in the extract, indicating the need for further optimisation, the S-octanoylated pharmacophore **25** was clearly the major product of the reactions.

### Proposed pathway for pharmacophore assembly

Together, the above results delineate a remarkable pathway for conserved pharmacophore assembly in romidepsin biosynthesis (**Fig. 6**). This commences with the loading of octanoyl and L-cysteinyl residues onto the first and second CP domains of DepA by the upstream AL and A domains, respectively. The DepA C domain then catalyses transfer of the octanoyl group onto the thiol of the cysteinyl residue. FabD1, from fatty acid biosynthesis, functions as *trans*-acting AT to load a malonyl group onto the CP domain of DepB^32^. The KS domain of DepB accepts the *S*-octanoyl-cysteinyl moiety from the C-terminal CP domain of DepA onto its active site Cys residue and catalyses two-carbon chain elongation using the CP-bound malonyl group. The KR and DH domains of DepB then catalyse canonical β-ketoreduction and α, β-dehydration reactions.

The resulting allylic amine is converted to the corresponding γ-keto thioester via enamine and imine intermediates. Conversion of the allylic amine to the imine is likely catalysed by the DepB DH domain, because DH domains in other systems are known to catalyse similar double bond migration reactions. However, further experiments are needed to verify this. Whether the DH domain catalyses hydrolysis of the imine also remains to be experimentally investigated. Reduction of the γ-ketone to the corresponding alcohol is catalysed by the DepB KR domain. Bifunctional PKS KR domains are rare, but there is precedent for them in other systems, such as the *trans*-AT PKS responsible for bacillaene assembly^44^.

At this juncture, DepG catalyses in *trans* phosphorylation of the γ-hydroxy group, improving its leaving group ability. DepF also acts in *trans* to convert the resulting γ-phosphoryloxy thioester to the corresponding α, β-enone. While it might be tempting to suggest that this proceeds via β, γ-elimination of phosphoric acid, followed by β, γ to α, β double bond isomerisation, it unlikely that an enzymatic base would be capable of deprotonating the β-position of a γ-phosphoryloxy thioester, due to the very low acidity of the β-protons. DepF shows sequence similarity to acyl-CoA dehydrogenases (ACADs), flavin-dependent enzymes that typically catalyse conversion of saturated thioesters to their α, β-unsaturated counterparts. ACADs employ an active site Glu residue to deprotonate the α-carbon of their substrate concomitant with transfer of hydride from the β-carbon to the bound flavin cofactor. DepF has an Asp residue in place of the active Glu residue of canonical ACADs, suggesting the reaction it catalyses may proceed via initial α-carbon deprotonation. Several pathways to the product can be envisaged from the resulting nascent enolate, most of which involve participation of the bound flavin as a transient redox-active cofactor. Further studies will be required to establish (i) whether the flavin cofactor of DepF participates in catalysis or plays a purely structural role, as has been proposed for some other ACAD homologues^45^, and (ii) the full mechanistic details of the DepF-catalysed reaction.

The final steps in pharmacophore biosynthesis are catalysed by DepC, which also relies on FabA1 to catalyse transfer of a malonyl group onto its CP domain in *trans*. The DepC KS and KR domains then catalyse canonical chain elongation and ketoreduction reactions on the α, β-enoyl thioester intermediate attached to the DepB CP domain. Following NRPS-mediated fusion of the tetrapeptidyl cap onto the fully assembled *S*-octanoylated pharmacophore and chain release via macrolactonisation, we propose that DepI unmasks the terminal thiol group enabling DepH to catalyse disulfide formation with the thiol group of the cysteine residue embedded in the tetrapeptidyl cap (**Fig. 6**). Accumulation of a romidepsin derivative bearing an octanoyl group on the terminal thiol of the pharmacophore in a *depI* mutant (**Extended Data Fig. 4c**) is consistent with this hypothesis. However, this mutant still produces significant quantities of romidepsin, suggesting other enzymes encoded by genes outside the *dep* gene cluster can also catalyse partial unmasking of the pharmacophore thiol. The universal conservation of *A*, *B*, *C*, *E*, *F*, G and *H* genes in all known depsipeptide HDAC inhibitor BGCs, strongly suggests a unified mechanism for pharmacophore assembly. It is worth noting, however, that *spiP* and *pcdP* in the spiruchostatin and FR901375 BGCs, respectively, encode ATs (**Fig. 2**) that likely supply malonyl extender units to PcdB/SpiB and PcdC/SpiC, in place of FabD1.

**Figure 6.**
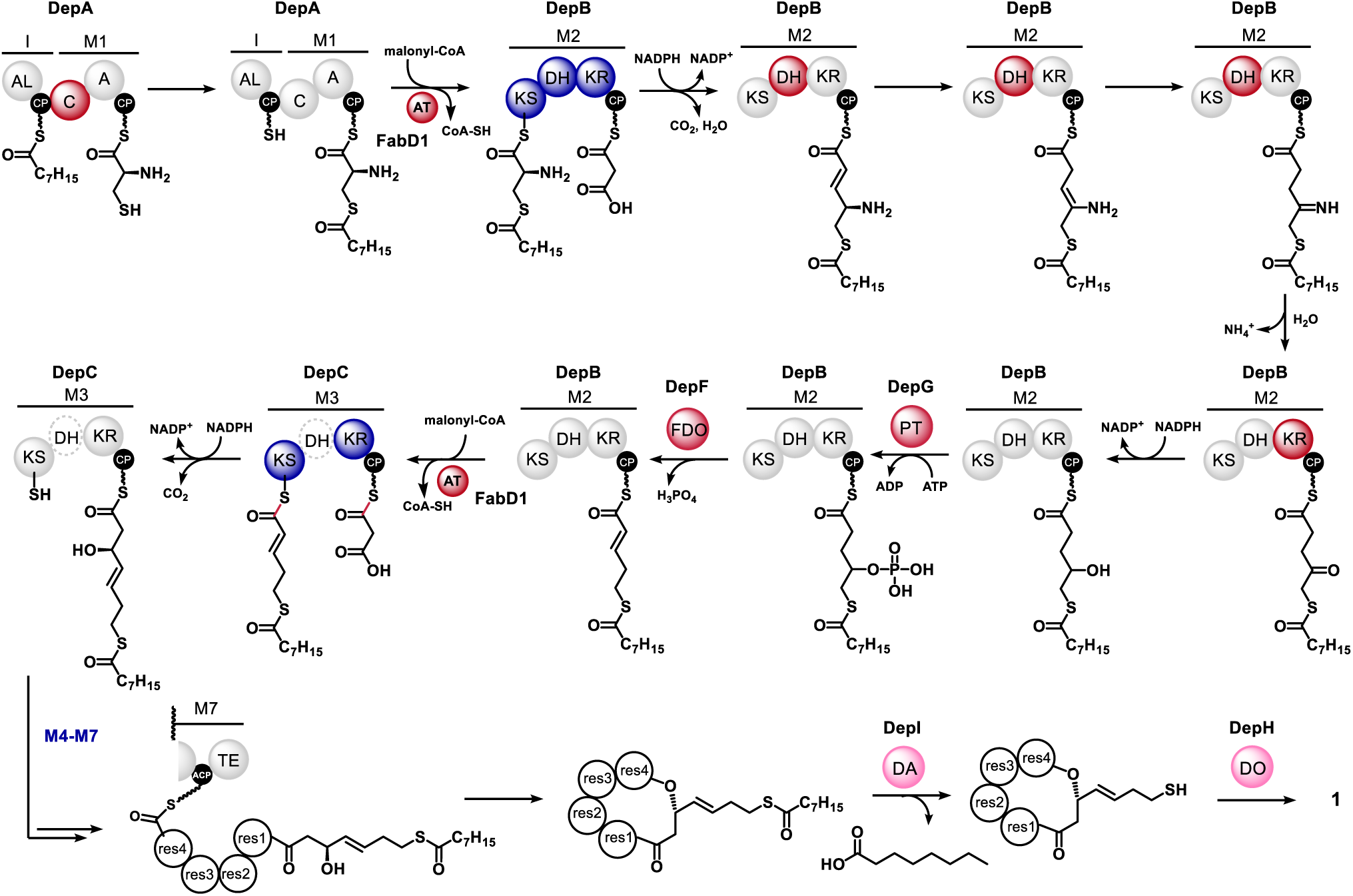
Proposed pathway for romidepsin biosynthesis showing each CP-bound intermediate in pharmacophore biosynthesis. NRPS or PKS domains and *trans*-acting enzymes responsible for canonical and non-canonical transformations are highlighted in dark blue and red, respectively. Enzymes catalysing post-NRPS de-octanoylation and disulfide formation are coloured pink. The biosynthesis of the other disulfide-containing members of the depsipeptide HDAC inhibitor family is hypothesised to employ an identical pathway for pharmacophore assembly, de-acylation and disulfide formation, expect that the ATs encoded by *spiP* and *pcdP* are used in place of FabD1 in spiruchostatin and FR901375 biosynthesis.

## Discussion

In this study we report, to the best of our knowledge, the first example of nitrogen deletion in small molecule biosynthesis, significantly expanding the scope of Nature’s functional group deletion repertoire. Our bioactivity evaluation of an FR901375 analogue bearing a hydroxyl group at the site of nitrogen deletion highlights that removal of branching heteroatoms from the portion of the conserved pharmacophore occupying the active site tunnel in HDAC-targeting depsipeptides is critical for potent inhibition. This is likely due to a combination of steric factors and the energetic cost of desolvating a polar functional group upon inhibitor binding.

Reconstitution of pharmacophore assembly in vitro using the conserved components from romidepsin biosynthesis revealed some intriguing features. The first is that the thiol of a CP-bound cysteine residue is protected by octanoylation via the action of a unique NRPS C domain at the onset of pharmacophore assembly. This protecting group appears to remain in place throughout depsipeptide biosynthesis, only being cleaved in the penultimate step by a deacylase, enabling formation of the disulfide pro-drugs. Intriguingly, in largazole, a biosynthetically uncharacterised depsipeptide HDAC inhibitor, the pharmacophore thiol is also masked by an octanoyl group. This not only suggests a unified biosynthetic pathway for all depsipeptide HDAC inhibitors but affords new opportunities for exploration of their biological activity, in light of recent reports that largazole has oral availability and can cross the blood-brain barrier^46^.

The second intriguing feature of pharmacophore assembly is the pathway for nitrogen deletion, which involves unusual multifunctional PKS domains and the unique *trans*-acting DepG and DepF enzymes. The mechanisms of DepF-catalysed conversion of the γ-phosphoryloxy thioester to an α, β-enoyl thioester, and for recruitment of DepG (which is homologous to resistance-conferring aminoglycoside-3’-phosphotransferases) and DepF to the DepB CP domain are fascinating directions for future enquiry.

Building on our characterisation of the novel chain initiation and nitrogen deletion mechanisms, we reconstituted the entire pathway for pharmacophore assembly in a single vessel. This remarkable process employs eight purified recombinant enzymes and four exogenous coenzymes to assemble the *S*-octanoylated pharmacophore as the major product from three building blocks via a cascade of at least seventeen distinct reactions. Future work to optimise this process, for example by incorporation of stoichiometric cofactor regeneration and flux balancing strategies, could result in a fast, scalable, and sustainable pharmacophore synthetic route that underpins rapid solid-phase peptide synthesis-based approaches to the creation of novel analogues of depsipeptide HDAC inhibitors.

Overall, our reconstitution and dissection of the pathway for conserved pharmacophore assembly in depsipeptide HDAC inhibitor biosynthesis provides significant new insight into NRPS and PKS catalytic versatility and complexity, showcasing novel enzymatic cascades for chemically challenging nitrogen deletion.

## Extended Data Figures

**Extended Data Fig 1.**
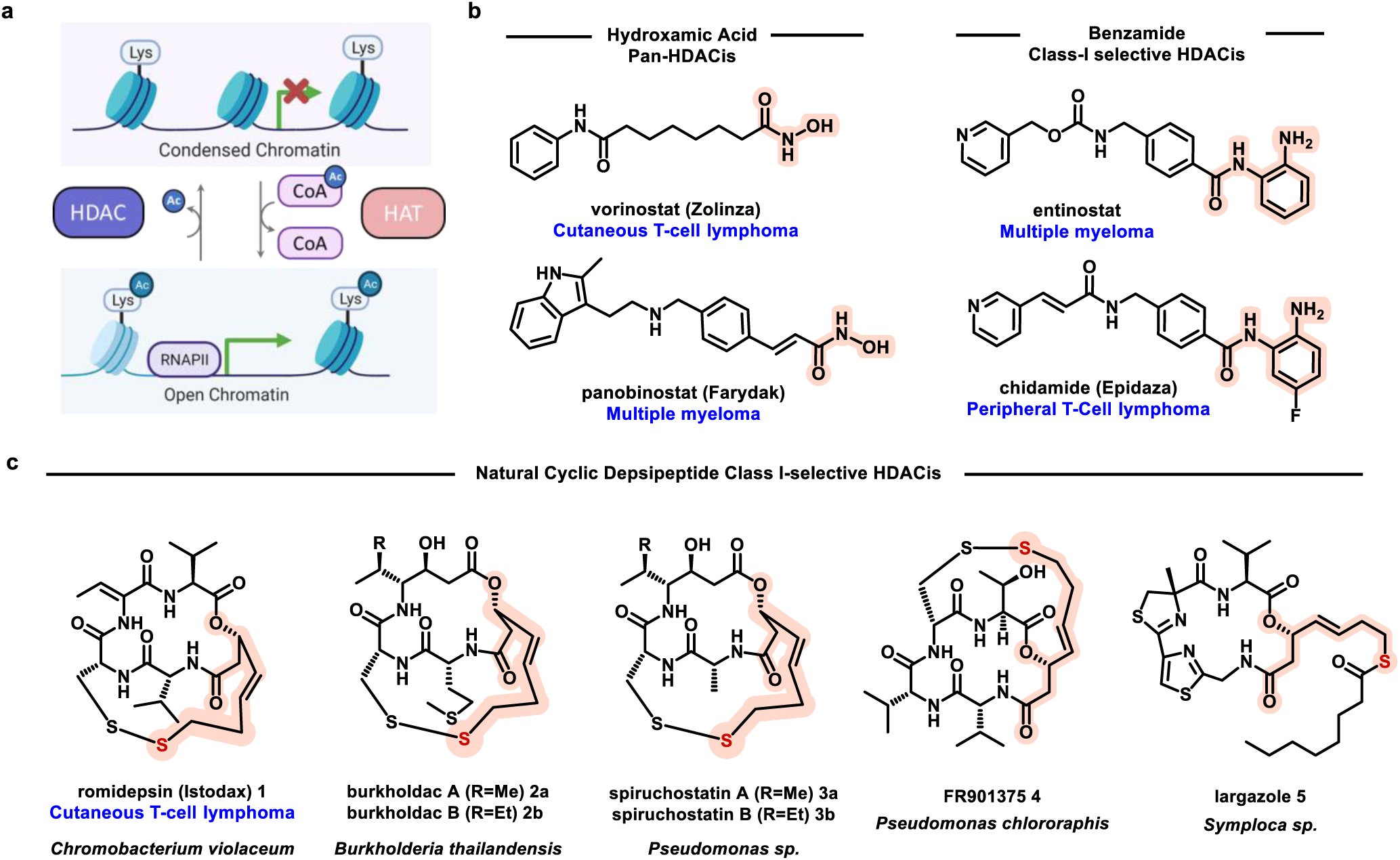
Role played by HDACs in regulating gene expression and HDAC inhibitors as cancer chemotherapeutics. **a**, Functions of HDAC and HAT in epigenetic regulation of gene expression. **b**, Synthetic HDAC inhibitors approved for the treatment of multiple myeloma and cutaneous / peripheral lymphoma. vorinostat, entinostat and panobionstat are approved by the U.S. Food and Drug Administration, and Chidamide is approved by the China Food and Drug Administration. Zinc-binding groups are highlighted in orange. **c**, Depsipeptide HDAC inhibitors produced by beta and gamma-proteobacteria and marine cyanobacteria. These contain a conserved zinc-binding pharmacophore highlighted in orange. Romidepsin is approved by the U.S. Food and Drug Administration for treatment of cutaneous T-cell lymphoma.

**Extended Data Fig 2.**
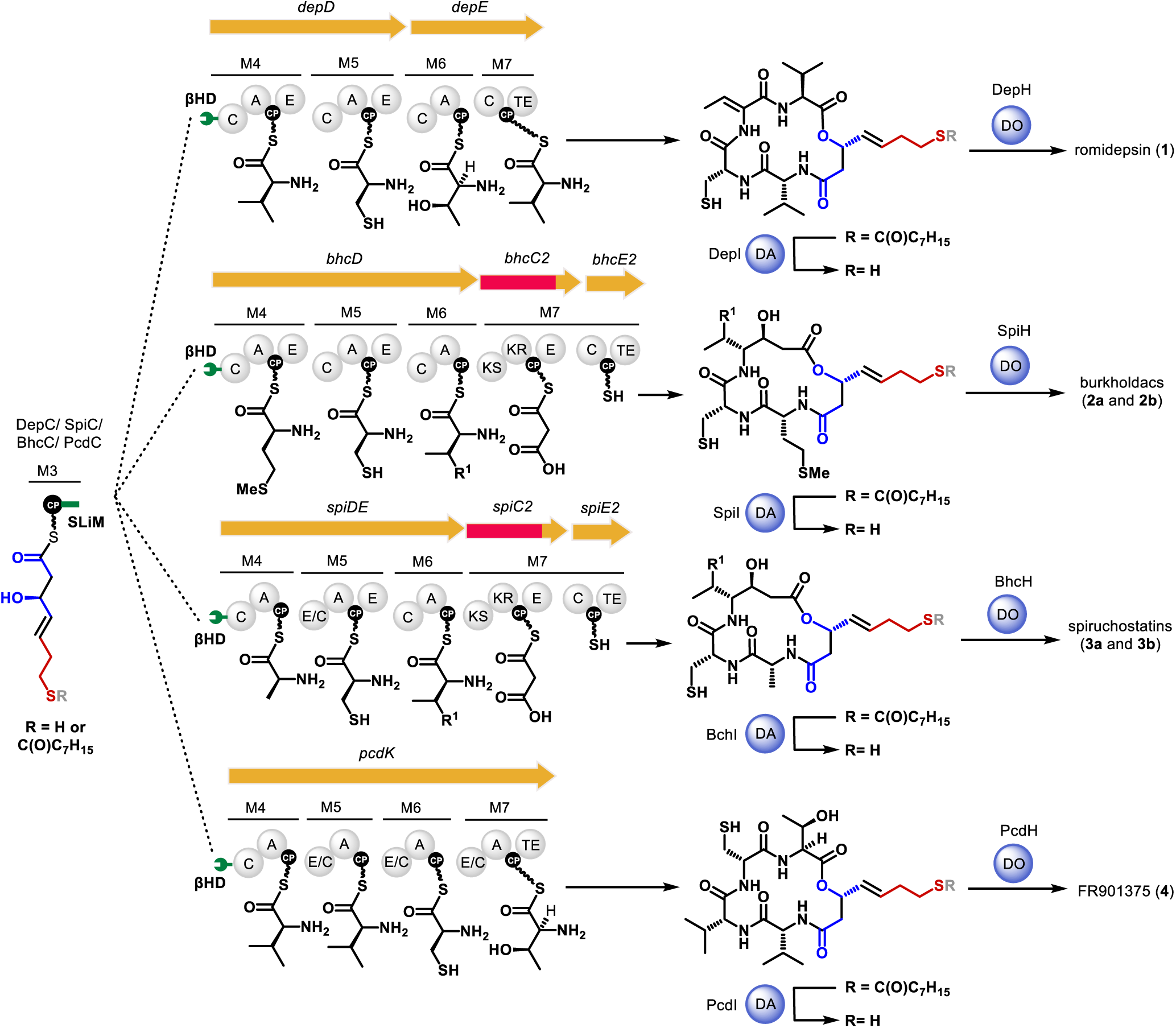
Originally proposed and revised pathways for peptidyl cap assembly and disulfide formation in depsipeptide HDAC inhibitor biosynthesis. In the originally proposed pathway,^21,29,30^ the conserved pharmacophore is assembled with a free thiol that is bonded to the cysteine thiol in each peptidyl cap by a dithiol oxidase (DO). In the revised pathway the pharmacophore thiol is protected with an octanoyl group that is unmasked after assembly of the peptidyl caps by a de-acylase (DA) prior to DO-catalysed disulfide formation.

**Extended Data Fig 3.**
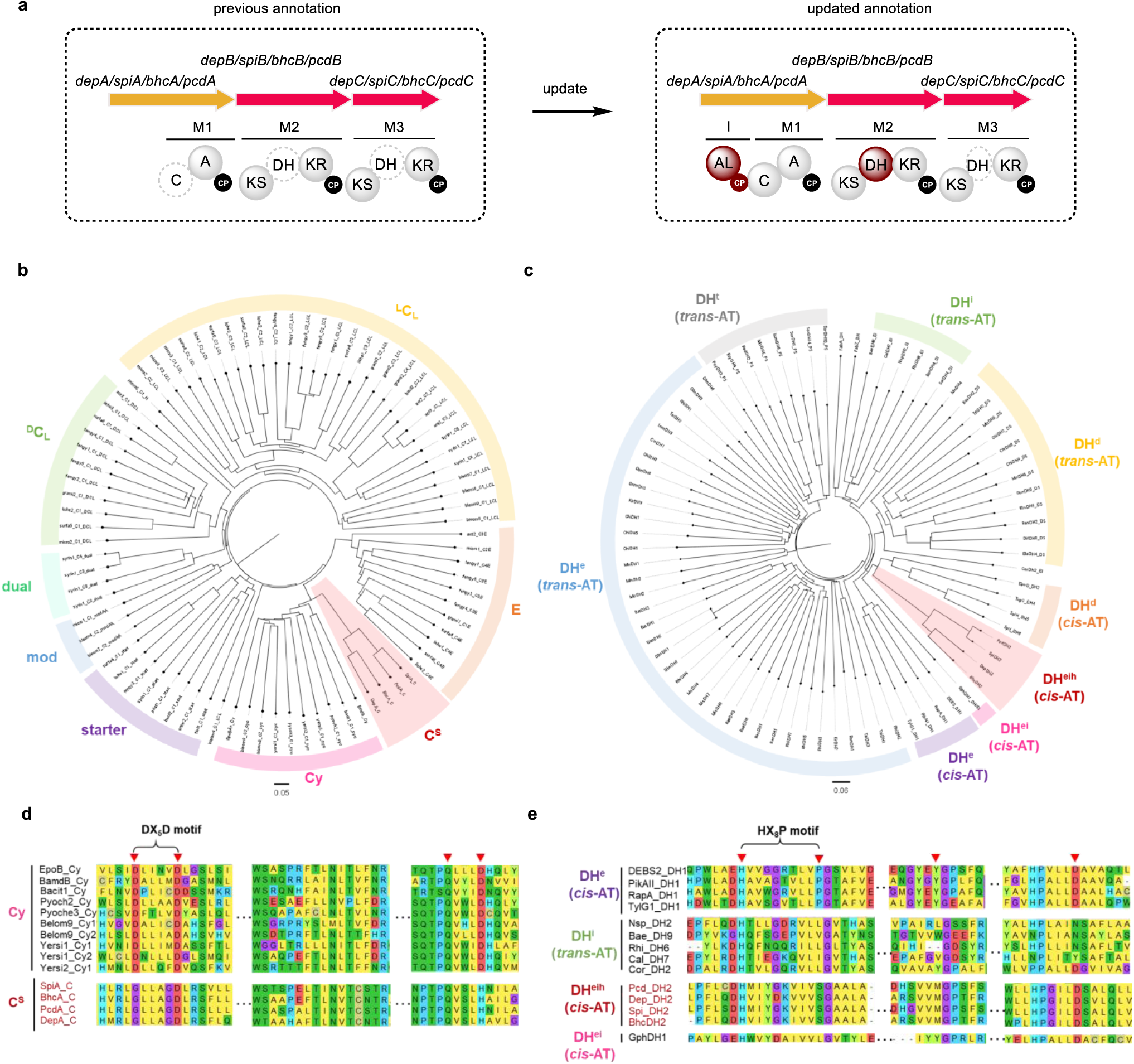
Bioinformatics analysis of A, B and C subunits of depsipeptide HDAC inhibitor assembly lines. **a**, Revised domain and module annotation of A, B and C subunits. Domains re-annotated as functional or missing from the original annotation are highlighted in dark red. Domains proposed to be inactive are denoted with dashed spheres. **b**, Phylogenetic analysis of module 1 C domains, ^L^CL: L-aminoacyl-PCP condensing C domains, ^D^CL: D-aminoacyl-PCP donor and a L-aminoacyl-PCP acceptor condensing C domains, E: epimerisation domains, Cy: heterocyclization domains, starter: chain-initiating C domains, mod: dehydrating C domains, dual: dual function epimerising and condensing C domains, C^S^: thioester synthesising C domains (module 1 C domains in this work). **c**, Phylogenetic analysis of module 2 DH domains. DH^e^: enoyl thioester-forming domains, DH^d^: 2, 4-dienoyl thioester forming domains, DH^T^: tetrahydropyran/tetrahydrofuran-forming domains, DH^i^: enoyl isomerising domains, DH^ei^: bifunctional enoyl thioester-forming and isomerising domains, DH^eih^: trifunctional enoyl thioester-forming, isomerising, and imine hydrolysing domains (module 2 DH domains in this work), *trans*-AT: *trans*-acyltransferase PKSs, *cis*-AT: *cis*-acyltransferase PKSs. **d**, Sequence alignment of reported Cy domains and module 1 C^S^ domains. **e**, Sequence alignment of reported DH-like domains and module 2 DH domains.

**Extended Data Fig 4.**
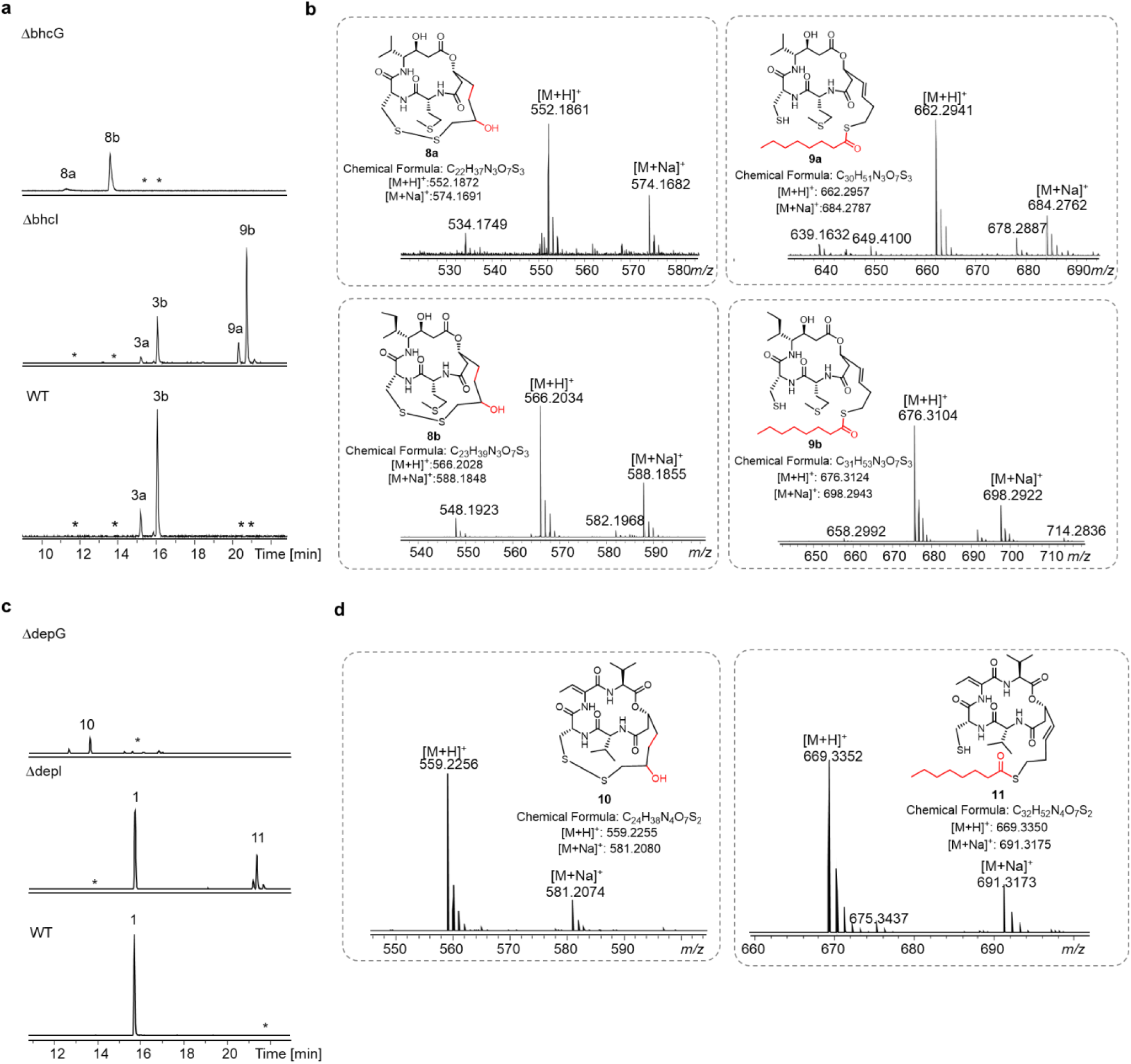
UHPLC-ESI-Q-ToF-MS analysis of intermediates and shunt metabolites accumulated in gene deletion mutants. **a**, Extracted ion chromatograms corresponding to [M+H]^+^ ions for **8a**/**b**, **9a**/**b** and burkholdac A/B **3a**/**b** from *bhcG* mutant (top), *bhcI* mutant (middle) and wild type (WT) *B. thailandensis* DSM13276 (bottom), respectively. **b**, Observed and calculated *m/z* values for [M+H]^+^ ions of **8a/b** and **9a**/**b**. **c**, Extracted ion chromatograms corresponding to [M+H]^+^ ions for **10**, **11** and romidepsin **1** from *depG* mutant (top), *depI* mutant (middle) and wild type (WT) *C. violaceum* BP1968 (bottom). **d**, Observed and calculated *m/z* values for [M+H]^+^ ions of **10** and **11.**

**Extended Data Fig 5.**
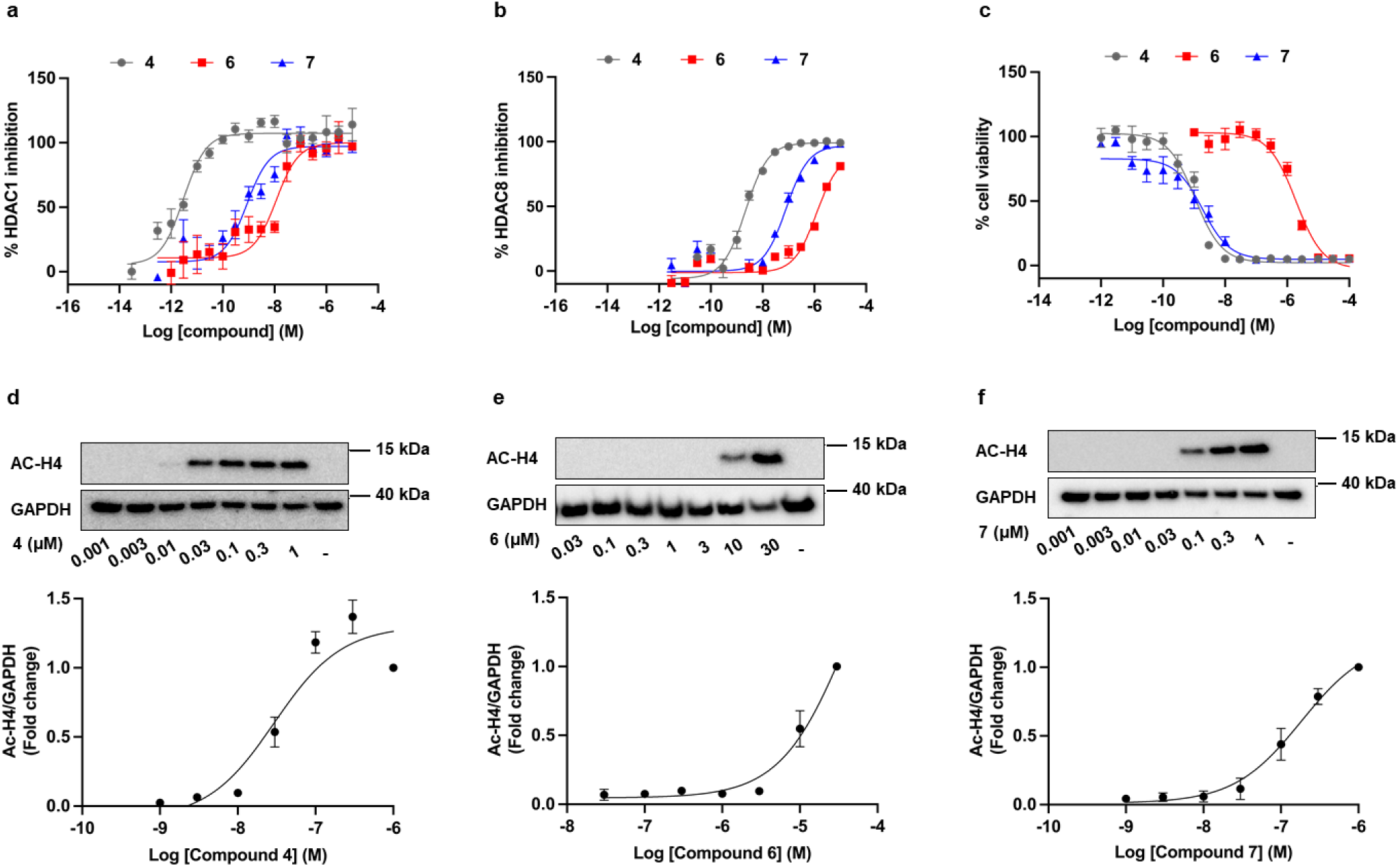
*In vitro* activity of FR901375 and derivatives. **a**, Concentration-dependent inhibition for **4**, **6** and **7** against recombinant human HDAC1. Concentration-dependent inhibition for **4**, **6** and **7** against recombinant human HDAC8. **c**, Comparison of MTT cytotoxicity concentration-response curves for **4**, **6** and **7** against MM96L cells. Cell viability was determined after 72 h of incubation with compounds at the indicated concentrations. **d**-**f:** Concentration-dependent accumulation of acetylated histone H4 induced by **4** (d), **6** (e) and **7** (f), in MM96L cells. Cells were treated with the indicated concentrations of each inhibitor for 24 h and accumulation of acetylated histone H4 (Ac-H4) was measured by western blot. Intensity of Ac-H4 relative to glyceraldehyde-3-phosphate dehydrogenase (GAPDH) was quantified using ImageJ, then normalized to the untreated control as fold change. Blots are representative of three independent experiments and GAPDH was used as the loading control.

**Extended Data Fig 6.**
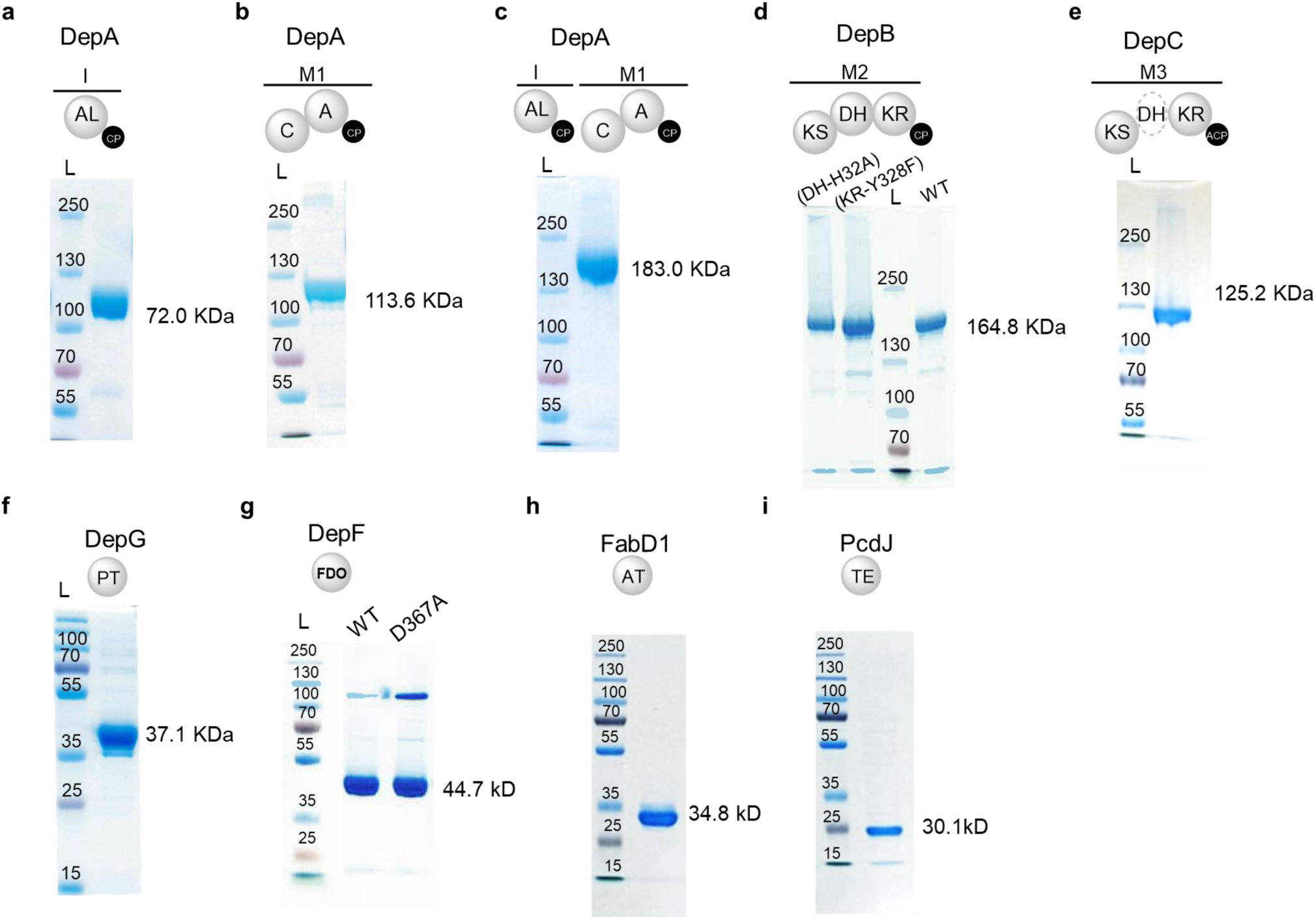
SDS-PAGE analysis of purified recombinant proteins used in this work. Construct compositions are shown on top with site-directed mutation indicated respectively.

**Extended Data Fig 7.**
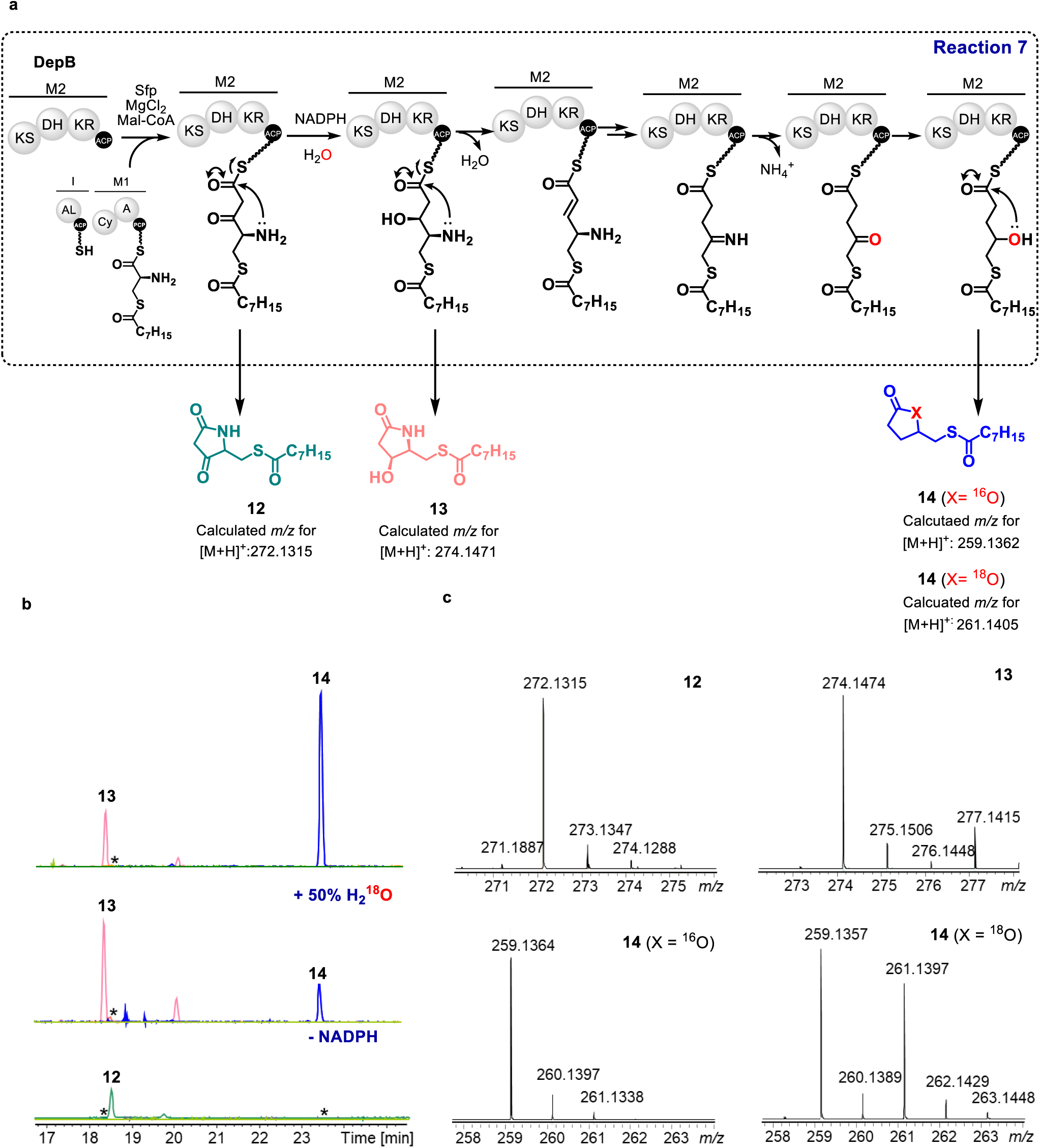
UHPLC-ESI-Q-TOF-MS analysis of products resulting from spontaneous lactamization and lactonization in the module 2 chain elongation assay (Reaction 7). **a**, Schematic representation of Reaction 7 showing proposed structures of CP-bond intermediates and corresponding spontaneously lactamized (**12** and **13**) and lactonized (**14**) products. **b**, Extracted ion chromatograms corresponding to [M+H]^+^ ions for **12**-**14** from Reaction 7 (top), Reaction 7 conducted in buffer made from 1:1 H_2_O/H_2_^18^O (middle), Reaction 7 with NAPDH omitted (bottom). **c**, Observed *m/z* values of [M+H]^+^ ions for **12**-**14**.

**Extended Data Fig 8.**
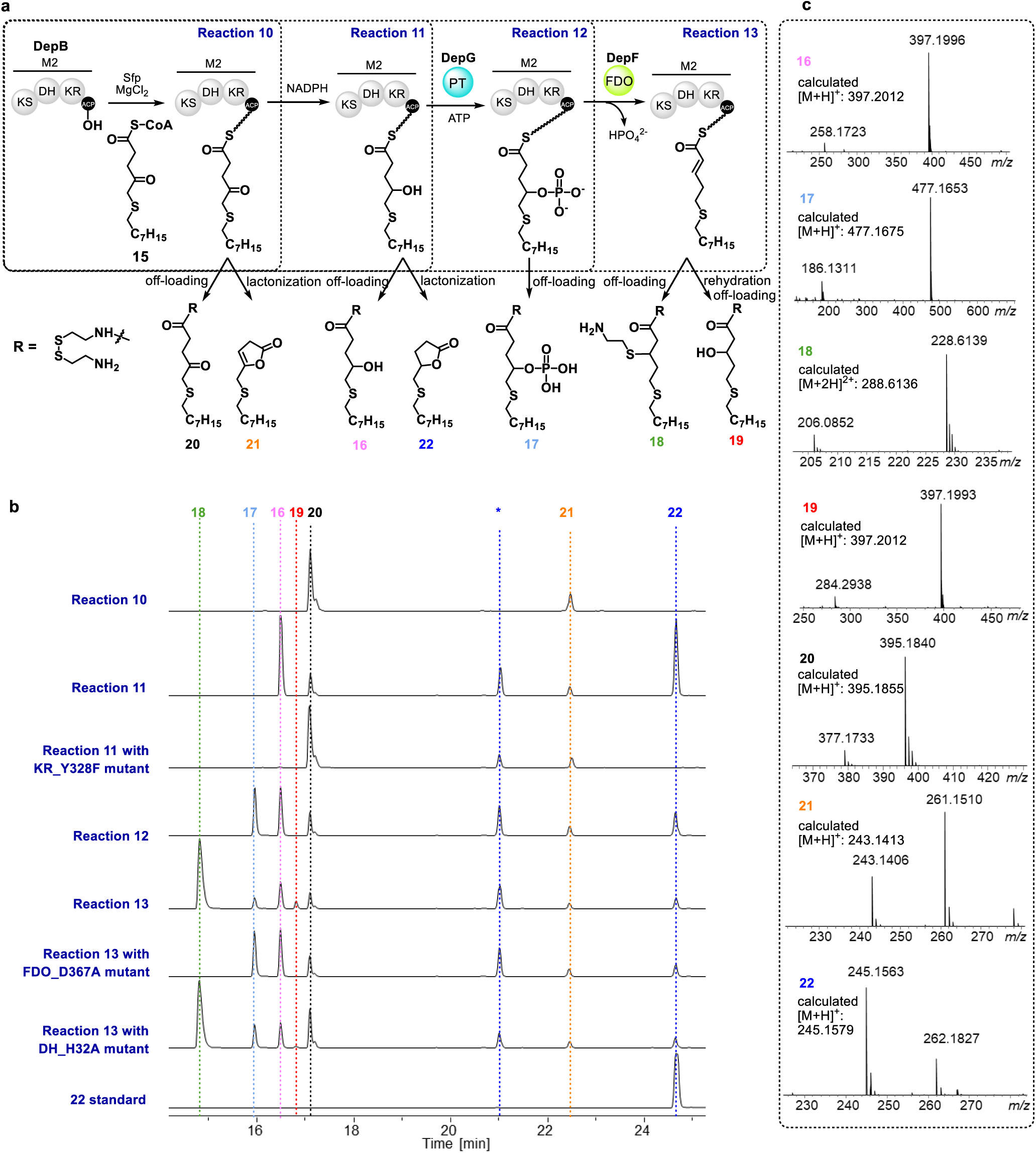
UHPLC-ESI-Q-TOF-MS analyses of *in vitro* assays employing synthetic thioether substrate analogue. **a**, Schematic representation of Reactions 7-10 showing structures of proposed CP-bond intermediates and respective cysteamine off-loaded / spontaneously lactonization / rehydrated products (**16**-**22**) shown. **b**, Extracted ion chromatograms corresponding to [M+H]^+^ ions for products **16**-**22** from reactions 10-13. **c**, calculated and observed [M+H]^+^ values for products **16**-**22**.

**Extended Data Fig 9.**
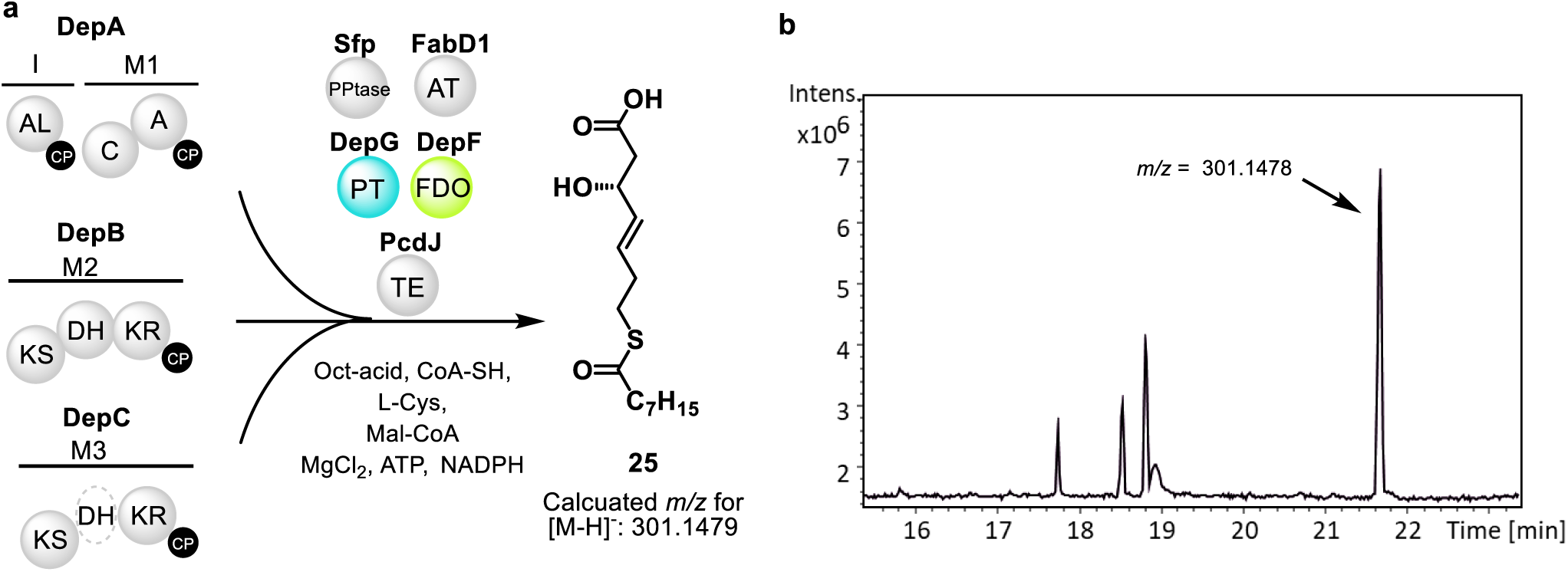
Enzymatic synthesis of octanoylated pharmacophore in a single reaction vessel. **a**, Schematic representation of *S*-octanoylated pharmacophore synthesis in a single reaction vessel; **b**, Base peak chromatogram from UPLC-ESI-Q-ToF-MS analysis of the reaction showing a prominent species at 21.7 min giving rise to an ion with *m/z* = 301.1478, corresponding to the [M-H]^-^ ion of **25** (calculated *m/z* = 301.1479.

## References

1. Li, F., Zhang, X. & Renata, H. Enzymatic C–H functionalizations for natural product synthesis. Curr. Opin. Chem. Biol. 49, 25–32 (2019).

2. Knox, H. L., Sinner, E. K., Townsend, C. A., Boal, A. K. & Booker, S. J. Structure of a B12-dependent radical SAM enzyme in carbapenem biosynthesis. Nature 602, 343–348 (2022).

3. Inahashi, Y. et al. Watasemycin biosynthesis in Streptomyces venezuelae: thiazoline C-methylation by a type B radical-SAM methylase homologue. Chem. Sci. 8, 2823–2831 (2017).

4. Sydor, P. K. et al. Regio- and stereodivergent antibiotic oxidative carbocyclizations catalysed by Rieske oxygenase-like enzymes. Nat. Chem. 3, 388–392 (2011).

5. Cupp Vickery, J. R. & Poulos, T. L. Structure of cytochrome P450eryF involved in erythromycin biosynthesis. Nat. Struct. Biol. 2, 144–153 (1995).

6. Alkhalaf, L. M. et al. Binding of distinct substrate conformations enables hydroxylation of remote sites in thaxtomin D by cytochrome P450 TxtC. J. Am. Chem. Soc. 141, 216–222 (2019).

7. Zhang, Z. et al. Structural origins of the selectivity of the trifunctional oxygenase clavaminic acid synthase. Nat. Struct. Biol. 7, 127–133 (2000).

8. Barry, S. M. et al. Cytochrome P450-catalysed L-tryptophan nitration in thaxtomin phytotoxin biosynthesis. Nat. Chem. Biol. 8, 814 (2012).

9. He, F. et al. Molecular basis for the P450-catalyzed C–N bond formation in indolactam biosynthesis. Nat. Chem. Biol. 15, 1206–1213 (2019).

10. Berkovitch, F., Nicolet, Y., Wan, J. T., Jarrett, J. T. & Drennan, C. L. Crystal structure of biotin synthase, an S-ddenosylmethionine-dependent radical enzyme. Science (80-. ). 303, 76–79 (2004).

11. Roach, P. L. et al. Structure of isopenicillin N synthase complexed with substrate and the mechanism of penicillin formation. Nature 387, 827–830 (1997).

12. McLaughlin, M. I. et al. Crystallographic snapshots of sulfur insertion by lipoyl synthase. Proc. Natl. Acad. Sci. U. S. A. 113, 9446–9450 (2016).

13. Dong, C. et al. Structural biology: Tryptophan 7-halogenase (PrnA) structure suggests a mechanism for regioselective chlorination. Science (80-. ). 309, 2216–2219 (2005).

14. Blasiak, L. C., Vaillancourt, F. H., Walsh, C. T. & Drennan, C. L. Crystal structure of the non-haem iron halogenase SyrB2 in syringomycin biosynthesis. Nature 440, 368–371 (2006).

15. McDonough, M. A. et al. Cellular oxygen sensing: Crystal structure of hypoxia-inducible factor prolyl hydroxylase (PHD2). Proc. Natl. Acad. Sci. U. S. A. 103, 9814–9819 (2006).

16. Fyfe, C. D. et al. Crystallographic snapshots of a B12-dependent radical SAM methyltransferase. Nature 602, 336–342 (2022).

17. Darnet, S. & Schaller, H. Metabolism and biological activities of 4-methyl-sterols. Molecules 24, 451 (2019).

18. Hubbard, B. K., Thomas, M. G. & Walsh, C. T. Biosynthesis of L-p-hydroxyphenylglycine, a non-proteinogenic amino acid constituent of peptide antibiotics. Chem. Biol. 7, 931–942 (2000).

19. Paiva, P. et al. Animal fatty acid synthase: a chemical nanofactory. Chem. Rev. 121, 9502– 9553 (2021).

20. Soohoo, A. M., Cogan, D. P., Brodsky, K. L. & Khosla, C. Structure and mechanisms of assembly-line polyketide synthases. Annu. Rev. Biochem. 93, 471–498 (2024).

21. Cheng, Y. Q., Yang, M. & Matter, A. M. Characterization of a gene cluster responsible for the biosynthesis of anticancer agent FK228 in Chromobacterium violaceum No. 968. Appl. Environ. Microbiol. 73, 3460–3469 (2007).

22. Glozak, M. A. & Seto, E. Histone deacetylases and cancer. Oncogene 26, 5420–5432 (2007).

23. Falkenberg, K. J. & Johnstone, R. W. Histone deacetylases and their inhibitors in cancer, neurological diseases and immune disorders. Nat. Rev. Drug Discov. 13, 673–691 (2014).

24. Taori, K., Paul, V. J. & Luesch, H. Structure and activity of largazole, a potent antiproliferative agent from the Floridian marine cyanobacterium Symploca sp. J. Am. Chem. Soc. 130, 1806– 1807 (2008).

25. Cole, K. E., Dowling, D. P., Boone, M. A., Phillips, A. J. & Christianson, D. W. Structural basis of the antiproliferative activity of largazole, a depsipeptide inhibitor of the histone deacetylases. J. Am. Chem. Soc. 133, 12474–12477 (2011).

26. Furumai, R. et al. FK228 (depsipeptide) as a natural prodrug that inhibits class I histone deacetylases. Cancer Res. 62, 4916–4921 (2002).

27. Ying, Y., Taori, K., Kim, H., Hong, J. & Luesch, H. Total synthesis and molecular target of largazole, a histone deacetylase inhibitor. J. Am. Chem. Soc. 130, 8455–8459 (2008).

28. Micelli, C. & Rastelli, G. Histone deacetylases: structural determinants of inhibitor selectivity. Drug Discov. Today 20, 718–735 (2015).

29. Biggins, J. B., Gleber, C. D. & Brady, S. F. Acyldepsipeptide HDAC inhibitor production induced in Burkholderia thailandensis. Org. Lett. 13, 1536–1539 (2011).

30. Potharla, V. Y., Wang, C. & Cheng, Y. Q. Identification and characterization of the spiruchostatin biosynthetic gene cluster enable yield improvement by overexpressing a transcriptional activator. J. Ind. Microbiol. Biotechnol. 41, 1457–1465 (2014).

31. Wang, C. et al. Thailandepsins: Bacterial products with potent histone deacetylase inhibitory activities and broad-spectrum antiproliferative activities. J. Nat. Prod. 74, 2031–2038 (2011).

32. Wesener, S. R., Potharla, V. Y. & Cheng, Y. Q. Reconstitution of the FK228 biosynthetic pathway reveals cross talk between modular polyketide synthases and fatty acid synthase. Appl. Environ. Microbiol. 77, 1501–1507 (2011).

33. Wang, C., Wesener, S. R., Zhang, H. & Cheng, Y. Q. An FAD-dependent pyridine nucleotide-disulfide oxidoreductase is involved in disulfide bond formation in FK228 anticancer depsipeptide. Chem. Biol. 16, 585–593 (2009).

34. Li, J., Wang, C., Zhang, Z. M., Cheng, Y. Q. & Zhou, J. The structural basis of an NADP+-independent dithiol oxidase in FK228 biosynthesis. Sci. Rep. 4, 4145 (2014).

35. Masschelein, J. et al. A dual transacylation mechanism for polyketide synthase chain release in enacyloxin antibiotic biosynthesis. Nat. Chem. 11, 906–912 (2019).

36. Kosol, S. et al. Structural basis for chain release from the enacyloxin polyketide synthase. Nat. Chem. 11, 913–923 (2019).

37. Passmore, M. et al. Molecular basis for depsipeptide HDAC inhibitor combinatorial biosynthesis. *bioRxiv* 2025.03.13.642309 (2025) doi:10.1101/2025.03.13.642309.

38. Dowling, D. P. et al. Structural elements of an NRPS cyclization domain and its intermodule docking domain. Proc. Natl. Acad. Sci. U. S. A. 113, 12432–12437 (2016).

39. Hobson, C. et al. Diene incorporation by a dehydratase domain variant in modular polyketide synthases. Nat. Chem. Biol. 2022 1812 18, 1410–1416 (2022).

40. Dodge, G. J., Ronnow, D., Taylor, R. E. & Smith, J. L. Molecular basis for olefin rearrangement in the gephyronic acid polyketide synthase. ACS Chem. Biol. 13, 2699–2707 (2018).

41. Tng, J. et al. Achiral derivatives of hydroxamate AR-42 potently inhibit class i HDAC enzymes and cancer cell proliferation. J. Med. Chem. 63, 5956–5971 (2020).

42. Quadri, L. E. N. et al. Characterization of Sfp, a Bacillus subtilis phosphopantetheinyl transferase for peptidyl carder protein domains in peptide synthetases. Biochemistry 37, 1585–1595 (1998).

43. Belecki, K. & Townsend, C. A. Biochemical determination of enzyme-bound metabolites: Preferential accumulation of a programmed octaketide on the enediyne polyketide synthase CalE8. J. Am. Chem. Soc. 135, 14339–14348 (2013).

44. Calderone, C. T., Bumpus, S. B., Kelleher, N. L., Walsh, C. T. & Magarvey, N. A. A ketoreductase domain in the PksJ protein of the bacillaene assembly line carries out both α- and β-ketone reduction during chain growth. Proc. Natl. Acad. Sci. U. S. A. 105, 12809–12814 (2008).

45. Neumann, C. S. & Walsh, C. T. Biosynthesis of (-)-(1S,2R)-allocoronamic acyl thioester by an Fe II-dependent halogenase and a cyclopropane-forming flavoprotein. J. Am. Chem. Soc. 130, 14022–14023 (2008).

46. Al-Awadhi, F. H. et al. Largazole is a brain-penetrant class I HDAC inhibitor with extended applicability to glioblastoma and CNS diseases. ACS Chem. Neurosci. 11, 1937–1943 (2020).

