## Supplementary Information for "Nitrogen deletion in HDAC-targeting anticancer drug biosynthesis"

### Nitrogen deletion in the biosynthesis of HDAC-targeting anticancer drugs

#### Materials

##### Table S1 Strains, plasmids and culture media

| Strains | Descriptions | Media |
|---|---|---|
| *Pseudomonas chlororaphis* DSM21509 | FR901375 producer strain | LB for growing, M3 medium[1] for production |
| *Burkholderia Thailandensis* DSM13276 | Burkholdacs producer strain | LB for growing, M8 medium[1] for production |
| *Chromobacterium violaceum* FERM-BP1968 | Romidepsin producer strain | LB for growing, M1 medium[1] for production |
| *Pseudomonous sp*. FERM-BP6944 | Spiruchostatin producer strains | LB for growing, M3 medium[1] for production |
| *E. coli* S17-1 λpir | host for conjugation | LB with antibiotic |
| *E. coli* Top10 | host for cloning | LB with antibiotic |
| *E. coli* BL21 (DE3) | host for protein expression | LB with antibiotic |

##### Table S2 Primers list

| Name | Sequence (5'→3') | Description |
|---|---|---|
| pcdF-L-For | *agctcggtacccggggatcctctaga*TTCTCGTCGATCTCGGTC | *pcdF* in-frame deletion |
| pcdF-L-Rev | *ttgatatgtc*ACCACTCGCCAGTGCCGC | |
| pcdF-R-For | *ggcgagtggt*GACATATCAATAGCGAAA | |
| pcdF-R-Rev | *gtaaaacgacggccagtgccaagctt*CCGCAGCATACAGCTCTA | |
| pcdF-CK-For | TGTGGTGACGGGCTTGAG | |
| pcdF-CK-Rev | GGGCGATCATCGTTCTGT | |
| pcdG-L-For | *gtacccggggatcctctaga*GTGCAGAACTGGATTGCT | *pcdG* in-frame deletion |
| pcdG-L-Rev | *cctcgcgacc*GAAACGACGGTGTGCTTC | |
| pcdG-R-For | *ccgtcgtttC*GGTCGCGAGGTACCCAAT | |
| pcdG-R-Rev | *CGACGGCCAGTGCCAAGCTT*TCTGGGGTGAAGGACGTC | |
| pcdG-CK-For | TTTATGACGGGCCTGATGAA | |
| pcdG-CK-Rev | CGCATAGGCCTGGGCTTTCC | |
| pcdI-L-For | *agctcggtacccggggatcctctaga*GATTTCATTGTGATCGGT | |

| | | |
|---|---|---|
| pcdI-L-Rev | *gaaaggggtca*TAATCAAGGGTTTTGCCTTCA | |
| pcdI-R-For | *aaacccttgatt*ATGACCCCTTTCTTGCCGGA | |
| pcdI-R-Rev | *gtaaaacgacggccagtgccaagct*TATCCATATCGGGATCGTT | *pcdI* in-frame |
| pcdI-CK-For | GCGAAACCATGGAAACTTCG | deletion |
| pcdI-CK-Rev | GCGTGATAGGTAGATTGCAT | |
| pcdFG-L-For | *agctcggtacccggggatcctctaga*TTCTCGTCGATCTCGGTC | |
| pcdFG-L-Rev | *cctcgcgacc*ACCACTCGCCAGTGCCGC | *pcdF* and *pcdG* |
| pcdFG-R-For | *ggcgagtggt*GGTCGCGAGGTACCCAAT | double |
| pcdFG-R-Rev | *cgacggccagtgccaagctt*TACATGGAGCTGCTGGCA | in-frame |
| pcdFG-CK-For | TGTGGTGACGGGCTTGAG | deletion |
| pcdFG-CKRev | GGGCGATCATCGTTCTGT | |
| DepA_LM-For | ACTCCATATGTATTCCTCTTCCGATGTCGCATCGC | DepA_AL-CP |
| DepA_LM-Rev | ATATGGATCCTCAAGGCGCAGCGCTTCCATC | expression |
| DepA_M1-For | ATATCATATGGCTGCGCCTGTTTCGGTT | DepA_C-A-CP |
| DepA_M1-Rev | ATATGGATCCTTACAAAGCCTCCTTGCGTGCGC | expression |
| DepA-For | ACTCCATATGTATTCCTCTTCCGATGTCGCATCGC | DepA |
| DepA- Rev | ATATGGATCCTTACAAAGCCTCCTTGCGTGCGC | expression |
| DepB-For | GCGCCATATGACTCACTCAACACC | DepB |
| DepB-Rev | AGCTAAGCTTTCATCGCGCGCCCTTTCTA | expression |
| DepC-For | ATATGCTAGCATGAGCCAAATCGACAC | DepC |
| DepC-Rev | ATCGAAGCTTTCATAGCGTGATTTCCTC | expression |
| DepF-For | ATGCCATATGAATACACGAGTGAAGGC | DepF |
| DepF-Rev | ATATGGATCCTCATGCCAGCTCCCG | expression |
| DepG-For | ATGCCATATGAGCGCGCGAATAGC | DepG |
| DepG-Rev | ATATGGATCCTCATAGGCGGCCGC | expression |
| FabD1- For | *gtgccgcgcggcagccatATGGCGTTTGCATTTCTGTTCC* | FabD1 |
| FabD1- Rev | *acggagctcgaattcggatccTCAGTTCAGTTCGCCGCGAG* | expression |
| PcdJ-For | ATACATATGACCATCGGACCGCTTGGCTC | PcdJ |
| PcdJ-For | ATACTCGAGCTAGAGCAACACAGACCGCAG | expression |

### Methods

#### Construction of in-frame gene deletion mutants

Homologous arms (500 bp - 1000 bp) flanking a target gene were amplified from *P. chlororaphis* DSM21509 genomic DNA using primers listed in Table S2, seamlessly fused together by overlap PCR and then cloned into the suicide vector pK18mobsacB or pT18mobsacB in *E. coli* S17-1 λpir strain to make the in-frame gene deletion plasmid construct.

The gene deletion construct was transferred into host via bi-parental mating and exconjugants were selected on LB agar plate containing appropriate antibiotics (ampicillin 100 µg/mL and kanamycin 50 µg/mL for *P. chlororaphis* DSM21509; apramycin 100 µg/mL and tetracycline 100 µg/mL for *B. Thailandensis* DSM13276; ampicillin 100 µg/mL and tetracycline 50 µg/mL for *Pseudomonous sp*. FERM-BP6944; ampicillin 100 µg/mL and tetracycline 10 µg/mL for *C. violaceum* FERM-BP1968). The exconjugants were first confirmed as single crossover mutant by PCR and then grown on LB (15% sucrose) agar plate. The resulting colonies were then screened for phenotype of kanamycin sensitive (when using pK18mobsacB based plasmid) or tetracycline sensitive (when using pT18mobsacB plasmid). The colonies with correct phenotype were screened by colony PCR for gene deletion mutant.

#### Metabolite production, crude extraction and UHPLC-ESI-Q-TOF-HRMS analysis

*Burkholderia* strain was grown in 5 ml LB medium at 30 °C, 180 rpm for overnight as seed culture, which was used to inoculate 50 ml production media (table S1) supplemented with 1 % (w/v) HP-20 and 1 % (w/v) XAD-16 resins (Thermo Scientific) and cultured at 30 °C, 180 rpm for 72 h. After growing, cells and resins were harvested by centrifuging (4,000 g for 15 min), freeze dried and extracted with 10 ml ethyl acetate. The resulting solution was vacuum dried and residues was dissolved in 1 ml methanol followed with 20 times dilution for UHPLC-HRMS analysis.

The metabolite extract was analysed using a Dionex UltiMate 3000 UHPLC connected to a Zorbax Eclipse Plus column (C18, 100 × 2.1 mm, 1.8 µm) coupled to a Bruker MaXis IMPACT ESI-Q-TOF mass spectrometer. A gradient elution of 20% to 100% MeCN containing 0.1% formic acid at flow rate of 0.2 ml/min over 35 min was performed. The mass spectrometer was operated in positive ion mode with a scan range of 50-3000 *m/z*. Source conditions were: end plate offset at −500 V; capillary at −4500 V; nebulizer gas ($N_2$) at 1.6 bar; dry gas ($N_2$) at 8 L min$^{-1}$; dry temperature at 180 °C. Ion transfer conditions were: ion funnel RF at 200 Vpp; multiple RF at 200 Vpp; quadrupole low mass at 55 *m/z*; collision energy at 5.0 eV; collision RF at 600 Vpp; ion cooler RF at 50–350 Vpp; transfer time at 121 s;

pre-pulse storage time at 1 s. Calibration was performed with 1 mM sodium formate through a loop injection of 20 µL at the start of each run.

#### Metabolite derivatives purification and NMR characterization

Metabolite production and crude extraction was done similarly as above, while at 1-2 L scale. The crude EtOAc extract was re-dissolved in 50% MeOH in water for purification using Agilent 1260 Series HPLC instrument equipped with a HP Agilent 1260 Diode Array detector and an Thermo Scientific™ BetaSil™ C18 column (150 × 21.2 mm, 5 µm). Elution gradient for purification of **6** is: 0 min, 5% MeOH; 5 min, 5% MeOH; 15 min, 35% MeOH; 45 min, 35% MeOH; 60 min, 100% MeOH; 70 min, 100% MeOH at a flow rate of 15 mL/min and a UV monitoring at 210 nm, which afforded the compound **6** (10.0 mg, $t_R$ = 55 min). Elution gradient for purification of **7** is: 0 min, 5% MeOH; 5 min, 5% MeOH; 15 min, 35% MeOH; 45 min, 35% MeOH; 75 min, 100% MeOH; 90 min, 100% MeOH at a flow rate of 15 mL/min and a UV monitoring at 210 nm was applied, yielding the compound **7** (2.0 mg, $t_R$ = 73 min). The two samples were dissolved in 0.6 mL of deuterated MeOH in a Norell® standard series™ 5 mm NMR tube, and 1D/2D spectra ($^1$H, $^{13}$C, COSY, HSQC, HMBC, and NOESY) were obtained on a Bruker Avance III™ HD 500 MHz spectrometer. Chemical shifts (δ) are given in ppm and coupling constants (*J*) are given in hertz (Hz).

#### Biological activity charaterisation of metabolite derivatives

##### Human recombinant HDAC enzyme inhibition assays

Human recombinant HDAC enzyme assays were performed as previously described by J. Tng et al and J. Mak et al.[2,3] Briefly, the inhibition of human recombinant HDAC enzymes assay was assessed in vitro using a black 384-well plate (Corning Inc.). All compounds were dissolved in DMSO stock solutions and diluted to the indicated concentrations in assay buffer (25 mM Tris-HCl, 137 mM NaCl, 2.7 mM KCl, 1 mM MgCl2 and 0.1 mg/mL bovine serum albumin; pH 8) with 0.1mM DTT. For HDAC1 and HDAC8 enzyme assays, various concentrations of inhibitor were pre-incubated with 100 ng/mL HDAC1 (Reaction Biology) or 50 ng/mL HDAC8 (BPS Bioscience) enzyme for 30 min at room temperature on a plate shaker. Substrate Ac-Leu-Gly-Lys(Ac)-AMC (12 µM) for HDAC1, or substrate Ac-Leu-Gly-Lys(trifluroAc)-AMC (20 µM) for HDAC8 were then added and incubated for 90 min at 37°C in the dark. The enzymatic reaction was stopped with the developer solution (1 mg/mL trypsin and 25 µM vorinostat) co-incubated for 15 min at room temperature. Fluorescence intensity was measured using PHERAstar plate reader (BMG Labtech) at 350 nm excitation and 460 nm emission wavelengths.

**MTT Viability Assays**

Cells ($5x10^4$/mL) were added to a 96-well plate and allowed to adhere overnight. Media were then removed and cells were treated with 10% FBS DMEM media supplemented with varying concentrations of inhibitors for 72 h. MTT (3-(4,5dimethylthiazoly-2)-2,5-diphenyltetrazolium bromide), (1mg/mL, 50 μL/well) was incubated with cells for 60 min followed by addition of isopropanol to solubilize the formazan product. Absorbance was measured at wavelength λ = 570 nm using a PHERAstar plate reader. Cytoselectivity was defined as $IC_{50}$. Triplicate measurements were made for each data point from n≥3 independent experiments.

**Western Blot Analysis of Histone H4 Acetylation**

MM96L cells ($3.5x10^5$/mL) were seeded into 48-well plate and allowed to adhere overnight. Cells were treated for 24h with 10% FBS DMEM media supplemented with varying concentrations of inhibitors. After treatment, cells were lysed on ice with Cell Lysis Buffer (Cell Signaling Technology) supplemented with protease/phosphatase inhibitor (Roche). Cell lysates were immediately separated on 4-12% Bis-Tris gels and transferred using iBlot 2 Dry Blotting System. Membranes were then blocked with 5% skim milk for 1 h at room temperature and incubated with primary antibodies acetylated-histone H4 (Lys5, Lys8, Lys12 and Lys16, 1:20,000, ab177790, Abcam) or GAPDH (1:30,000, G9545, Sigma-Aldrich) for 1 h at room temperature. Image acquisition was performed using Amersham Imager 600 (GE Healthcare) and densitometric analysis of bands was quantified using ImageJ.

**Data analysis:**. All data were plotted and analysed using GraphPad Prism Version 10.2.0 for Mac OS X (GraphPad Software). Data point represents mean ± SEM of at least three independent experiments. Dose response curves were plotted using 3 parameters logistic model.

**Cloning and site-directed mutagenesis of protein overexpression constructs**

DNA fragments encoding the proteins of interest were amplified from *C. violaceum* FERM-BP1968 gDNA and cloned into pET-His8-G2K. Site-directed mutagenesis mutant were generated using the Q5® Site-Directed Mutagenesis Kit (New England Biolabs) following the manufacture's instructions. *E. coli* TOP10 cells were used for transformation and single colonies were picked and grown overnight at 37 °C and 180 rpm in LB medium containing 50 mg/mL kanamycin. Plasmids were isolated from overnight culture using the GeneJET Plasmid Miniprep Kit (Thermo Fisher Scientific) and verified by sequencing. The primers used to amplify an mutagenize protein-encoding DNA fragments are listed in Table S1.

#### Recombinant protein overproduction and purification

A single colony of *E. coli* BL21 (DE3) transformants containing the expression constructs was grown in 10 mL LB (50 mg/mL) kanamycin at 37 °C and 180 rpm overnight. 1L LB (50 mg/mL kanamycin) was inoculated with 10 mL overnight culture and grown at 37 °C and 180 rpm until an optical density at 600 nm of 0.6-0.8 was reached. 0.25 mM IPTG was added to culture and growth was continued at 15 °C and 180 rpm for 18-20 h. The cells were harvested by centrifugation (4,000 *g* at 4 °C for 15 min), resuspended in 10 mL of loading buffer (20 mM Tris, 300 mM NaCl, 20 mM imidazole, pH 7.4), and lysed using a cell disruptor (Constant Systems). Lysate was centrifuged (14,000 *g* at 4 °C for 30 min) and the supernatant was filtered and loaded onto a HiTrap Chelating Column (GE HealthCare) that had been pre-loaded with 0.1 M $NiSO_4$ followed by equilibration with loading buffer. The column was then washed with 15 column volume (CV) loading buffer and the proteins were eluted in a stepwise manner using loading buffer containing increasing concentrations of imidazole: 50 mM (5 CV), 100 mM (3 CV), 200 mM (3 CV) and 300 mM (3CV). The proteins of interest was confirmed by SDS-PAGE analysis and was further purified by gel filtration using a Superdex 200pg column (GE Healthcare), as required. Fractions containing the same purified protein were combined, exchanged into storage buffer (20 mM Tris, 300 mM NaCl, 10% glycerol, pH 7.4), concentrated using a VivaSpin centrifuge filter (GE Healthcare) with an appropriate molecular weight cut off (MWCO), snap-frozen in liquid nitrogen and stored at -80 °C.

#### *In vitro* reconstitution assays and UHPLC-ESI-Q-TOF-MS analysis

##### *In vitro* reconstitution of Octanoyl-CoA loading on DepA_AL-CP (Fig. 4a)

**Reaction 1**: 125 µM *apo* DepA_AL-CP was incubated with 8 µM Sfp, 10 mM $MgCl_2$ and 1 mM CoA in reaction buffer in a total volume of 50 µL at room temperature for 30min.

**Reaction 2**: 100uM *holo* DepA_AL-CP from Reaction 1 was mixed with 1mM octanoic acid, 1mM ATP in reaction buffer in a total volume of 50 µlL at room temperature for 1h.

After completion, Reaction 1 and 2 mixtures were analysis by intact protein UHPLC-ESI-Q-TOF-MS analysis, respectively.

##### *In vitro* reconstitution of L-cysteine loading and S-acylation on DepA_C-A-CP (Fig. 4b)

**Reaction 3**: 200 µM *apo* DepA_C-A-CP was incubated with 8 µM Sfp, 10 mM $MgCl_2$ and 1 mM CoA in reaction buffer in a total volume of 50 µL at room temperature for 30 min.

**Reaction 4**: 100 uM *holo* DepA_C-A-CP from Reaction 3 was mixed with 500 µM L-Cys, 1mM ATP, 2mM DTT in reaction buffer in a total volume of 50 µL at room temperature for 1 h.

**Reaction 5**: 50 µM Cys-DepA_C-A-CP from RXN4 was mixed with 50 µM octanoyl-DepA_AL-CP from Reaction 2 in reaction buffer in a total volume of 50 µL at room temperature for 1 h.

After completion, Reaction 3-5 mixtures were analysis by intact protein UHPLC-ESI-Q-TOF-MS analysis, respectively.

***In vitro* reconstitution of nitrogen deletion on DepB (Fig. 4b & Extended Data Fig. 6)**

**Reaction 6**: 250 µM *apo* DepB was incubated with 8 µM Sfp, 10 mM $MgCl_2$ and 1 mM malonyl-CoA in reaction buffer in a total volume of 50 µL at room temperature for 30 min.

**Prior Reaction 7-10:** 100 uM DepA was first converted to octanoyl-Cys-DepA by incubation with 8 µM Sfp, 10 mM $MgCl_2$, 1 mM CoA, 1mM octanoic acid, 500uM L-Cys, 1 mM ATP, 2mM DTT in reaction buffer in a total volume of 50 µL at room temperature for 1h.

**Reaction 7**: 100 uM malonyl-DepB from Reaction 6 was mixed with 50 uM octanoyl-Cys-DepA and 1mM NADPH in reaction buffer in a total volume of 50 µL at room temperature for 2 h. For monitoring $O^{18}$ incorporation in lactonized product, the reaction mixture was immediately concentrated to 25 µL after adding all components and added with 25 µL of reaction buffer prepared in $H_2O^{18}$.

**Reaction 8**: 100 uM malonyl-DepB from Reaction 6 was mixed with 50 uM octanoyl-Cys-DepA, 1mM NADPH, 20 uM of DepG and 1 mM ATP in reaction buffer in a total volume of 50 µL at room temperature for 2 h.

**Reaction 9**: 100 uM malonyl-DepBfrom Reaction 6 was mixed with 50 uM octanoyl-Cys-*holo*-DepA, 1mM NADPH, 20 uM of DepG , 1 mM ATP and 20 uM of DepF in reaction buffer in a total volume of 50 µL at room temperature for 2 h.

After completion, Reaction 6-9 mixtures were analysis by intact protein UHPLC-ESI-Q-TOF-MS analysis, respectively. Reaction 7 mixture was also extracted by 2x100 µL EtAc and the organic phase was dried and dissolved in 100 µL MeOH for UHPLC-ESI-Q-TOF-MS analysis for monitoring lactonized and lactamized products.

***In vitro* reconstitution of DepB, DepG and DepF with 5-(octylthio)-4-keto-pentanoyl-CoA thioester (OTKP-CoA) 15 (Extended Data Fig. 7)**

**Reaction 10**: 250 µM *apo* DepB was incubated with 8 µM Sfp, 10mM $MgCl_2$ and 500 µM **15** in reaction buffer in a total volume of 50 µL at room temperature for 30 min.

**Reaction 11**: 150 μM OTKP-DepB from Reaction 10 was mixed with 1mM NADPH in reaction buffer in a total volume of 50 μL at room temperature for 2 h.

**Reaction 12**: 150 μM OTKP-DepB from Reaction 10 was mixed with 1mM NADPH, 30 μM DepG, 1.5 mM ATP in reaction buffer in a total volume of 50 μL at room temperature for 2 h.

**Reaction 13**: 150 uM OTKP- DepB from Reaction 10 was mixed with 1mM NADPH, 30 μM DepG, 1.5 mM ATP, 30 μM DepF in reaction buffer in a total volume of 50 μL at room temperature for 2 h.

Upon completion Reaction 10-13 were individually added with 0.5 M cysteamine (pH7.0) and incubated at 4 °C for 16 h. The reaction mixture was then extracted by 2x100 μL EtOAc, and the organic phase was dried and dissolved in 100 μL MeOH for UHPLC-ESI-Q-TOF-MS analysis.

**One-pot reconstitution of pharmacophore biosynthesis (Extended Data Fig.9)**

**Reaction 14**: Prior to the one-pot reaction, DepA, DepB and DepC were converted to their *holo* forms by incubating 100 μM of individual proteins with 8 μM Sfp, 10 mM $MgCl_2$ and 500 μM CoA in reaction buffer in a total volume of 50 μL at room temperature for 30 min. The three *holo* proteins (30 μM) were then mixed with 10 μM FabD1, 1 mM malonyl-CoA, 1 mM L-Cys, 2mM DTT, 1 mM ATP, 1 mM octanoic acid, 10 μM DepG, 10 μM DepF in reaction buffer in a total volume of 50 μL at room temperature for 1 h before 5 μM PcdJ was added. The reaction was continued at room temperature for another 2 h. After completion, the reaction mixture was added with 100 μL MeOH to precipitate proteins and centrifuge at 15,000 rpm for 10 min, and the supernatant was used for UHPLC-ESI-Q-TOF-MS analysis.

**Intact protein UHPLC-ESI-Q-TOF-MS analysis**

Purified recombinant proteins and enzymatic assays were analysed using a Bruker MaXis Impac II ESI-Q-TOF-MS connected to a Bruker MaXis Impact II ESI-Q-TOF-MS connected to a Bruker Elute UHPLC fitted with an Phenomenex bioZenTM WidePore C4 RP column (100 x 2.1 mm, 2.6 μm). The column was eluted at 30 °C with a linear gradient of 5%-80% MeCN containing 0.1% formic acid at a flow rate of 0.2 ml/min for 40 min. The mass spectrometer was operated in positive ion mode with a scan range of 200–3000 *m/z*. Source conditions were: end plate offset at −500 V; capillary at −4500 V; nebulizer gas ($N_2$) at 1.8 bar; dry gas ($N_2$) at 9.0 L $min^{-1}$; dry temperature at 200 °C. Ion transfer conditions were: ion funnel RF at 400 Vpp; multiple RF at 200 Vpp; quadrupole low mass at 300 *m/z*; collision energy at 8.0 eV; collision RF at 2000 Vpp; transfer time at 110.0 μs; pre-pulse storage time at 10.0 μs.

**UHPLC-ESI-Q-TOF-MS analysis of offloaded products**

The cysteamine or enzymatic-offloaded products were analysed using by UHPLC-ESI-Q-TOF-MS/MS on a or Bruker MaXis Impact II mass spectrometer coupled to a Bruker Elute UHPLC fitted with an Kinetex Core-Shell C18 column (100 × 2.1 mm, 1.7 µm). Using a flow rate of 0.2 mL/min, the column was eluted by a combination of water and acetonitrile with following condition: 5 % (v/v) acetonitrile for 5 min, 5−80 % (v/v) acetonitrile over 21 min, 80 % (v/v) acetonitrile for 3 min, 80−5 % (v/v) acetonitrile over 2 min and 5 % (v/v) acetonitrile for 3 min. The mass spectrometer was operated in negative ion mode with a scan range of 50-3000 m/z. Source conditions were: end plate offset at −500 V; capillary at −4500 V; nebulizer gas (N2) at 1.6 bar; dry gas (N2) at 8 L min−1; dry temperature at 180 °C. Ion transfer conditions were: ion funnel RF at 200 Vpp; multiple RF at 200 Vpp; quadrupole low mass at 55 m/z; collision energy at 5.0 eV; collision RF at 600 Vpp; ion cooler RF at 50–350 Vpp; transfer time at 121 µs; pre-pulse storage time at 1 µs. Calibration was performed with 1 mM sodium formate through a loop injection of 10 µL at the start of each run.

#### Chemical Synthesis

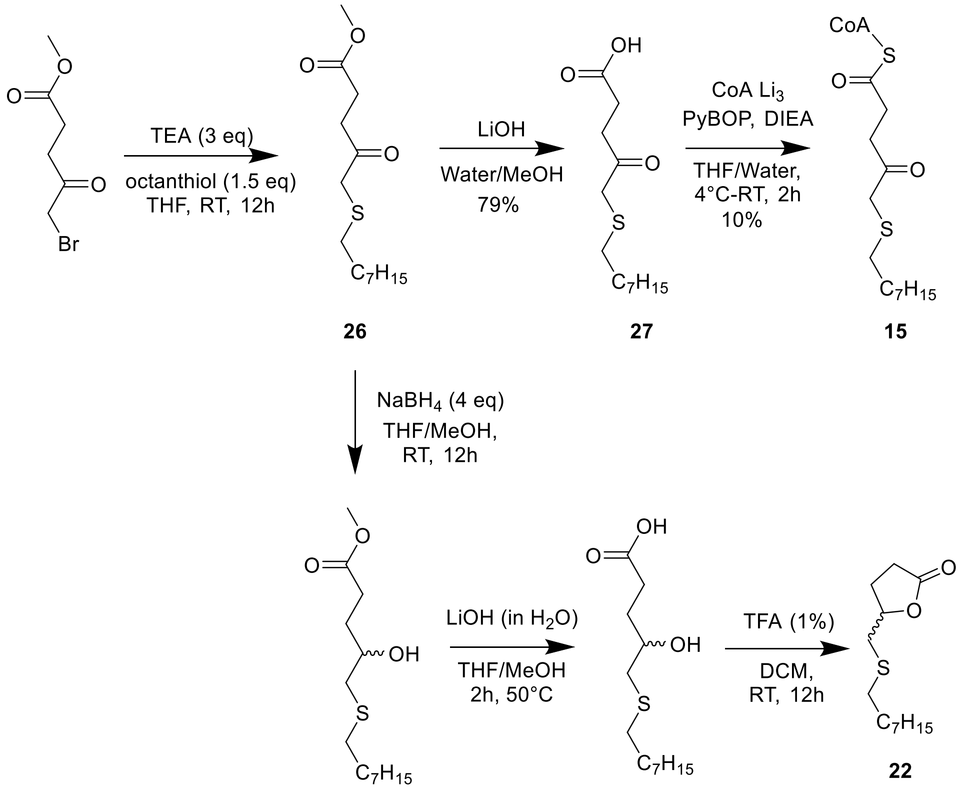


**Scheme 1. Synthetic routes for 5-(octylthio)-4-keto-pentanoyl CoA thioester (15) and 5-((octylthio)methyl)dihydrofuran-2(3H)-one (22).**

**Synthesis of 5-(octylthio)-4-keto-pentanoic acid 27**:

120 µL of octanthiol (1.5 eq) was dissolved in THF with stirring and 200 µL (3.0 eq) of TEA was then added. 100 mg of methyl 5-bromo-4-oxo-pentanoate was dissolved in a small amount of THF and added dropwise to the stirring solution. After ~60 seconds the clear solution became cloudy and the solution was allowed to stir at room temperature for 12h. The solvent was removed in vacuo and the mixture was then dissolved in MeOH : $H_2O$, 1 : 1 and 130 µL of LiOH (1.5 M in water) was then added and the reaction was allowed to stir at 50 °C for 2 h or until complete. The lithium carboxylate was then evaporated under *in vacuo* yielding **27** (colourless needles, 103mg, 79%). (Expected $[M-H]^-$: 259.0, Observed $[M-H]^-$:259.1). $^1$H-NMR (400MHz, $CDCl_3$) $CH_3$ 0.87 (3H, t, J=6.74 Hz), $CH_2$ 1.54 (10H, m), $CH_2CH_2S$ 2.47 (3H, d, J=14.81 Hz), $COCH_2$ 2.65 (2H, d, J=13.04 Hz), $COOHCH_2$ 2.95 (3H, d, J=13.04 Hz), $COCH_2S$ 2.24 (2H, s). $^{13}$C-NMR (100MHz, $CDCl_3$) 204.19 (*C*OCHS), 178.49 ($CO_2H$), 40.98 (CO*C*HS), 34.39 ($CH_2CO$), 32.23 ($CH_2CHS$), 31.78 ($CH_2CH_2$), 29.14 ($CH_2CH_2$), 29.11 ($CH_2CH_2$), 28.93 ($CH_2CH_2$), 28.73 ($CH_2CH_2$), 28.01 ($CH_2CH_2$), 22.62 ($CH_2CH_3$), 14.06 ($CH_3$).

**Synthesis of 5-(octylthio)-4-keto-pentanoyl CoA thioester 15**:

Based on a similar procedure[4], 10 mg of **27** was dissolved in THF on ice with strong stirring where DIEA (16µL, 2.5 eq) and PyBOP (28.4 mg, 1.5 eq) then added. The mixture was allowed to stir for 30 minutes before CoA trilithium salt in water was then added (28.6 mg, 1.0 eq). the final solvent ratio was THF:$H_2O$ 7:3. **15** was then purified by RP-HPLC (3.5mg, 10%). (Expected $[M+H]^+$: 1010.3, Observed $[M+H]^+$:1010.1). $^1$H-NMR (400MHz, $CD_3OD$) Ar-*H* 8.69 (1H, s), Ar-*H* 8.41 (1H, s), N*H* 8.01 (1H, s), Ar-C*H* 6.14 (1H, d, J=5.68 Hz), C*H*O 5.02 (1H, m), C*H*O 4.67(1H, m), C*H*O 4.57(1H, m), $CH_2OP$ 4.40 (2H, m), C*H*O 4.11 (1H, m), C*H*O 4.01 (1H, s), 3.82 (1H, m), $CH_2NH$ 3.49 (2H, m), $CH_2$ 3.02 (6H, m), $CH_2$, OH 2.88 (3H, m), $CH_2$ 2.49 (3H, m), $CH_2$ 2.44 (3H, t, J=6.58 Hz), $CH_2$ 1.58 (2H, m), $CH_2$ 1.34 (12H, m), $CH_2$ 1.08 (3H, m), $CH_3$ 0.93 (6H, m). $^{13}$C-NMR (100MHz, $CD_3OD$) 205.02, 198.10, 174.11, 172.60, 73.78, 48.25, 48.03, 47.82, 47.61, 47.40, 47.18, 46.97, 40.20, 38.82, 38.69, 37.25, 35.58, 35.03, 34.47, 31.61, 31.57, 28.92, 28.83, 28.65, 28.35, 27.81, 22.30, 20.43, 18.38, 13.05. $^{31}$P-NMR (162MHz, $CD_3OD$) -0.66, -11.61, -12.04.

**Synthesis of 5-((octylthio)methyl)dihydrofuran-2(3H)-one 22**:

To 26.2 mg of **26** was added 6mL of THF and 600 µL of MeOH followed by 14.5mg of $NaBH_4$. The reaction was completed after 12 h at RT with partial cleavage of the ester. 40 µL of 1.5 M LiOH in water was then added and the reaction was allowed to stir at 50 °C for 2 h until complete. The solvent was removed *in vacuo* and the mixture was then suspended in 15 mL of DCM where 150 µL of TFA was then added. The solution was monitored by TLC and after 12 h it was washed with water and the aqueous phase was also extracted 1x with DCM. The combined organic phases were dried and the oil was redissolved in DMF and purified by RP-HPLC to yield colourless oil **22** (15.3 mg, 65.5%). (Expected $[M+H]^+$: 245.2 Observed $[M+H]^+$:245.1). $^1$H NMR (400 MHz, $CDCl_3$) δ 4.67 (1H, m), 4.67 (d, *J* = 11.9 Hz, 1H), 2.80 (qd, *J= 13.9 Hz,* 4.8 Hz, 2H), 2.64 (d, *J* = 4.5 Hz, 1H), 2.60 (d, *J* = 3.1 Hz, 1H), 2.57 (d, *J* = 2.8 Hz, 1H), 2.56 (m, 4H), 2.39 (m, 1H), 2.05 (m, 1H), 1.57 (m, 2H), 1.36 (m, 11H), 0.87 (t, *J* = 6.7 Hz, 3H). $^{13}$C-NMR (100MHz, $CDCl_3$) ($\mathit{C}O_2C$)176.74, (*C*HOC) 79.81, ($\mathit{C}H_2CH_2$) 36.61, ($\mathit{C}H_2CH_2$) 33.28, ($\mathit{C}H_2CH_2$) 31.79, ($\mathit{C}H_2CH_2$) 29.70, ($\mathit{C}H_2CH_2$) 29.17, ($\mathit{C}H_2CH_2$) 28.76, ($\mathit{C}H_2CH_2$) 28.57, ($\mathit{C}H_2CH_2$) 26.90, ($\mathit{C}H_2CH_2$) 22.64, ($\mathit{C}H_3$) 14.09.

### Figures

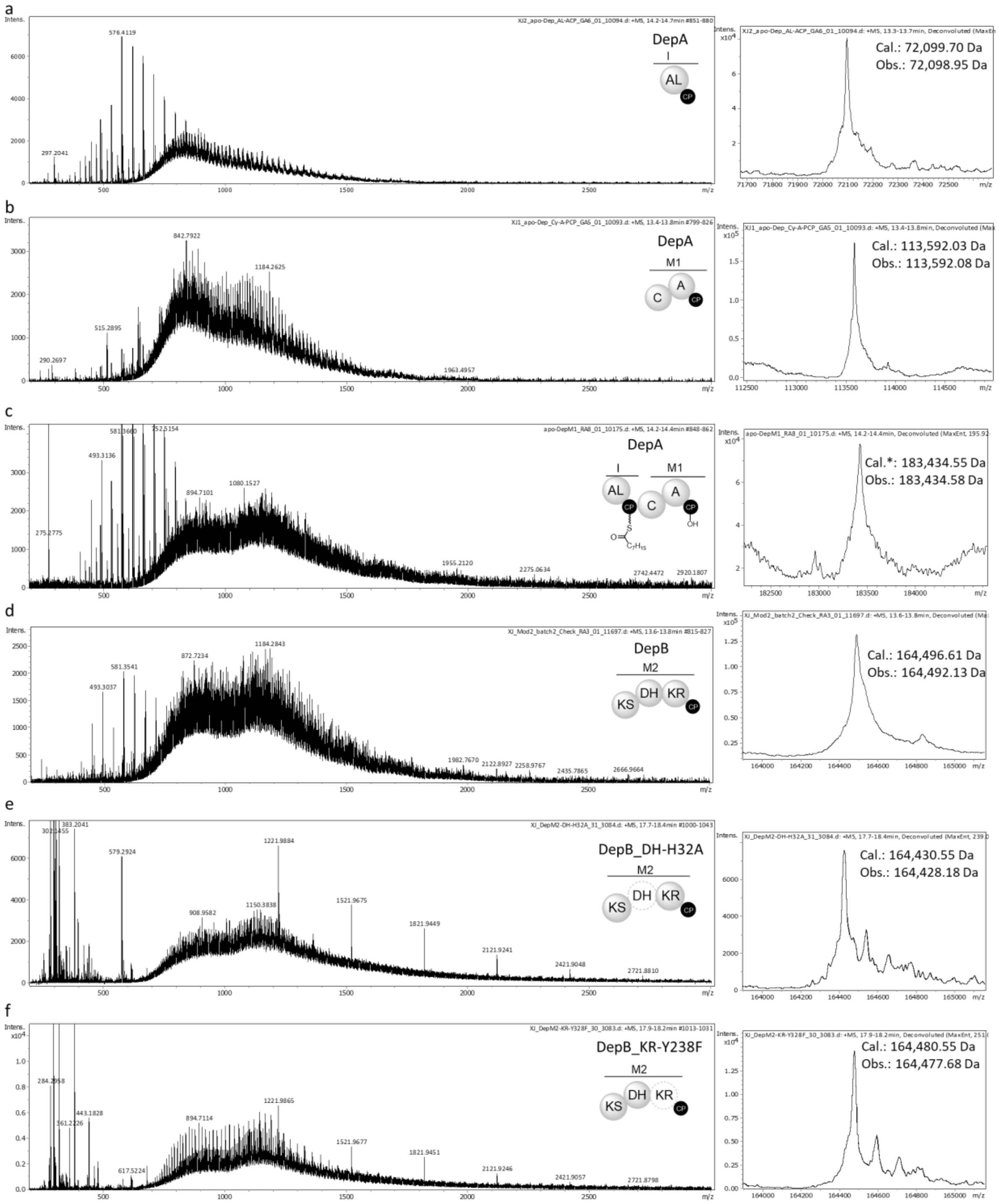


**Figure S1 UHPLC-ESI-QTOF-MS analysis of intact purified proteins.**

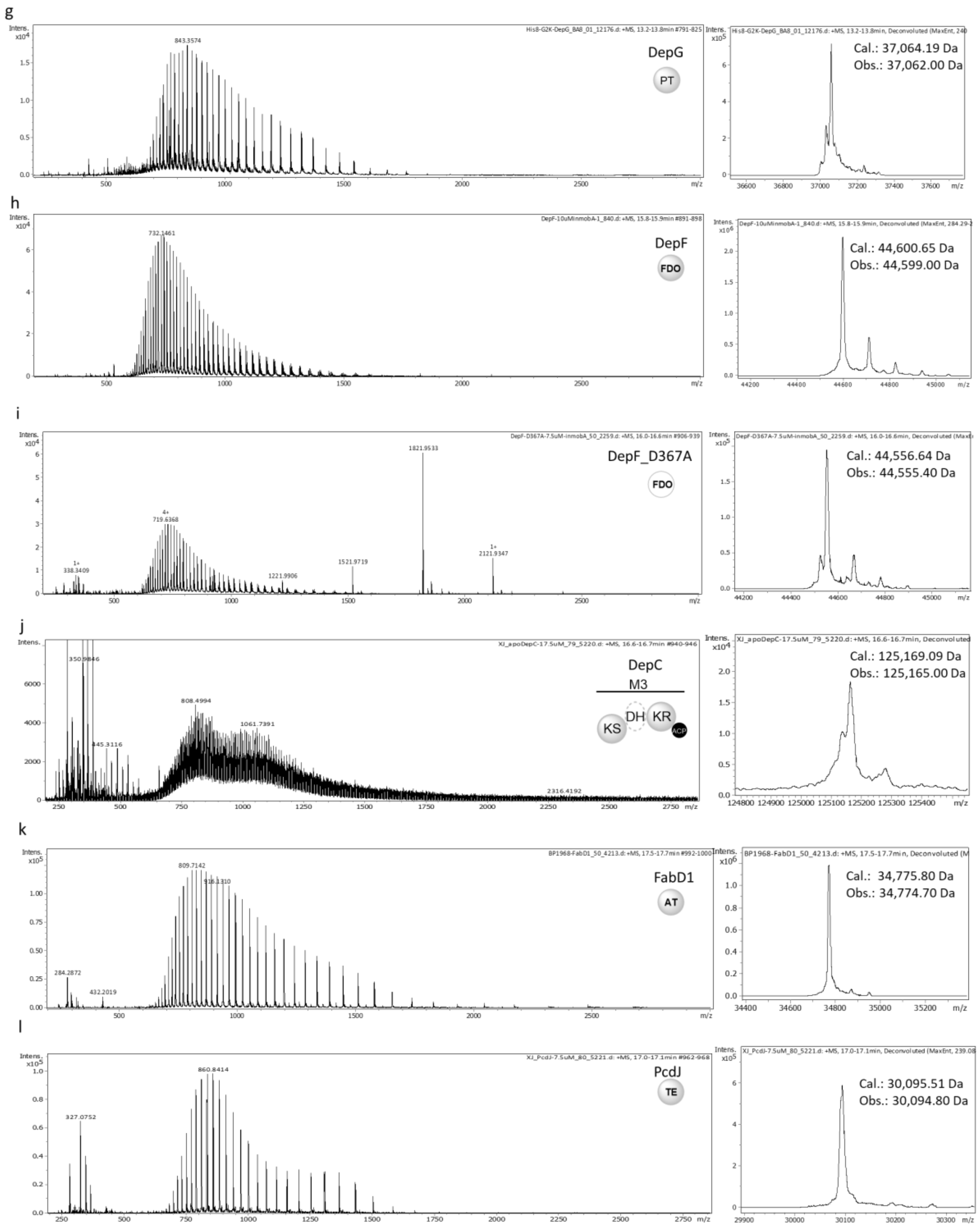


**Figure S1 (continue) UHPLC-ESI-QTOF-MS analysis of intact purified proteins.**

**Table S1. $^1$H (500 MHz) and $^{13}$C (125 MHz) NMR data for 6 in MeOD**

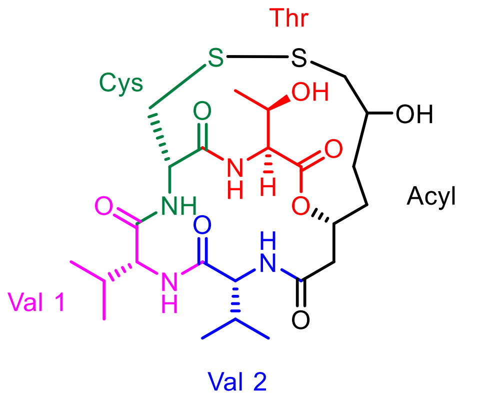


| | Position | $\delta_H$ (*J* in Hz) | $\delta_C$, type |
|---|---|---|---|
| Thr | C=O | - | 171.3, C |
| | 2 | 4.47, d (1.8) | 59.1, CH |
| | 3 | 4.39, m | 68.8, CH |
| | 4 | 1.11, d (6.5) | 20.1, $CH_3$ |
| | NH | 6.94, d (9.3) | - |
| Cys | C=O | - | 171.6, C |
| | 2 | 4.78, dd (8.0, 2.6) | 54.7, CH |
| | 3 | 3.40, dd (14.1, 2.6) / 3.55, dd (14.1, 5.4) | 44.1, $CH_2$ |
| | NH | 8.97, d (8.0) | - |
| Val 1 | C=O | - | 175.5, C |
| | 2 | 3.35, d (10.5) | 68.0, CH |
| | 3 | 2.78, m | 28.8, CH |
| | 4 | 0.95, d (6.7) | 19.9, $CH_3$ |
| | 5 | 1.02, overlapped | 20.7, $CH_3$ |
| | NH | 8.92, d (8.0) | - |
| Val 2 | C=O | - | 174.8, C |
| | 2 | 4.21, d (7.2) | 60.7, CH |
| | 3 | 2.05, m | 32.4, CH |
| | 4 | 1.00, overlapped | 18.7, $CH_3$ |
| | 5 | 1.04, overlapped | 19.9, $CH_3$ |
| | NH | 7.94, d (8.3) | - |
| Acyl | C=O | - | 172.2, C |
| | 2 | 2.65, dd (14.2, 5.2) / 2.84, dd (14.2, 4.3) | 42.0, $CH_2$ |
| | 3 | 5.23, m | 73.4, CH |
| | 4 | 1.89, m / 2.14, m | 26.8, $CH_2$ |
| | 5 | 1.28, m / 1.59, m | 32.7, $CH_2$ |
| | 6 | 4.01, m | 67.2, CH |
| | 7 | 2.34, dd (13.7, 4.2) / 2.92, dd (14.2, 2.5) | 45.8, $CH_2$ |

**Table S2. $^{1}$H (500 MHz) and $^{13}$C (125 MHz) NMR data for 7 in MeOD**

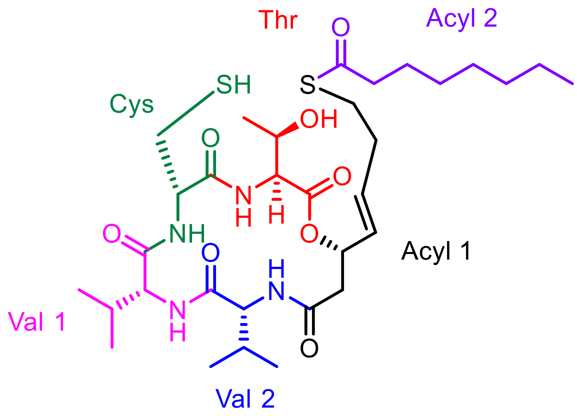


| | Position | $\delta_H$ (*J* in Hz) | $\delta_C$, type |
|---|---|---|---|
| Thr | C=O | - | 169.5, C |
| | 2 | 4.48, d (2.9) | 57.5, CH |
| | 3 | 4.29, m, overlapped | 66.9, CH |
| | 4 | 1.13, d (6.6) | 18.2, $CH_3$ |
| | NH | Not detected | - |
| Cys | C=O | - | 172.6, C |
| | 2 | 4.29, m, overlapped | 56.5, CH |
| | 3 | 3.05, m | 24.1, $CH_2$ |
| | NH | Not detected | - |
| Val 1 | C=O | - | 171.1, C |
| | 2 | 3.81, d (8.1) | 62.4, CH |
| | 3 | 2.44, m | 29.7, CH |
| | 4 | 1.00, overlapped | 18.6, $CH_3$ |
| | 5 | 1.00, overlapped | 18.7, $CH_3$ |
| | NH | Not detected | - |
| Val 2 | C=O | - | 173.3, C |
| | 2 | 4.06, d (7.9) | 60.6, CH |
| | 3 | 2.05, m | 30.1, CH |
| | 4 | 1.00, overlapped | 17.8, $CH_3$ |
| | 5 | 1.00, overlapped | 18.5, $CH_3$ |
| | NH | Not detected | - |
| Acyl 1 | C=O | - | 170.5, C |
| | 2 | 2.61, m / 2.71, m | 39.7, $CH_2$ |
| | 3 | 5.58, m, overlapped | 72.0, CH |
| | 4 | 5.58, m, overlapped | 128.8, CH |
| | 5 | 5.76, m | 131.5, CH |
| | 6 | 2.29, m, | 32.0, $CH_2$ |
| | 7 | 2.92, m | 27.4, $CH_2$ |
| Acyl 2 | C=O | - | 199.6, C |
| | 2 | 2.55, t (7.4) | 43.4, $CH_2$ |
| | 3 | 1.63, m | 25.4, $CH_2$ |
| | 4 | 1.31, m, overlapped | 28.5, $CH_2$ |
| | 5 | 1.31, m, overlapped | 28.6, $CH_2$ |
| | 6 | 1.31, m, overlapped | 22.3, $CH_2$ |
| | 7 | 1.31, m, overlapped | 31.4, $CH_2$ |
| | 8 | 0.90, t (7.1) | 13.0, $CH_3$ |

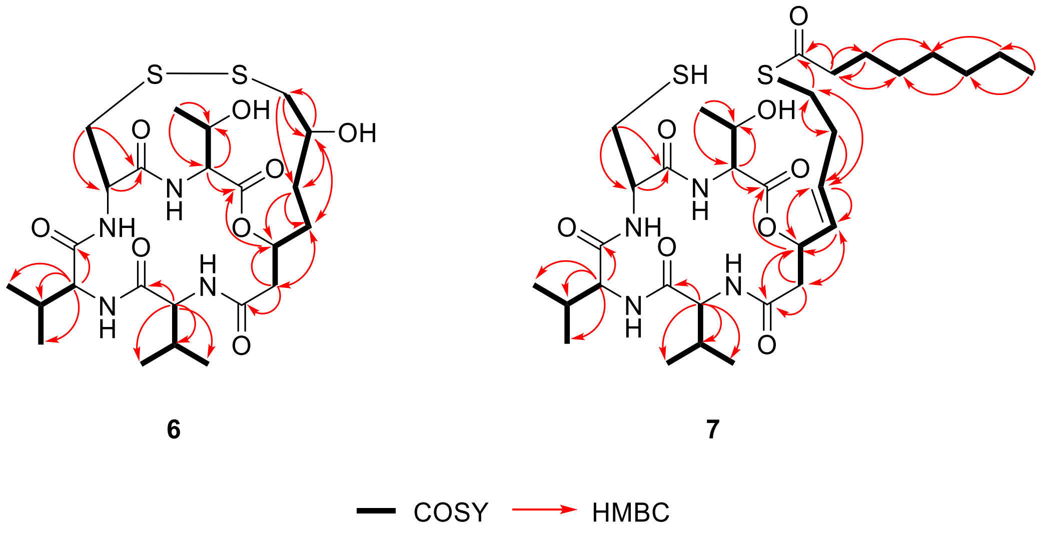


**Figure S3. COSY and HMBC correlations observed for 6 and 7 in NMR spectra**

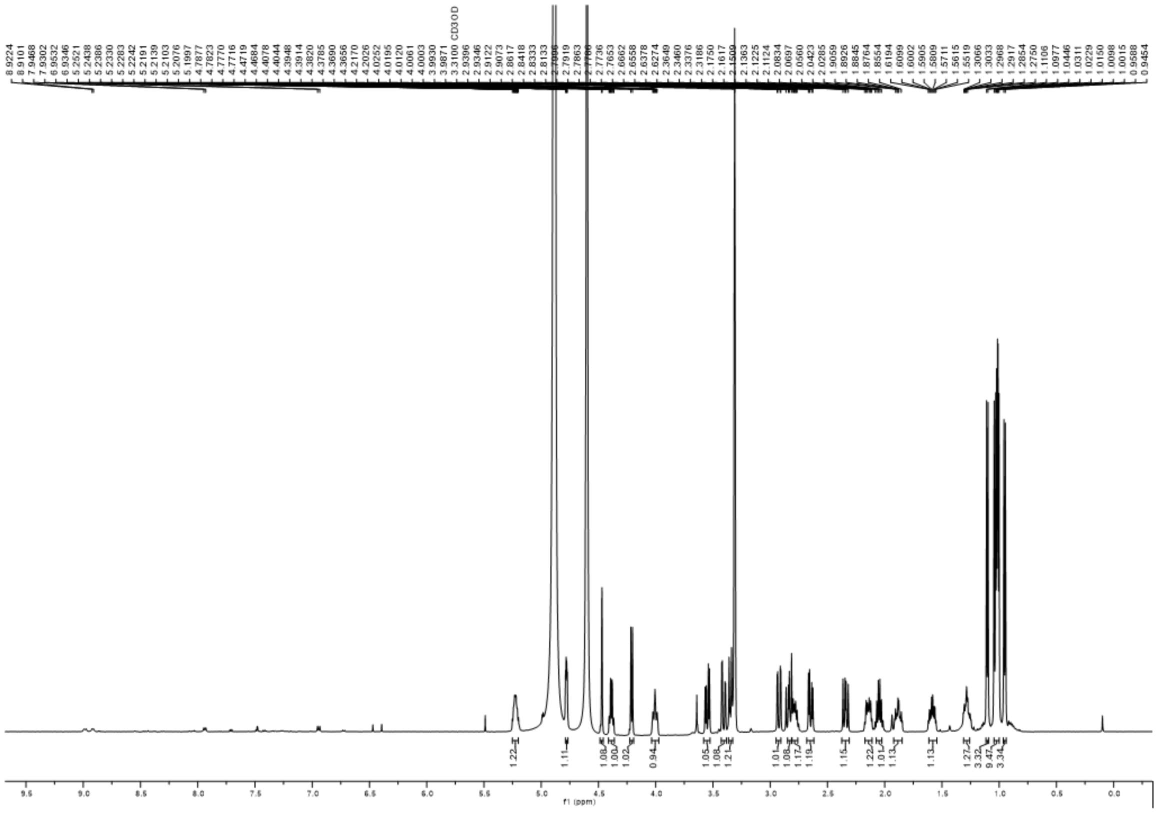


**Figure S4. $^{1}H$ NMR spectrum of 6 in MeOD at 500 MHz**

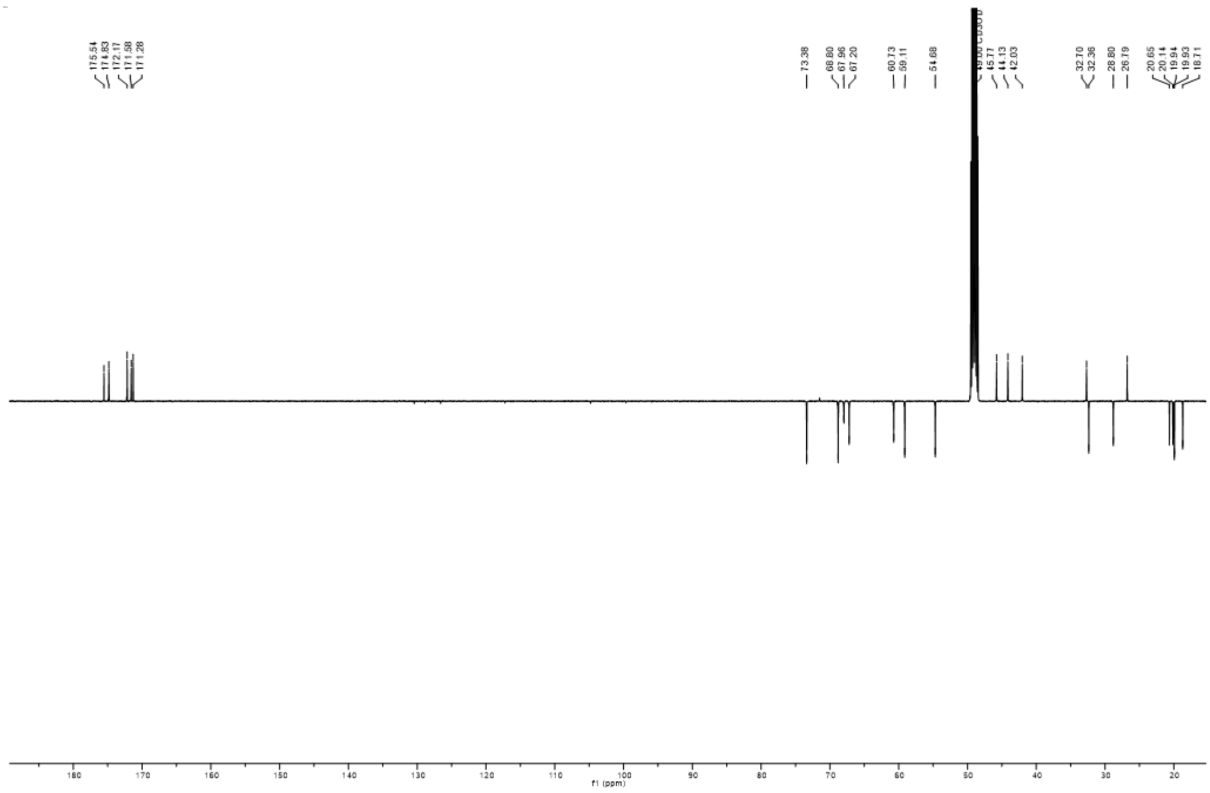


**Figure S5.** $^{13}C$ **NMR spectrum of 6 in MeOD at 125 MHz**

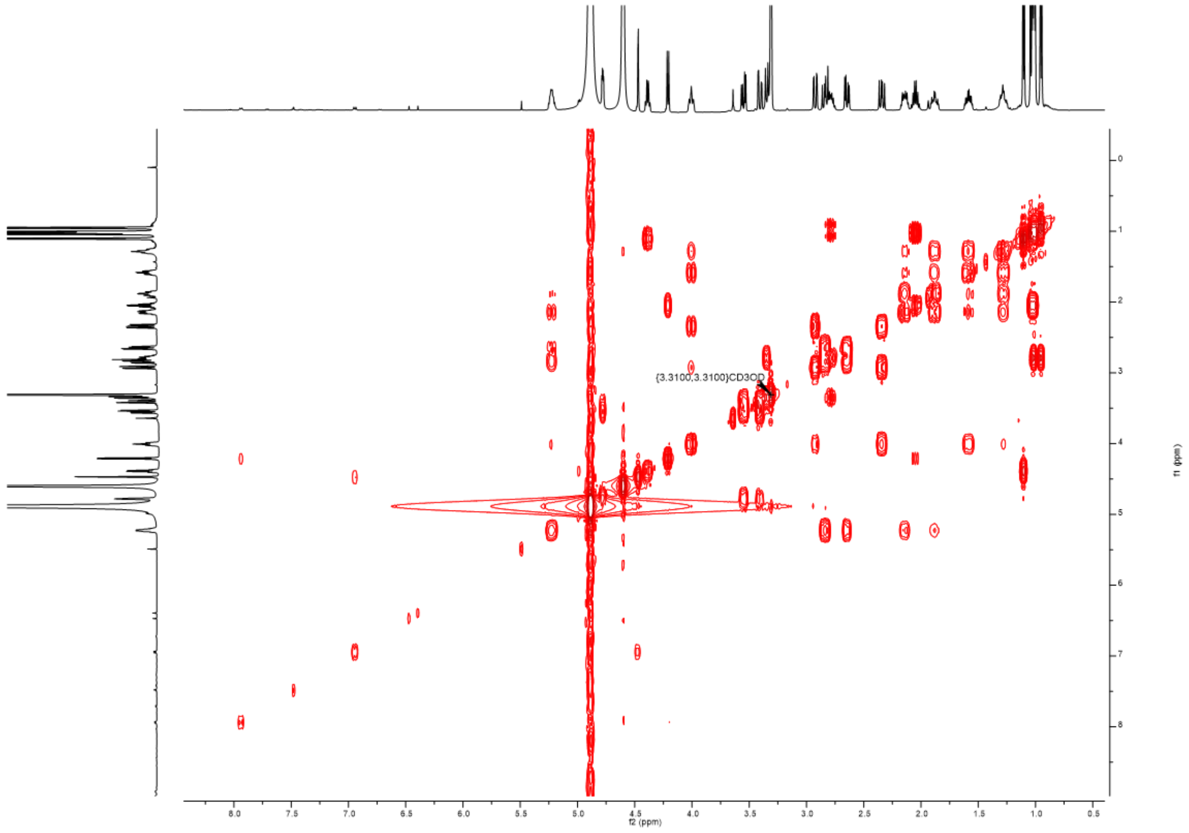


**Figure S6. COSY NMR spectrum of 6 in MeOD at 500 MHz**

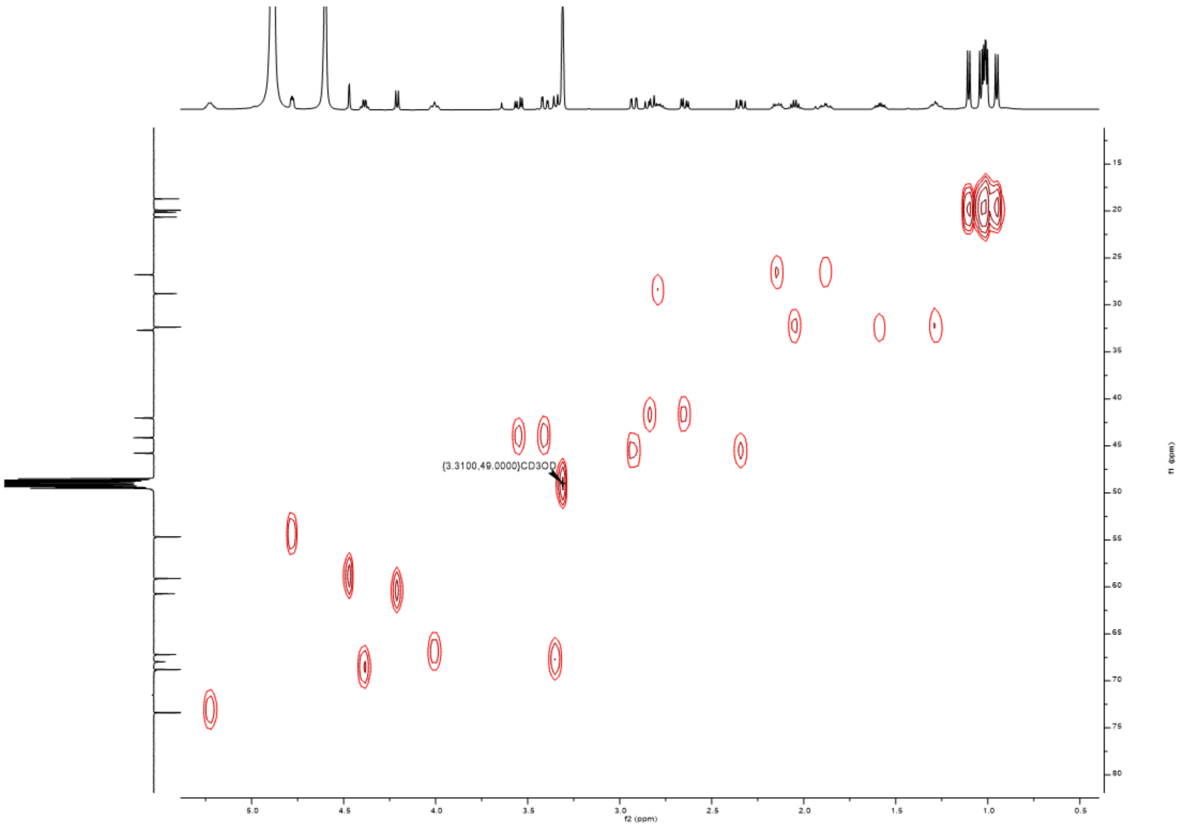


**Figure S7. HSQC NMR spectrum of 6 in MeOD at 500 MHz**

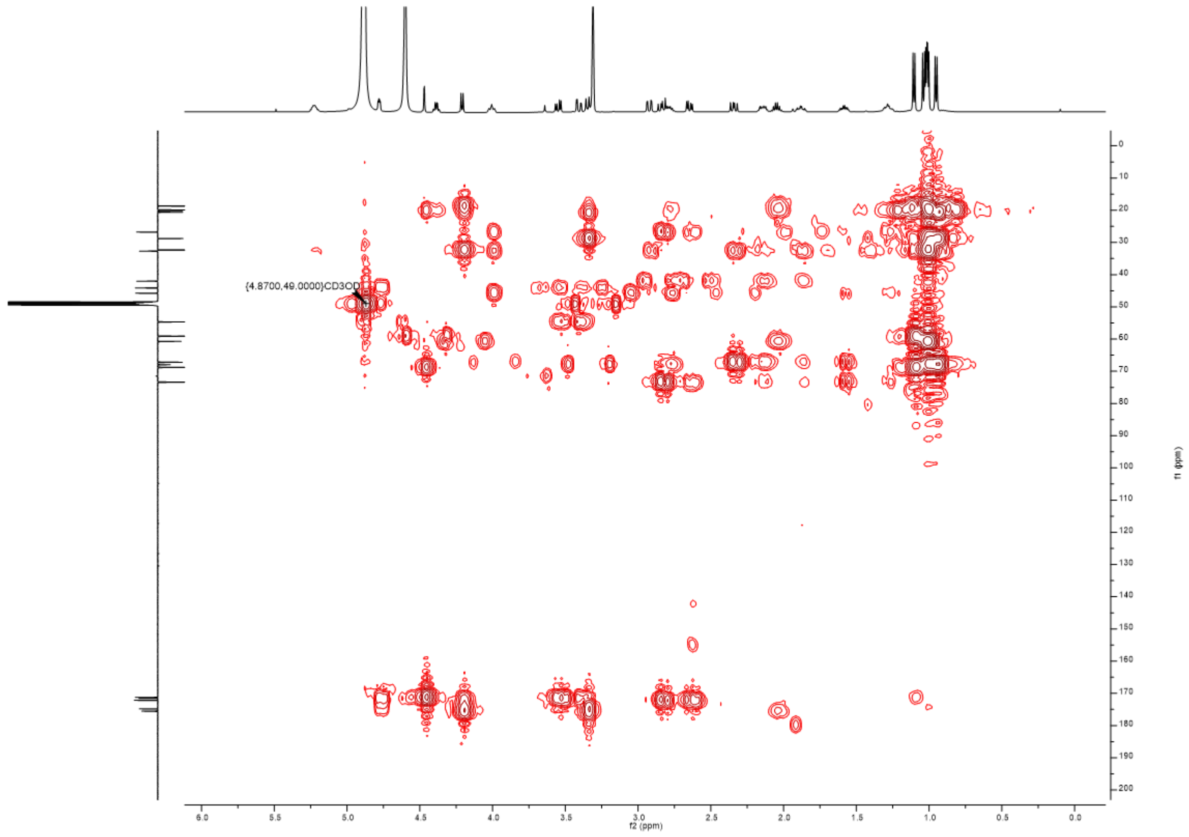


**Figure S8. HMBC NMR spectrum of 6 in MeOD at 500 MHz**

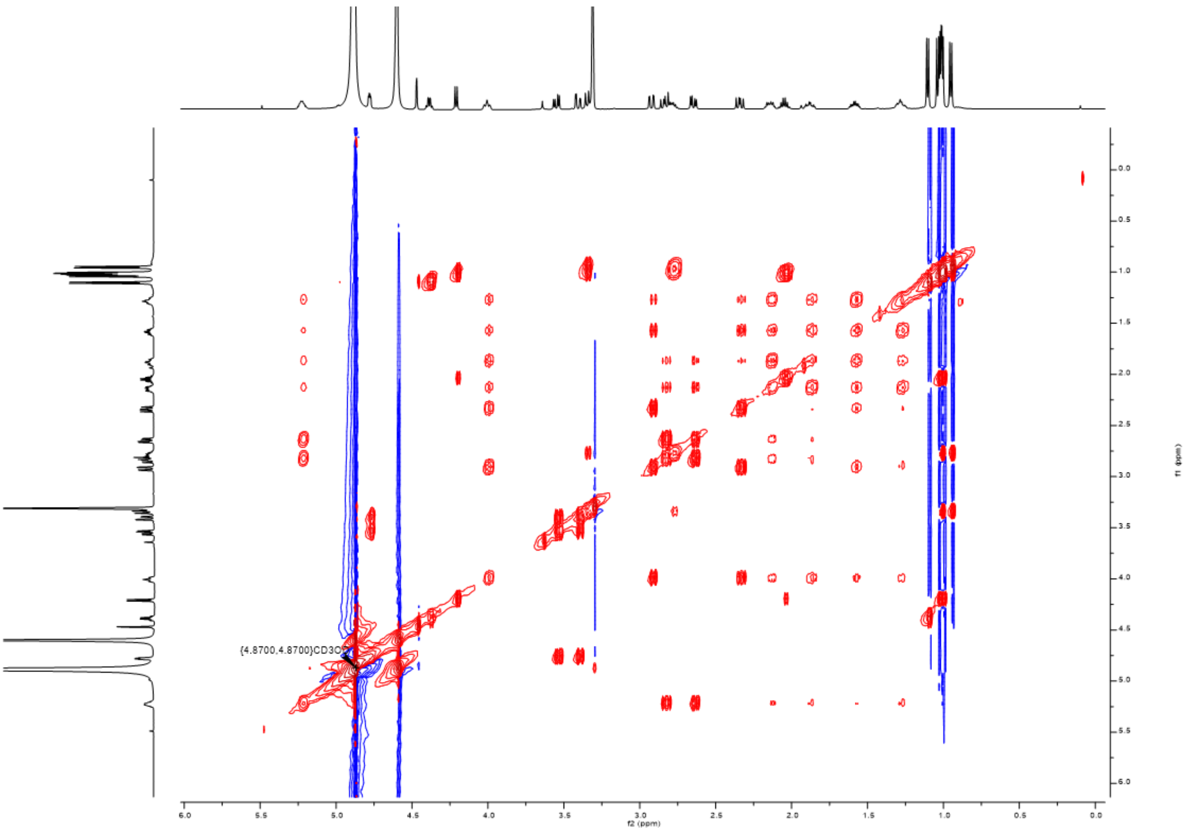

**Figure S7. NOESY NMR spectrum of 6 in MeOD at 500 MHz**

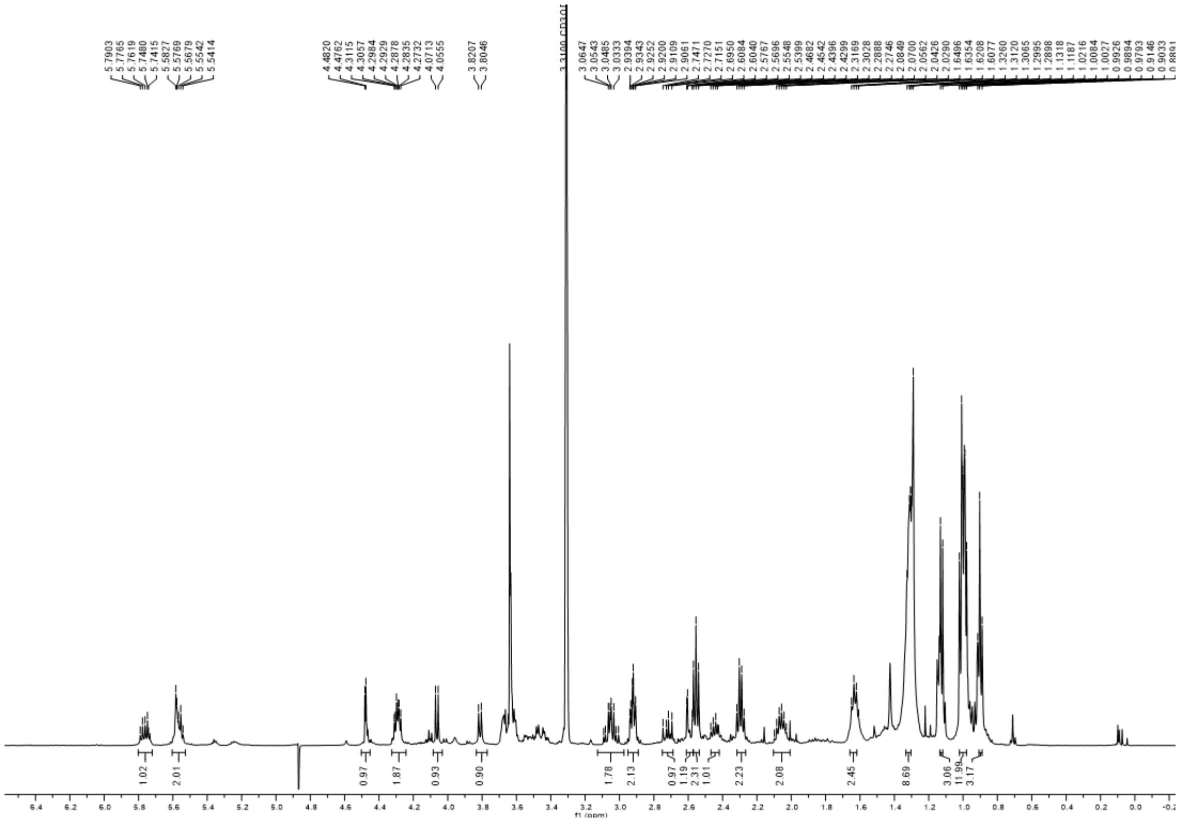

**Figure S9. $^{1}$H NMR spectrum of 7 in MeOD at 500 MHz**

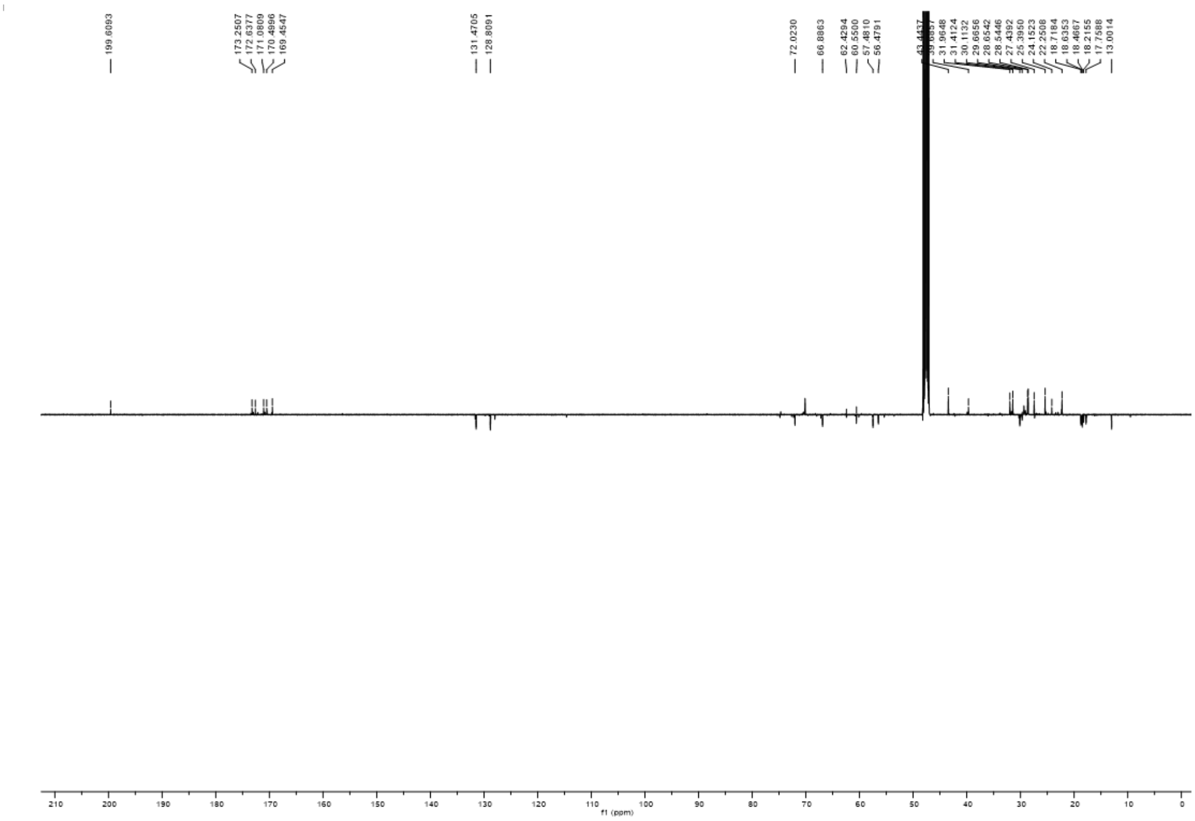

**Figure S10.** $^{13}$**C NMR spectrum of 7 in MeOD at 125 MHz**

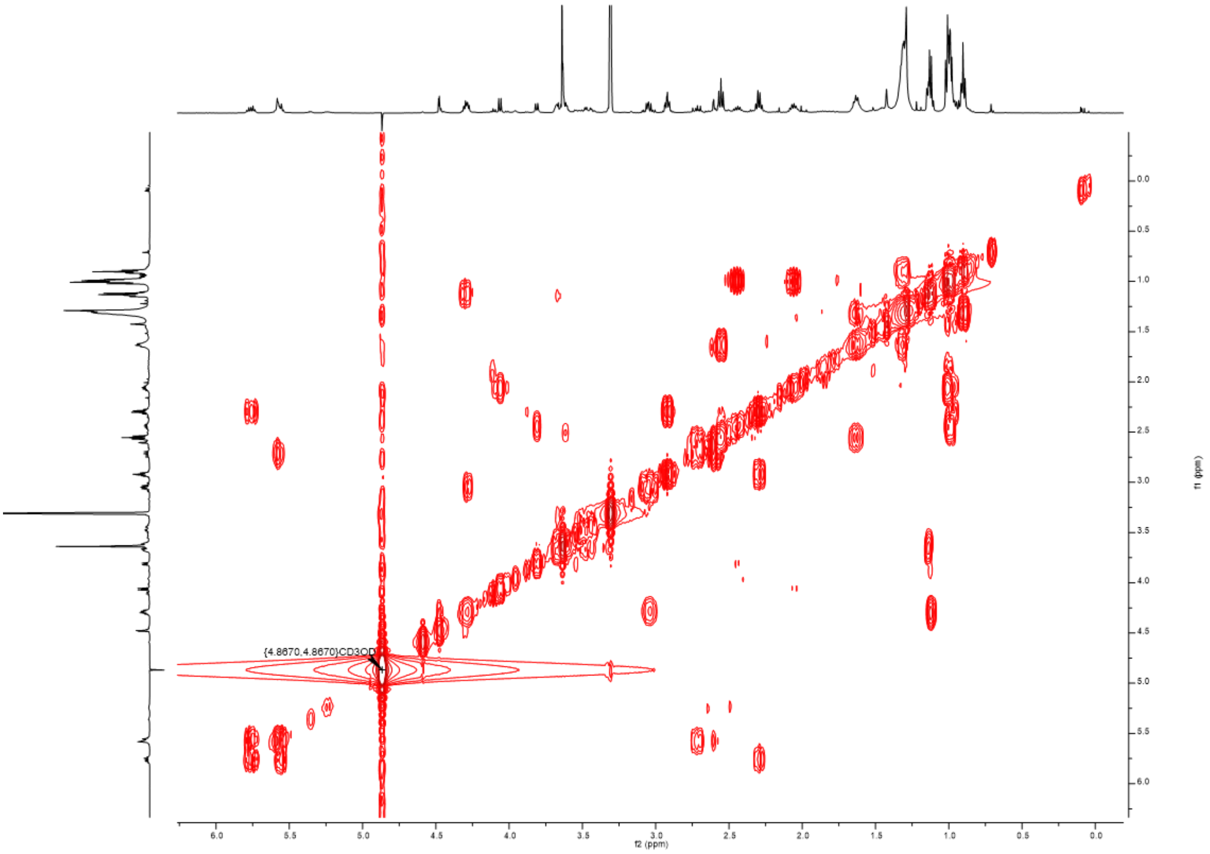

**Figure S11. COSY NMR spectrum of 7 in MeOD at 500 MHz**

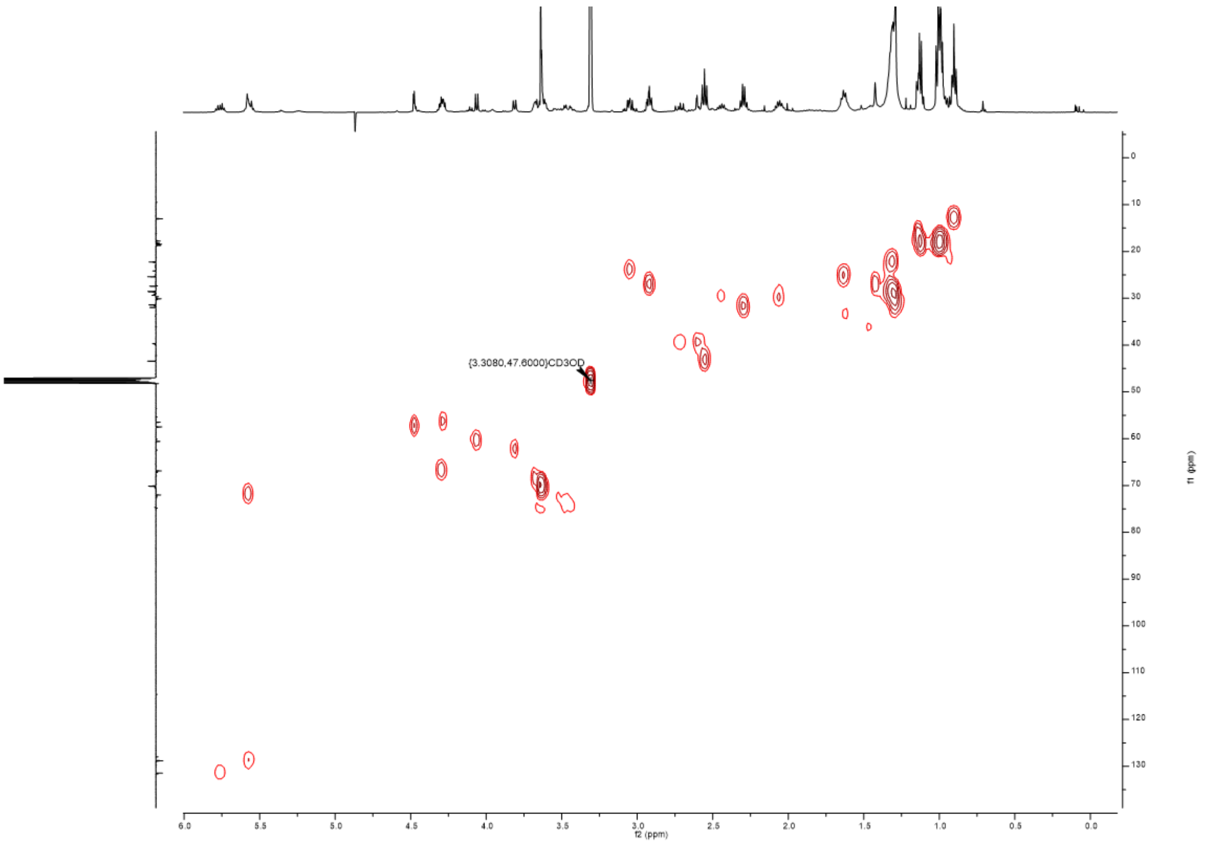


**Figure S12. HSQC NMR spectrum of 7 in MeOD at 500 MHz**

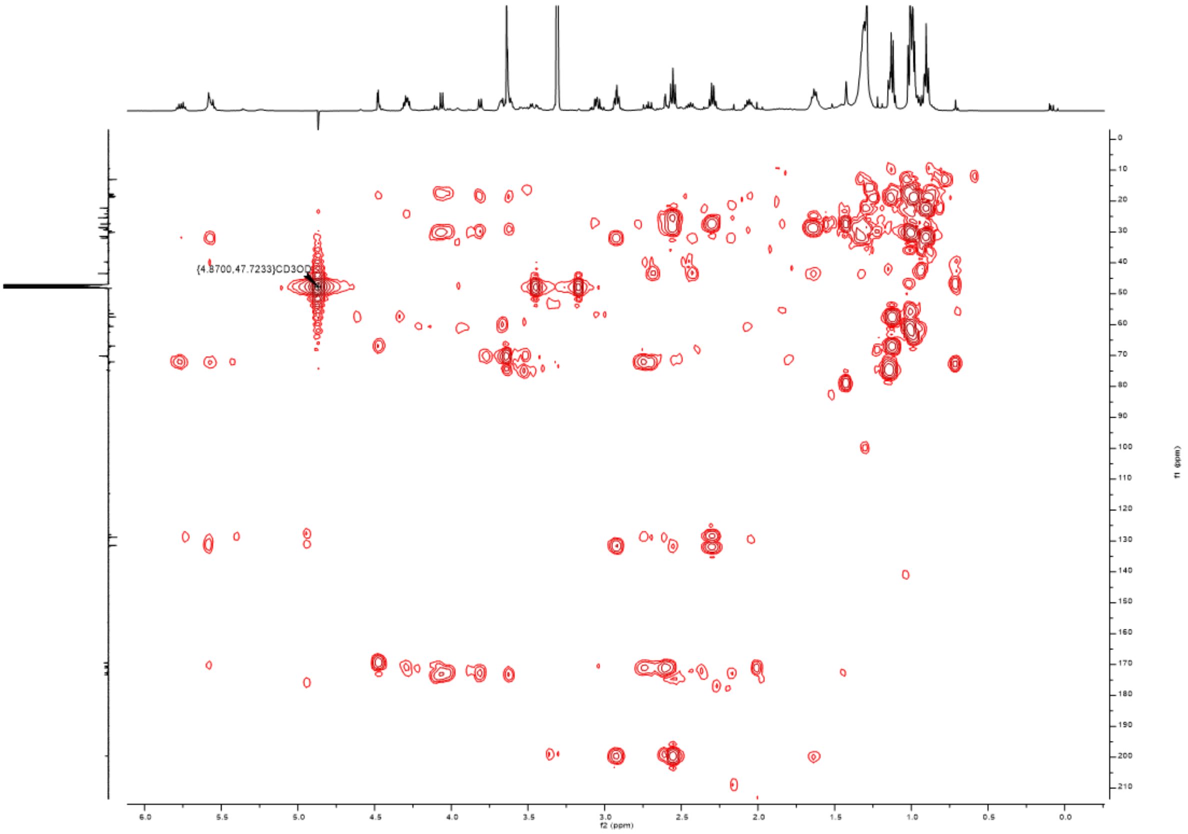


**Figure S13. HMBC NMR spectrum of 7 in MeOD at 500 MHz**

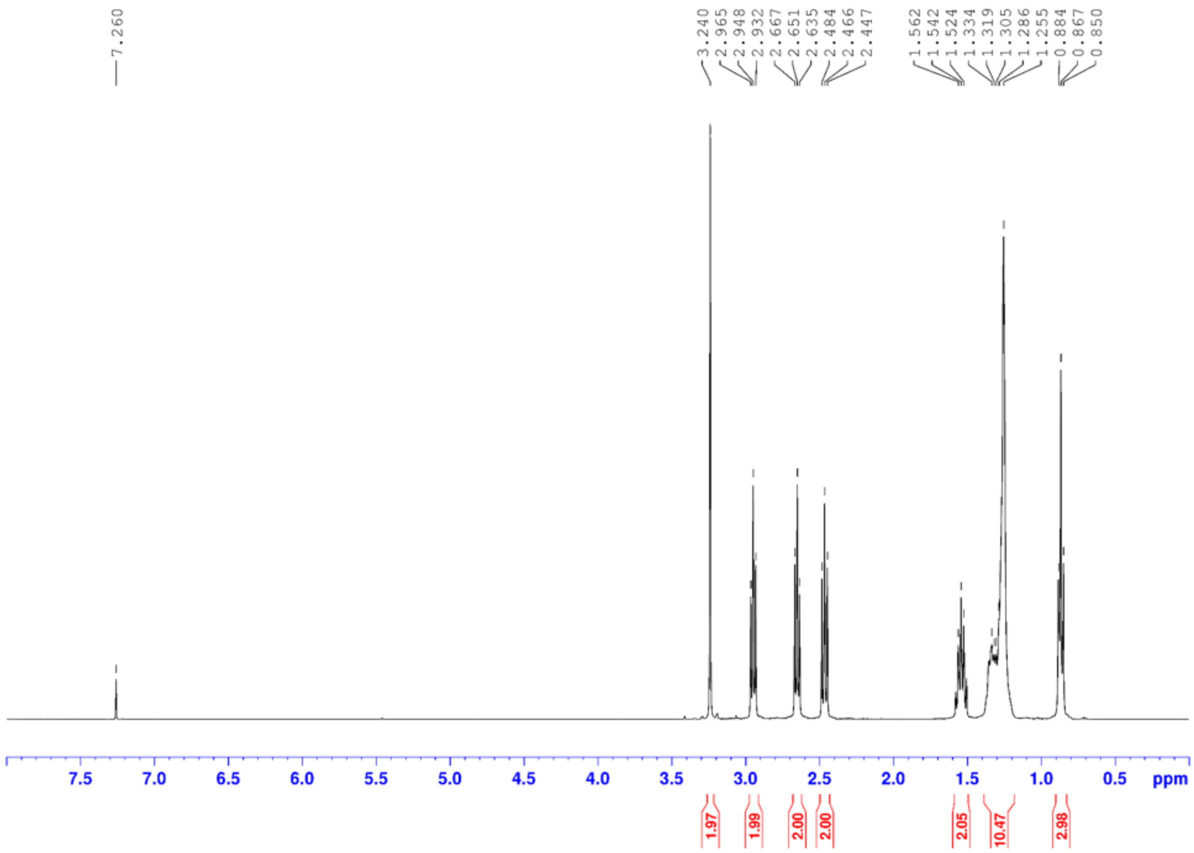


**Figure S14 $^{1}H$ NMR spectrum of 27 in $CDCl_3$ at 400MHz.**

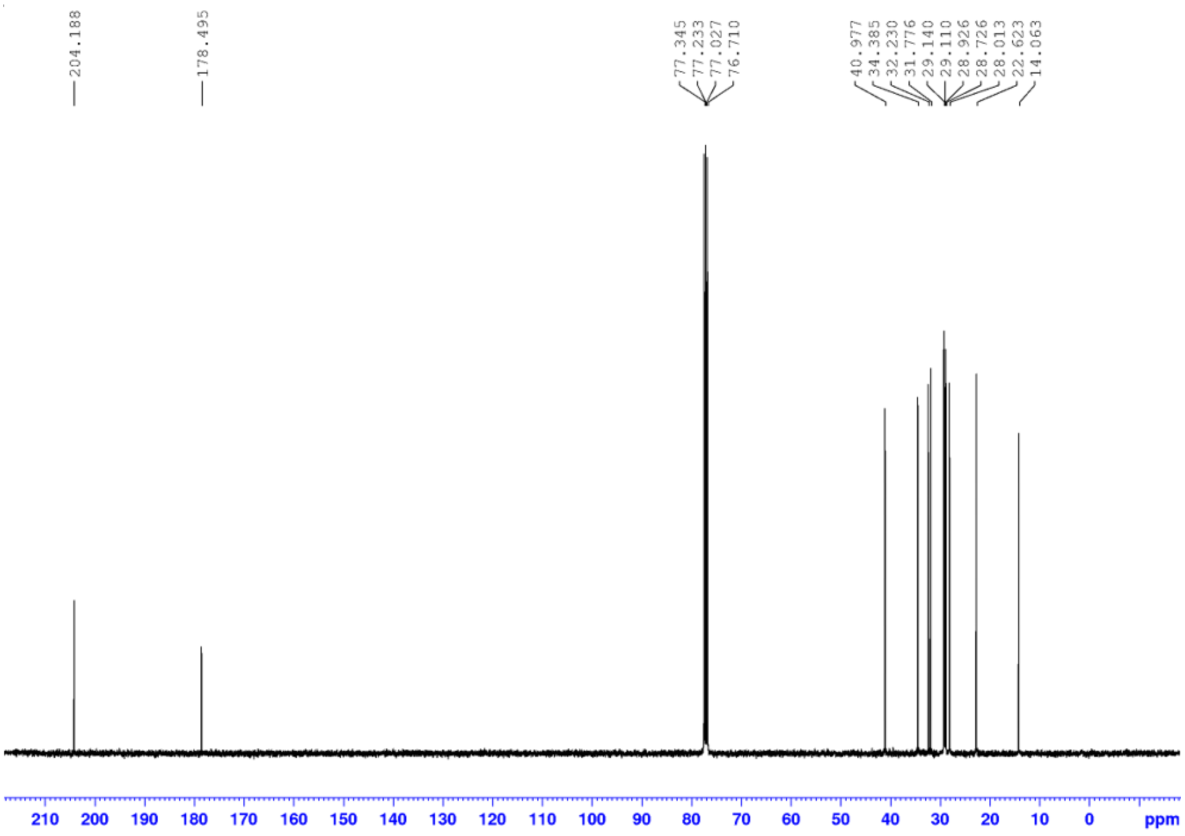


**Figure S15 $^{13}C$ NMR spectrum of 27 in $CDCl_3$ at 100MHz.**

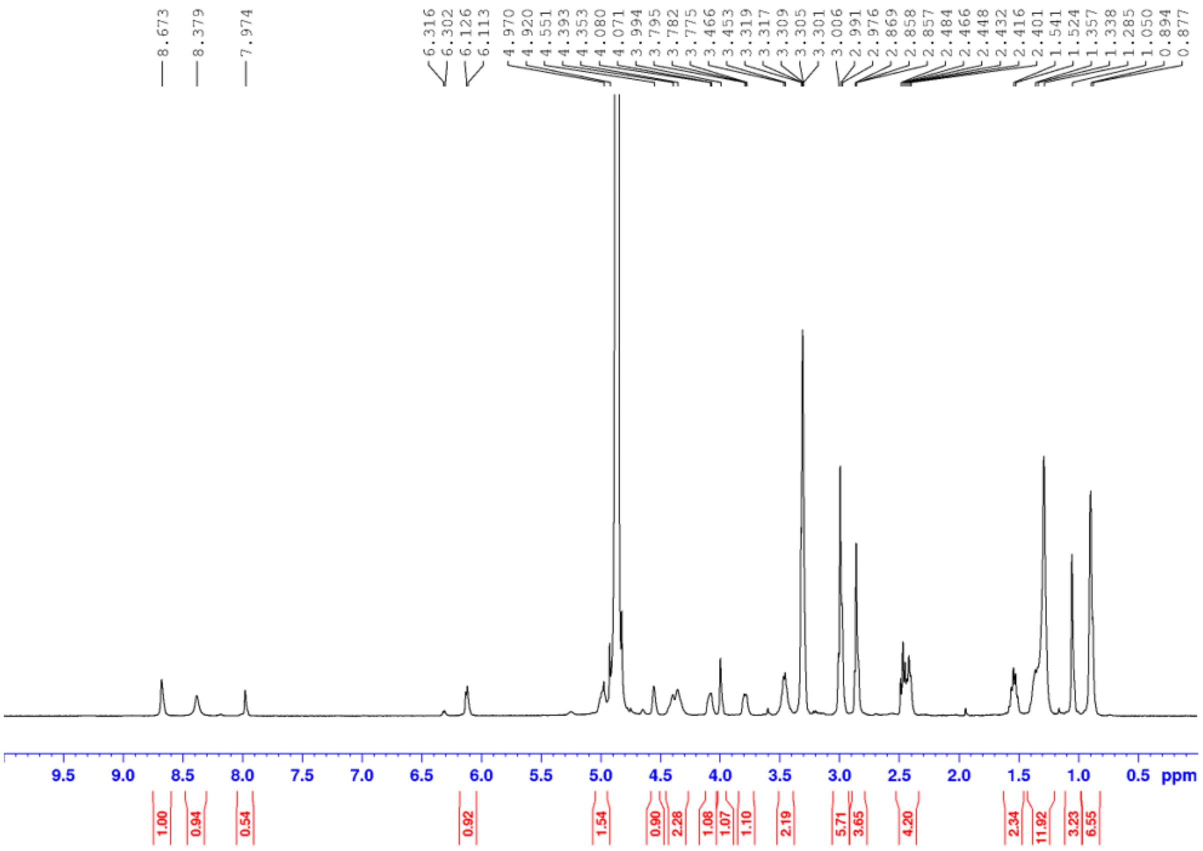

**Figure S16 $^{1}$H NMR spectrum of 15 in CD3OD at 400MHz.**

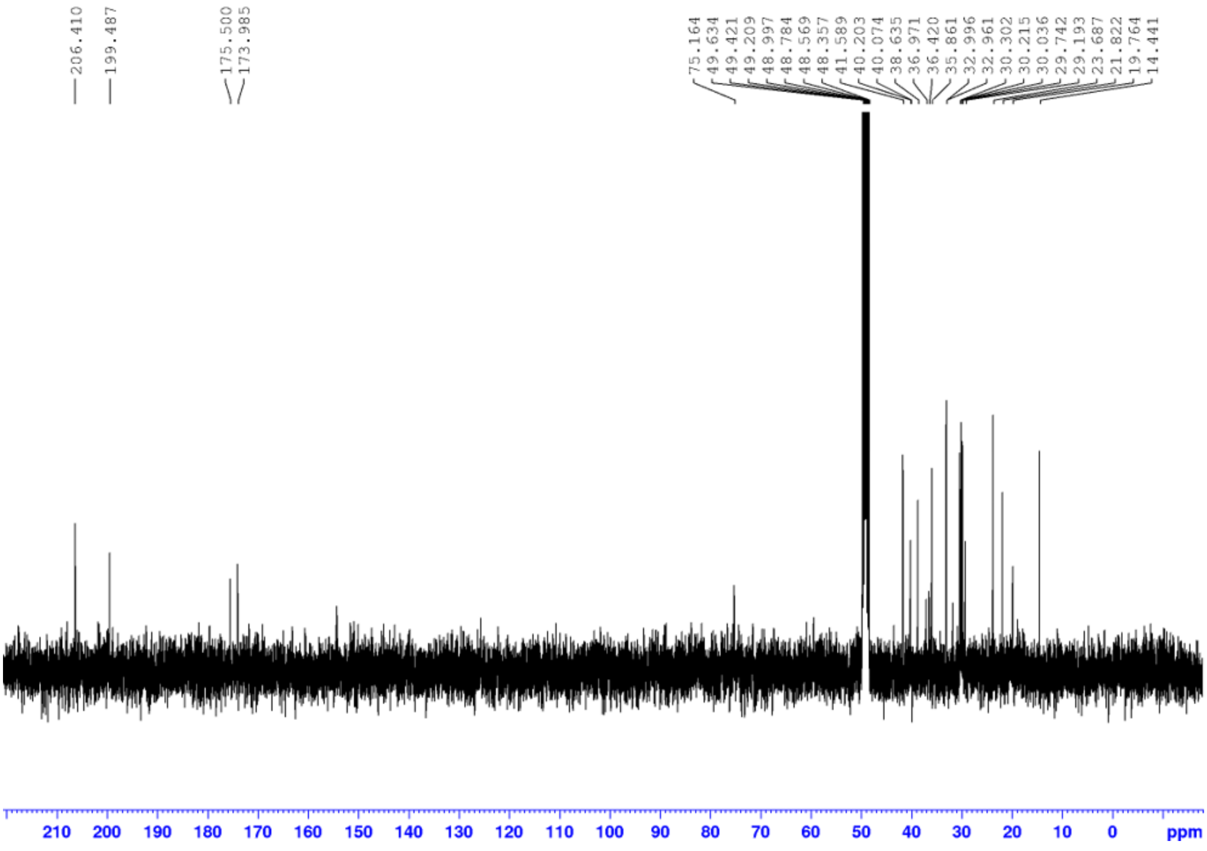

**Figure S17 $^{13}$C NMR spectrum for 15 in CD3OD at 400MHz.**

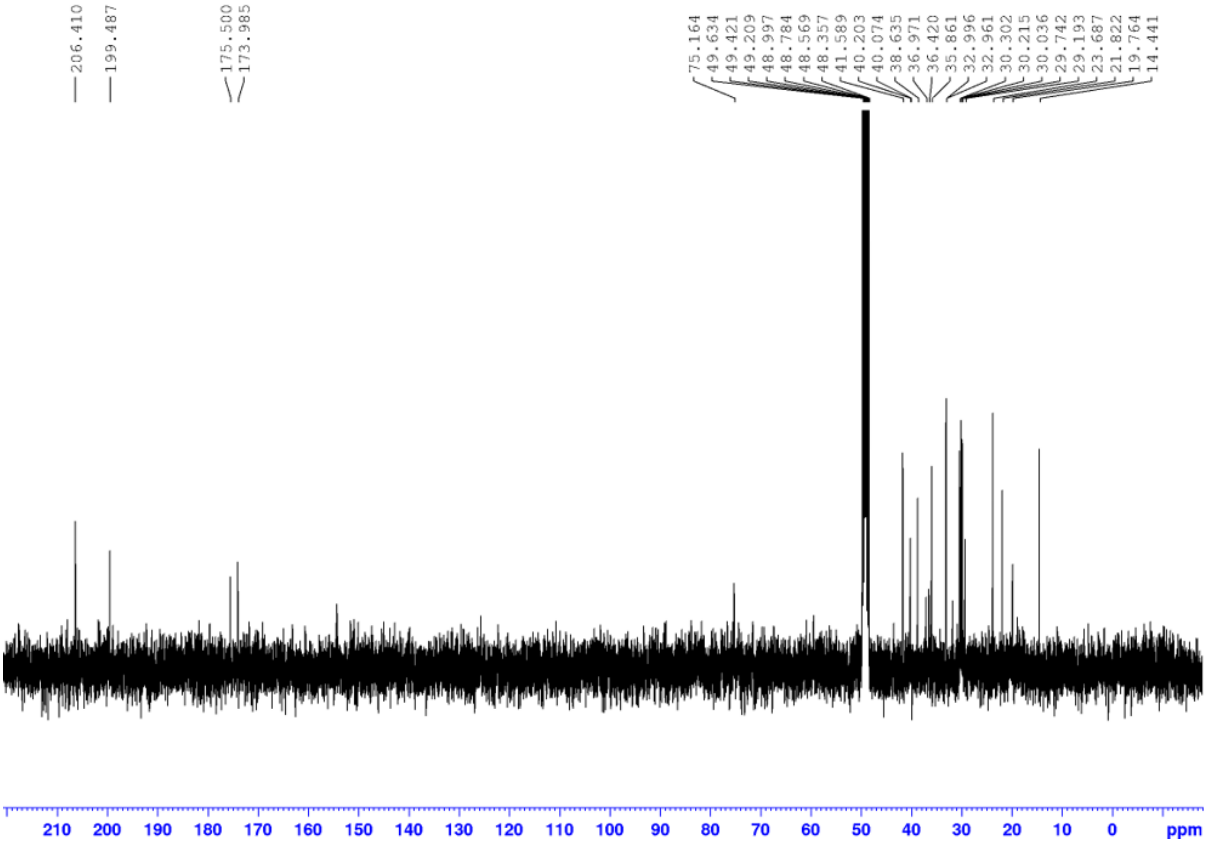


**Figure S18 $^{31}$P NMR spectrum for 15 in CD3OD at 162MHz.**

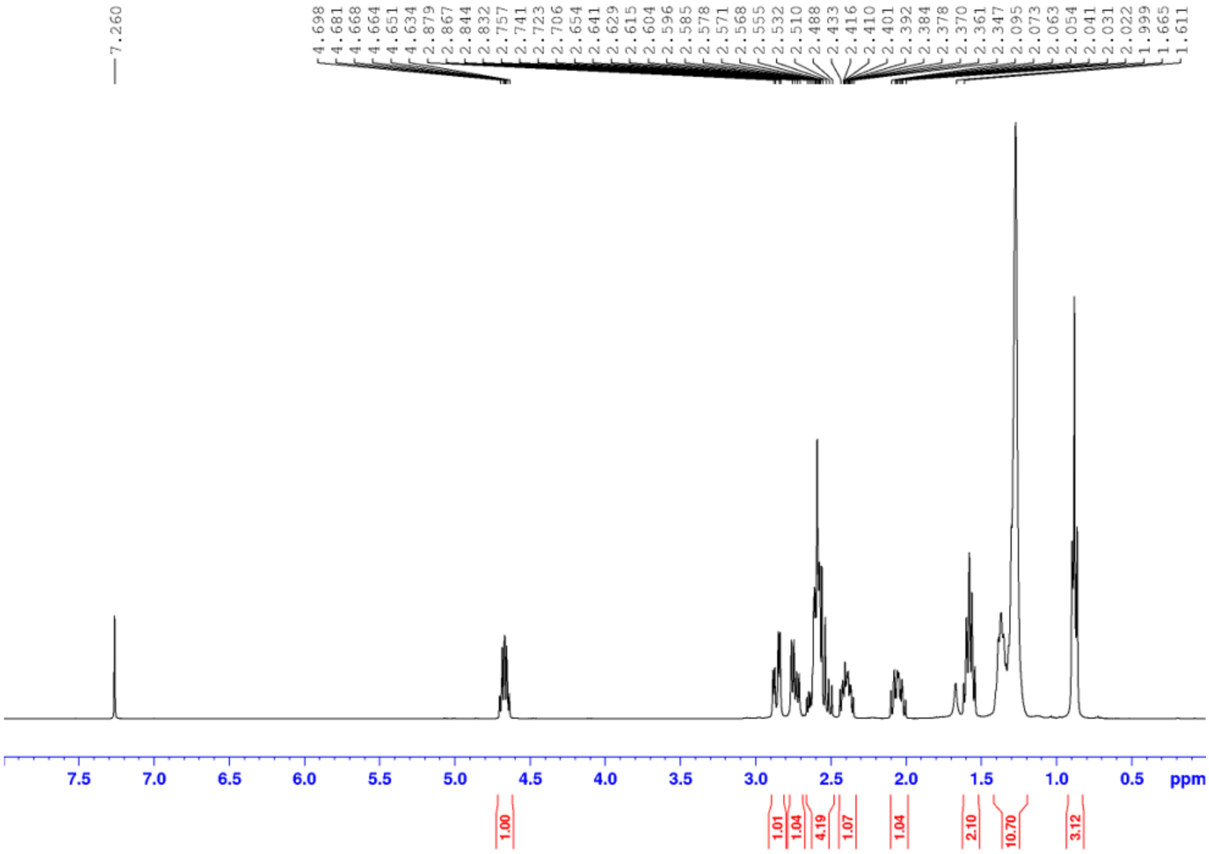


**Figure S19 $^{1}$H NMR spectrum of 22 in $CDCl_3$ at 400 MHz.**

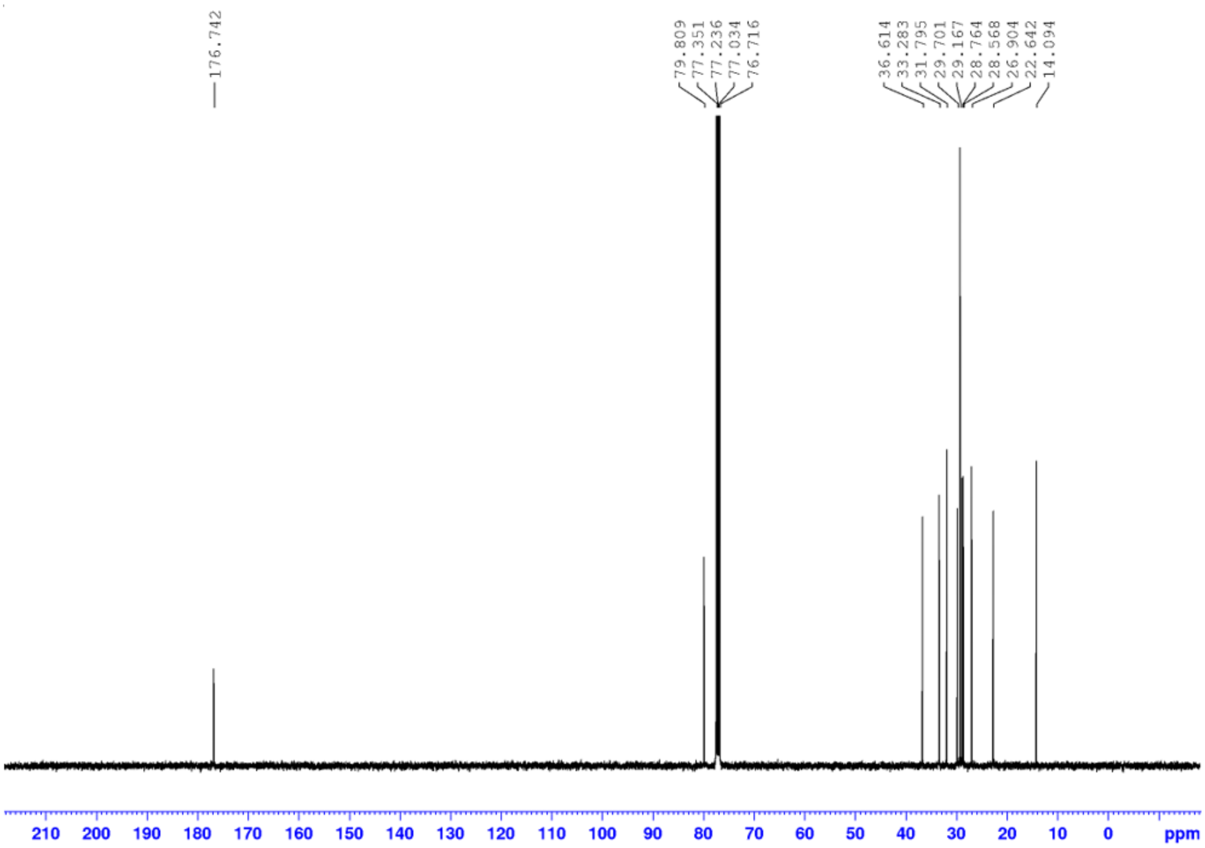


**Figure S20 $^{13}$C NMR spectrum for 22 in $CDCl_3$ at 100 MHz.**

#### References


1. Wang, C. *et al.* Thailandepsins: Bacterial products with potent histone deacetylase inhibitory activities and broad-spectrum antiproliferative activities. *J. Nat. Prod.* **74**, 2031–2038 (2011).

2. Tng, J. *et al.* Achiral derivatives of hydroxamate AR-42 potently inhibit class i HDAC enzymes and cancer cell proliferation. *J. Med. Chem.* **63**, 5956–5971 (2020).

3. Mak, J. Y. W. *et al.* HDAC7 Inhibition by Phenacetyl and Phenylbenzoyl Hydroxamates. *J. Med. Chem.* **64**, 2186–2204 (2021).

4. Furumochi, S. *et al.* Effect of carbon chain length in acyl coenzyme A on the efficiency of enzymatic transformation of okadaic acid to 7-O-acyl okadaic acid. *Bioorg. Med. Chem. Lett.* **26**, 2992–2996 (2016).
